# Repeated VU0155069 Treatment Is Associated with Region- and Pathological Stage-Specific Dendritic Spine Remodeling in Female 3xTg-AD Mice

**DOI:** 10.64898/2026.10.26.762038

**Authors:** Sanjana Mohanty, Sarah Bosworth, Karthik Ramaswamy, Chandramouli Natarajan, Balaji Krishnan

## Abstract

**Background:** Alzheimer’s disease (AD) is increasingly recognized as a disorder of synaptic integrity in which changes in dendritic spine abundance and morphology accompany progressive cognitive dysfunction. Phospholipase D1 (PLD1), a regulator of membrane trafficking and cytoskeletal signaling, is aberrantly elevated in AD-related pathological states (ADRD). Previous studies from our lab demonstrated that repeated VU0155069 treatment promotes functional resilience in female 3xTg-AD mice, including improvements in novel-object recognition, fear-conditioning performance, hippocampal LTP, later-stage LTP and LTD, selected neuropathological outcomes, and most importantly, CA1 spine integrity.

**Objective:** To determine how this previously demonstrated functional resilience is represented structurally across cortical, hippocampal, and amygdalar circuits at earlier and later stages of AD-like pathology.

**Methods:** Female 3xTg-AD mice received vehicle or VU0155069 at 1 mg/kg intraperitoneally every other day for one month beginning at 5 or 11 months of age, with Golgi-Cox analysis at 6 or 12 months, respectively. Age-matched vehicle-treated 3xTg-AD littermates served as the primary comparator for evaluating the structural response to pharmacological attenuation of disease-associated PLD1 signaling.

**Results:** Auditory and visual cortical fields showed selective changes in spine-subtype distribution and morphometry. Hippocampal cornu ammonis (CA)2, CA3, and dentate gyrus (DG) exhibited distinct patterns involving spine abundance, density, and geometry, whereas the central, lateral, and basolateral amygdalar nuclei showed nucleus- and stage-dependent remodeling. Across the multiregional dataset, recurring responses comprised spine-subtype redistribution, changes in spine abundance, and morphometric remodeling rather than uniform structural rescue.

**Conclusions:** VU0155069 engages circuit- and pathological stage-specific forms of dendritic spine remodeling across distributed recognition- and associative-memory networks.

**Graphical Abstract:** 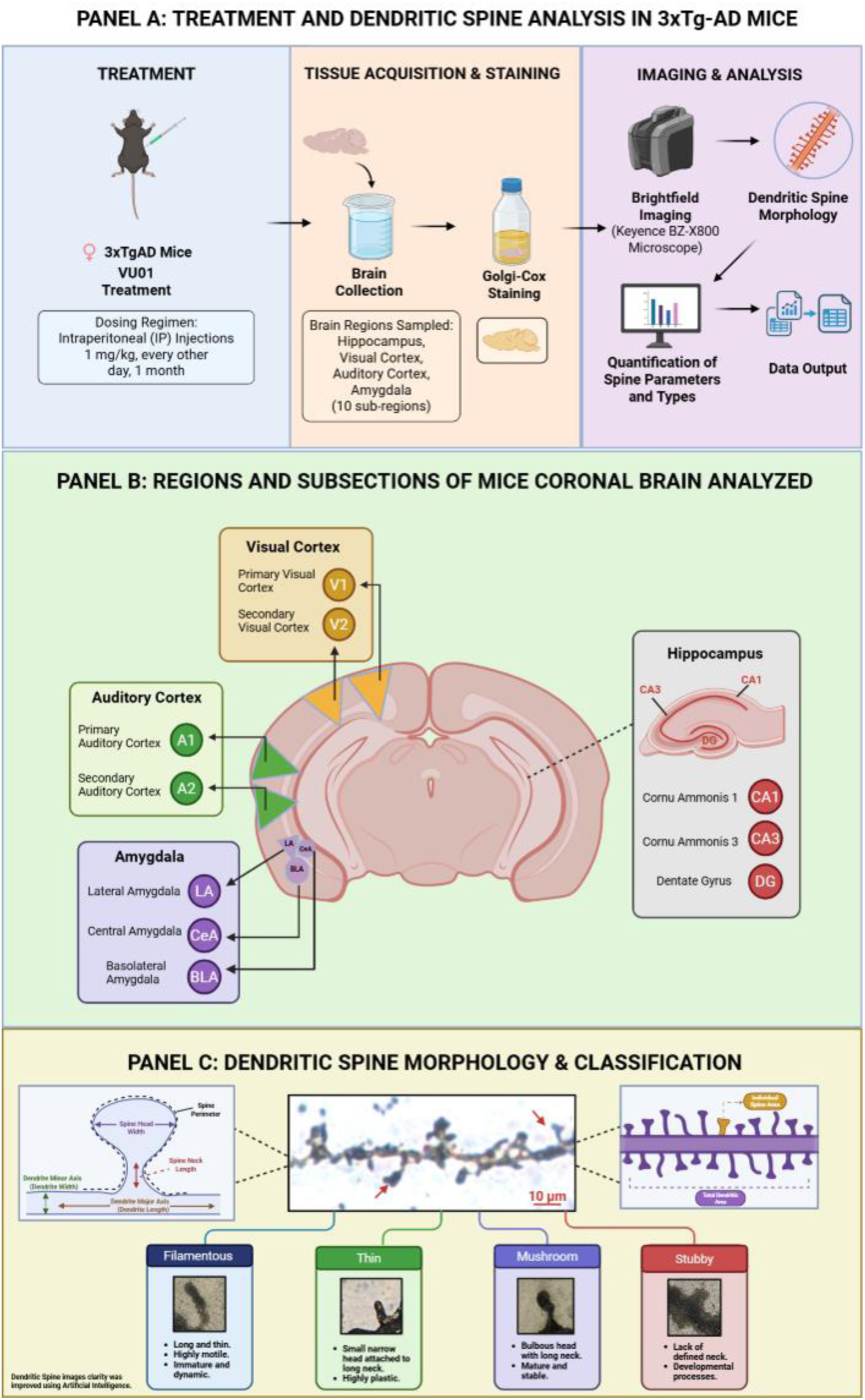

## Introduction

Alzheimer’s disease (AD), the leading cause of dementia, is increasingly understood as a synaptopathy, where widespread synaptic failure precedes and drives cognitive decline[1]. The primary structural correlates of this synaptic failure are the loss or morphological alterations of the dendritic spines (the postsynaptic sites of most excitatory synapses in the brain)[2–4]. The integrity of these spines is highly vulnerable to the toxic insults of amyloid-β (Aβ) and pathological tau [5,6]. Progressive neuronal loss following such insults, form the basis of the memory loss and neuropsychiatric symptoms of AD [7,8]. Thus, quantitative assessment of dendritic spine integrity provides a direct structural readout of synaptic dysfunction and serves as a critical endpoint for evaluating therapeutic strategies aimed at restoring synaptic health.

Phospholipase D1 (PLD1) is an inducible isoform belonging to the superfamily of phospholipases, which are lipolytic enzymes, identified for its role in generating phosphatidic acid and choline by breaking down phosphatidyl choline. Phospholipase D1 (PLD1) was identified as a key signaling element involved in normal synaptic function[9–11] and implicated in AD-related synaptotoxicity[12–15]. Under normal conditions, regulated PLD1 is essential for membrane trafficking and cytoskeletal dynamics that support synaptic plasticity[16–20]. However, in the context of AD, PLD1 activity becomes aberrantly upregulated, contributing to synaptic dysfunction and exacerbating pathology at the level of the dendritic spines[21,22]. This dual role positions PLD1 as a compelling therapeutic target whose modulation may restore synaptic stability by rebalancing structural plasticity mechanisms disrupted in AD. Accordingly, we explored the effects of repeated small molecule attenuation of PLD1 signaling as a strategy to promote synaptic resilience through preservation and/or remodeling of dendritic spine architecture.

Previously, we report that repeated administration of low dose of PLD1 inhibitor VU0155069, in 6- and 12-month-old 3xTg-AD mice, rescued both cognitive deficits in associative memory mechanism and underlying long-term potentiation (LTP) in hippocampal Schaffer Collateral (CA3->CA1) synapses[13,14]. While these studies demonstrated a clear functional and cognitive rescue, the underlying structural modifications at the circuit level remained uncharacterized. A critical next step was to determine how PLD1 attenuation remodels the specific synaptic structures underlying these functional improvements.

AD pathology does not progress uniformly throughout the brain; rather, different neural circuits exhibit distinct vulnerabilities as the disease progresses due to their highly specialized cellular compositions. Dendritic spine integrity can be an effective quantitative tool to assess the degree of disease progression[23–27]. The 3xTg-AD model recapitulates this heterogeneity, with early and severe pathology in the hippocampus and basolateral amygdala, followed by a later-stage progression into sensory cortices[28–30]. Together, these studies established a longitudinal therapeutic framework in which VU0155069 promoted functional resilience across recognition and associative-memory outcomes, hippocampal synaptic plasticity, selected neuropathological measures, and CA1 spine integrity at earlier and later stages of 3xTg-AD pathology. However, these earlier findings raised a more anatomically resolved question: does the structural response extend uniformly across the brain, or does it depend on the identity and pathological context of the region examined?

Therefore, the present study was designed to determine how this established therapeutic response is represented structurally across the broader cortical, hippocampal, and amygdalar networks that support the acquisition, integration, and expression of these behaviors. Novel-object recognition recruits visual and perirhinal-hippocampal processing, whereas fear conditioning engages auditory and visual sensory input together with hippocampal contextual processing and amygdalar associative circuitry. A multiregional spine analysis therefore provides the anatomical resolution needed to connect previously demonstrated functional resilience with the distributed circuits supporting those outcomes. This study aimed to provide the first comprehensive, circuit-level morphological analysis of the effects of PLD1 attenuation across ten distinct brain regions at both an early (6-months) and advanced (12-months) stage of pathology in the 3xTg-AD model. We selected regions that collectively span sensory processing, hippocampal memory computation, and amygdala-dependent associative function. Within each domain, paired fields or subregions were examined to determine whether anatomically related structures exhibit convergent or divergent responses to VU0155069. More specifically, we hypothesized that PLD1 inhibition would engage distinct structural response programs, including spine subtype redistribution, changes in spine abundance, and alterations in dendritic geometry, depending on circuit identity and disease stage.

The auditory cortex, which includes the primary (A1) and secondary (A2) regions, is crucial for processing acoustic information, with A1 handling basic features like frequency and A2 managing more complex auditory integration[31–33]. In AD, central auditory processing deficits are a salient and early feature, correlating with established AD biomarkers and often preceding overt cognitive decline[34,35]. The primary auditory cortex (A1), in particular, is vulnerable to AD-related pathology, including intracellular Aβ_42_ in layer 5 pyramidal neurons as demonstrated in post-natal 5xFAD mice[36]. The 3xTg-AD mouse model recapitulates this progressive pathology[28–30]. At 6-months, cortical neurons (including A1 and A2) already exhibit intraneuronal deposition of Aβ and accumulation of phosphorylated tau. By 12 months, this cortical pathology advances to include widespread extracellular Aβ plaques and more intense, aggregated tau, mirroring a more advanced disease state. This temporal progression provides an opportunity to evaluate whether PLD1 attenuation can differentially preserve or remodel dendritic spine structure across early and late stages of cortical pathology.

The hippocampal formation is essential for episodic memory, with its subfields (the dentate gyrus (DG), cornu ammonis 2 (CA2) and 3 (CA3) performing distinct computations. DG executes pattern separation while CA3 subfield performs pattern completion via its recurrent network, and the CA2 region acts as a critical hub for social memory and network modulation[37,38]. This circuit is a primary target of AD pathology. In the 3xTg-AD mouse model, the hippocampus is the first region to develop amyloid plaques, with intraneuronal Aβ and phosphorylated tau evident in the DG, CA2, and CA3 by 6-months of age[39,40]. This pathology intensifies by 12-months with robust extracellular plaque deposition and tau aggregation throughout the hippocampal subfields, accompanied by progressive dendritic spine loss[39,41]. Given the distinct functional roles of these subfields and their shared vulnerability in this model, we analyzed the DG, CA2, and CA3 to determine if VU0155069 treatment could exert differential effects within the hippocampal circuit. This approach enables direct assessment of subfield-specific therapeutic responsiveness within a single interconnected memory circuit.

The visual system, with the primary (V1) and secondary (V2) cortices, processes visual information hierarchically, from basic features in V1 to more complex attributes in V2[42,43]. Visual processing deficits are a well-documented symptom of AD, correlating with pathological changes in posterior cortical and occipital regions[44]. The 3xTg-AD mouse model reflects an age-dependent progression of this pathology wherein cortical regions including the visual cortex are largely devoid of extracellular plaques at 6-months but exhibit significant amyloid and tau pathology by 12-months, accompanied by progressive dendritic spine loss[27,39]. This staged progression enables evaluation of how PLD1 attenuation influences synaptic structure in circuits transitioning from relatively preserved to pathologically engaged states.

The amygdala is critical for emotional processing, with its nuclei showing differential vulnerability in AD. The basolateral amygdala (BLA) is essential for emotional learning, the lateral amygdala (LA) for sensory integration, and the central amygdala (CeA) for autonomic output[45]. In the 3xTg-AD model, this vulnerability is recapitulated wherein the BLA exhibits the earliest and most severe pathology, with significant dendritic spine loss evident by 6-months that progresses by 12-months[40,46]. The LA is moderately affected, while the CeA is comparatively spared from major pathological changes at these time points[47,48]. This hierarchy of vulnerability provides a unique opportunity to determine whether PLD1 inhibition elicits uniform or nucleus-specific structural remodeling across associative memory circuits.

Thus, the auditory and visual cortices, hippocampal subfields, and amygdalar nuclei contribute to distinct sensory, mnemonic, and associative functions and may therefore exhibit different structural responses to intervention.

The primary and secondary auditory and visual cortices were included to resolve cortical responses at finer anatomical scale than prior broad regional physiology measurements. CA2, CA3, and the dentate gyrus were included because they perform distinct hippocampal computations, while the central, lateral, and basolateral amygdala were examined as anatomically and functionally differentiated nuclei. None of the regional measurements, statistical analyses, or figures presented here was included in our prior publication assessing CA1 dendritic spine morphology. The primary pharmacological comparison was between age-matched vehicle-and VU0155069-treated 3xTg-AD littermates. This design directly tests whether attenuation of aberrantly elevated PLD1 signaling modifies synaptic structure within the affected population.

Age-matched wild-type animals would provide complementary information regarding the physiological range of each regional endpoint, but they address a different question from the disease-state therapeutic comparison pursued here.

We hypothesized that repeated VU0155069 treatment would alter dendritic spine architecture in a region-and pathological stage-dependent manner. The primary analyses compared vehicle-and VU0155069-treated mice within each age, while the multiregional dataset was used exploratorily to identify recurring patterns of spine-subtype redistribution, altered spine abundance, and morphometric remodeling.

## Methods

### Animals, colony maintenance, and ethical approval

Female triple-transgenic Alzheimer’s disease mice, hereafter referred to as 3xTg-AD mice, were obtained from the established breeding colony maintained through the Mitchell Center for Neurodegenerative Diseases at The University of Texas Medical Branch at Galveston. The colony was originally established using 3xTg-AD mice obtained from The Jackson Laboratory. Breeding and colony maintenance were overseen by Batbayar Tumurbaatar and supported by the Mitchell Center for Neurodegenerative Diseases. The genetic integrity and relative copy-number consistency of the 3xTg-AD transgenic inserts were monitored by batch-level RT-PCR. Every breeding batch was evaluated, although individual copy-number analysis was not necessarily repeated for every study animal. All animals included in the present study were 3xTg-AD mice derived from breeding batches that satisfied the established colony-quality criteria.

Only females were studied because AD disproportionately affects women and female 3xTg-AD mice exhibit a deeply penetrant pathological phenotype[28,49–51]. All four age-by-treatment groups contained n = 5 animals. Thus, the study is not designed to compare biological sexes, and the present study cannot determine whether the dendritic spine response to VU0155069 generalizes to male 3xTg-AD mice, although our earlier published studies show functional improvement in both male and female 3xTg-AD cohort (separate from the current cohort of just female mice) synaptic function measured using electrophysiology, and cognitive function assessed using two different behaviors with both the early[13] and late[14] pharmacological interventions. Animals were group-housed five per cage at approximately 25°C and 50% relative humidity on a 12-hour dark cycle, with standard chow and water ad libitum and no additional enrichment. Vehicle-and VU0155069-treated littermates were housed together, and randomized treatment assignments were distributed across cages. Injections were administered during the light phase at approximately 2:00 PM. All procedures were approved by the UTMB Institutional Animal Care and Use Committee under protocol 1710061, active from October 2018 through both cohorts. Procedures followed the NIH Guide for the Care and Use of Laboratory Animals and applicable ARRIVE reporting principles. Predefined humane endpoints were in place; no animal reached an endpoint or required premature euthanasia.

Littermates were assigned by random-number generation to vehicle or VU0155069 treatment, with body weight considered during group balancing. The 6-and 12-month cohorts comprised distinct animals and were treated separately. 20 animals were utilized, divided equally into two cohorts for 6-months and 12-months (n = 10 each). Within each cohort, animals were randomized into the two treatment groups (n = 5 per group). Treatment began at 5 or 11 months and ended at 6 or 12 months, respectively.

VU0155069 (Tocris Bioscience, catalog 3575; purity >=98%) was prepared as a 20-mg/mL stock in anhydrous ethanol, aliquoted at -20°C, thawed once, and used or discarded. On each injection day, a fresh 1-mg/mL formulation was prepared in 5% ethanol, 10% Tween-80, and sterile 0.9% saline, with vortexing and sonication until clear. Vehicle contained the identical formulation without VU0155069. Animals were weighed before each injection and received 1 mg/kg intraperitoneally at 1 mL/kg every other day for one month (15 injections). Successive injections alternated abdominal sides. Body weight, survival, injection sites, appearance, activity, and signs of distress were recorded; no treatment-related exclusion occurred.

The present morphometric analysis used tissue from previously established treatment cohorts. The analysis used tissue from established cohorts containing five animals per age-by-treatment group. Cohort size reflected animal and tissue availability, prior laboratory experience with Golgi–Cox morphometry, and planned sampling of multiple eligible dendritic segments from each animal. Based on previously published approaches[13,14], each animal contributed up to 15 eligible dendritic segments per region, providing extensive within-animal morphometric sampling without increasing the number of independent biological replicates.

### Tissue Processing and Golgi-Cox Staining

Immediately after the final injection, animals were deeply anesthetized with isoflurane and transcardially perfused with artificial cerebrospinal fluid according to previously published protocols. Collections occurred at approximately the same time of day in intermixed treatment order. Brains were removed, rinsed, and randomized by hemisphere. One hemisphere underwent Golgi-Cox processing; the contralateral hemisphere was reserved for separate studies not reported here. Archived 6-and 12-month slides were generated in 2018 and 2019, respectively.

### Golgi-Cox impregnation, sectioning, and storage

Hemispheres were processed using the FD Rapid GolgiStain Kit (PK401; FD NeuroTechnologies). Samples were immersed in Solutions A and B in darkness for 15 days, with solution replacement after 24 hours, and then transferred to Solution C, which was replaced after 24 hours. FD NeuroTechnologies cut 30-micrometer coronal sections, mounted three sequential sections per gelatin-coated slide (Cat. No. PO101, FD NeuroTechnologies), developed, dehydrated, cleared, and coverslipped the specimens. Vehicle and VU0155069 samples were processed together within each age-specific batch using the same reagent lot. Slides were stored at room temperature in darkness until imaging.

The 6-and 12-month cohorts were processed in separate age-specific Golgi–Cox batches. Within each age batch, vehicle-and VU0155069-treated brains were processed together using reagents from the same lot, and treatment conditions were intermixed during sectioning.

Slides from the 6-month cohort were generated in 2018, and slides from the 12-month cohort were generated in 2019. Slides were stored at room temperature in darkness within slide boxes under comparable ambient-humidity conditions until the present regional imaging study. Before image acquisition, slides were inspected for fading, precipitate, tissue detachment, coverslip damage, and adequacy of Golgi–Cox impregnation. Individual slides or sections that did not provide adequate visualization were not imaged. When an individual slide or section was unsuitable, another eligible sequential section from the same animal and anatomical region was used. Every animal ultimately contributed usable measurements from all ten regions; no animal was replaced or excluded.

### Regional localization, microscopy, and sampling

Ten regions were examined: primary auditory cortex (A1), secondary auditory cortex (A2), primary visual cortex (V1), secondary visual cortex (V2), hippocampal CA2, hippocampal CA3, dentate gyrus (DG), central amygdala (CeA), lateral amygdala (LA), and basolateral amygdala (BLA). Regions were identified using cytoarchitecture and anatomical landmarks referenced to the Paxinos and Franklin mouse brain atlas[52].

For each animal and region, three sequential coronal sections were sampled. Five nonoverlapping fields were evaluated per section, with one eligible 10-micrometer spiny dendritic segment selected per field, yielding up to 15 segments per animal per region. Sampling was not restricted by neuronal class, cortical layer, apical/basal compartment, or proximal/distal location.

Sections were surveyed at 20x for regional localization and evaluated at 100x for segment eligibility. Eligible dendrites were fully impregnated, isolated, unobscured, longer than 30 micrometers, and approximately 1 micrometer in shaft diameter. When several dendrites met criteria, the first encountered clockwise was selected; the clearest uninterrupted 10-micrometer portion was then analyzed. Selection was not based on spine abundance. Sampling advanced in one direction across each slide to minimize repeated sampling of the same parent neuron.

Images were acquired through 2024 using the same Keyence BZ-X800E microscope, 100x oil objective, 3x optical zoom, and 0.3-micrometer z-step. Full-focus TIFF reconstructions were produced with Keyence software and analyzed in ImageJ after spatial-calibration verification. Illumination and exposure were adjusted for impregnation variability, while acquisition geometry and calibration were standardized. Vehicle and VU0155069 specimens were coded and imaged in intermixed order.

Microscope configuration, objective magnification, optical zoom, z-step interval, and full-focus reconstruction procedures were maintained consistently. Illumination and exposure were adjusted when necessary to accommodate variation in Golgi-Cox impregnation and image contrast and were not used as quantitative outcomes. Lengths were recorded in micrometers, and areas were recorded in square micrometers.

### Morphometric measurements and spine classification

Eligible dendrites were required to be adequately Golgi-Cox impregnated; isolated from adjacent neuronal processes; unobscured by precipitate or staining debris; longer than 30 µm; approximately 1 µm in shaft diameter; spiny; and suitable for selection of a completely visualized, uninterrupted 10-µm segment.

When more than one dendrite satisfied these criteria within a field, the first eligible dendrite encountered during a standardized clockwise survey was selected. From that dendrite, the clearest uninterrupted and fully visualized 10-µm portion was chosen for analysis. Analysts were not permitted to select or reject a segment based on apparent spine abundance or subtype composition. Only one 10-µm segment was selected from each field. Sampling proceeded systematically across each slide in one direction to minimize repeated imaging of branches that might arise from the same parent neuron. Only segments contained entirely within the image field were analyzed. No spine touching or crossing a segment or image boundary was counted.

Because the sampled segments were not traced to their parent somata, repeated sampling of branches from the same neuron cannot be excluded with absolute certainty. The one-direction slide survey and one-segment-per-field rule were used to minimize this possibility.

Spines were assigned to mutually exclusive categories using a common written protocol[53]. All fully visualized dendritic protrusions arising within each standardized 10-µm segment were counted. Spine abundance was reported as the number of spines within the standardized segment, and spine density was expressed as the number of spines per 10 µm of dendrite.

Each protrusion was assigned to one mutually exclusive morphological category using a common written classification protocol. Classification was based primarily on visual morphology, with measured dimensions consulted when a profile was ambiguous. Detailed criteria for dendritic spine classification and the exact parameters measured throughout the study are illustrated schematically in the graphical abstract. Mushroom spines displayed a prominent bulbous head visibly wider than a distinct, narrower neck. Thin spines displayed a small discernible head and an elongated, slender neck. Filamentous spines were longer protrusions lacking a discernible head and were morphologically consistent with filopodia. Stubby spines were short protrusions lacking a discernible neck and displayed approximately comparable length and width. Profiles that remained ambiguous after review of morphology and measured dimensions were conservatively assigned to the stubby category. Each protrusion was assigned to only one subtype, and subtype counts were reconciled with the corresponding total-spine count for each segment.

Individual spine area, perimeter, major axis, and minor axis were manually measured for every spine that could be delineated reliably, irrespective of subtype. The major axis was defined as the longest dimension of the two-dimensional spine profile, and the minor axis as the shorter perpendicular dimension. Mushroom-spine head width and neck length were measured only for protrusions previously classified as mushroom spines. Total dendritic area was determined by manually outlining the dendritic shaft and all associated, fully visualized spines within the standardized 10-µm segment.

If an eligible segment contained no spine of a particular subtype, the subtype count was recorded as zero. When no mushroom spine was identified within a sampled segment, mushroom-spine head width and neck length were also recorded as zero; these values indicate subtype absence rather than a measured dimension of zero. If a spine could be counted but a particular dimension could not be measured reliably, that dimension was treated as unavailable while the spine remained eligible for other measurable endpoints.

A subset of images was independently reviewed by more than one analyst during training and quality control to compare tracing procedures, numerical measurements, and subtype assignments. Discrepancies were reviewed with an experienced investigator and resolved by consensus before analysts continued independently. Codes were broken only after image-derived measurements were completed.

### Data handling, heatmaps, and statistical analysis

No statistical outlier rule was applied; verified extreme values were retained. Endpoint-specific n therefore varies with the number of available dendritic segments or spines. For descriptive heatmaps, measurements were averaged within animal and normalized to the 6-month vehicle mean for each endpoint, set to 100%. A fixed purple-to-green scale was used: purple indicates values at, below, or near the reference, green indicates higher values, and grey indicates a true zero rather than missing data. The heatmaps display animal level averages, normalized to the 6-month vehicle reference, allowing qualitative assessment of inter-animal consistency and regional response patterns alongside the quantitative analyses.

Data are displayed as mean ± SEM, with medians provided in the Supplementary Information. Pooled measurement-level observations were analyzed using two-sided Mann-Whitney U tests for prespecified vehicle-versus-VU0155069 comparisons within each age. Four-group distributions were evaluated by Kruskal-Wallis tests followed by Dunn adjustment of six pairwise comparisons within each endpoint family. Dunn’s procedure controlled the six prespecified pairwise comparisons within each region-and endpoint-specific analysis. Because the study was designed to characterize anatomically distinct response patterns across multiple morphometric domains, statistical interpretation emphasized consistency across related endpoints rather than applying a single study-wide adjustment. The animal was the biological replicate, with N = 5 animals per group. Endpoint-specific lowercase n values represent pooled dendritic segments, individual spines, or mushroom spines. Because these measurements are nested within animals, the reported p values characterize the sampled morphometric distributions and should be interpreted together with the biological sample size and exploratory multiregional design.

All technically valid observations were retained after verification against the corresponding source images, thereby preserving the complete range of biological and morphometric variability. Each endpoint was analyzed using all available non-missing observations, resulting in variation in measurement-level observations across endpoints, regions, and groups. The inferential tests used pooled measurement-level distributions. These measurements were nested within five animals per group and therefore do not constitute independent biological replicates. The reported p-values characterize differences among sampled morphometric observations and should be interpreted together with the biological sample size, measurement hierarchy, and exploratory multiregional design. A p value below 0.05 was considered statistically significant. Asterisks represent *p < 0.05, **p < 0.01, ***p < 0.001, and ****p < 0.0001; exact p values remain authoritative.

## Results

### Primary Auditory Cortex (A1)

Given that PLD1 attenuation improves synaptic function and underlying memory deficits due to preservation of dendritic spine integrity in the CA1 region of the hippocampus[13,14], we assessed whether similar protective effects extend to sensory cortical regions. The primary auditory cortex, A1, was one of the first that we examined to determine whether repeated VU0155069 treatment was associated with dendritic spine remodeling at the earlier and later pathological stages represented by the 6-and 12-month cohorts. The auditory cortex is among the earliest affected regions of AD and deficits in central auditory processing have been reported to precede the clinical diagnosis[33]. A1, in particular, is vulnerable to AD-related pathology, including intracellular amyloid beta 42 (Aβ_42_) in layer 5 pyramidal neurons as demonstrated in post-natal 5xFAD mice[36]. Golgi-Cox images acquired from archived specimens were used to quantify spine abundance, subtype distribution, density, and morphometric parameters in standardized 10-µm spiny dendritic segments from female 3xTg-AD mice (**Fig. 1**). Primary comparisons evaluated vehicle (Veh)-and VU0155069 (VU01)-treated mice within each age cohort using two-sided Mann-Whitney U tests. Four-group Kruskal-Wallis analyses followed by Dunn-adjusted comparisons provided additional age-cohort context.

**Figure 1.**
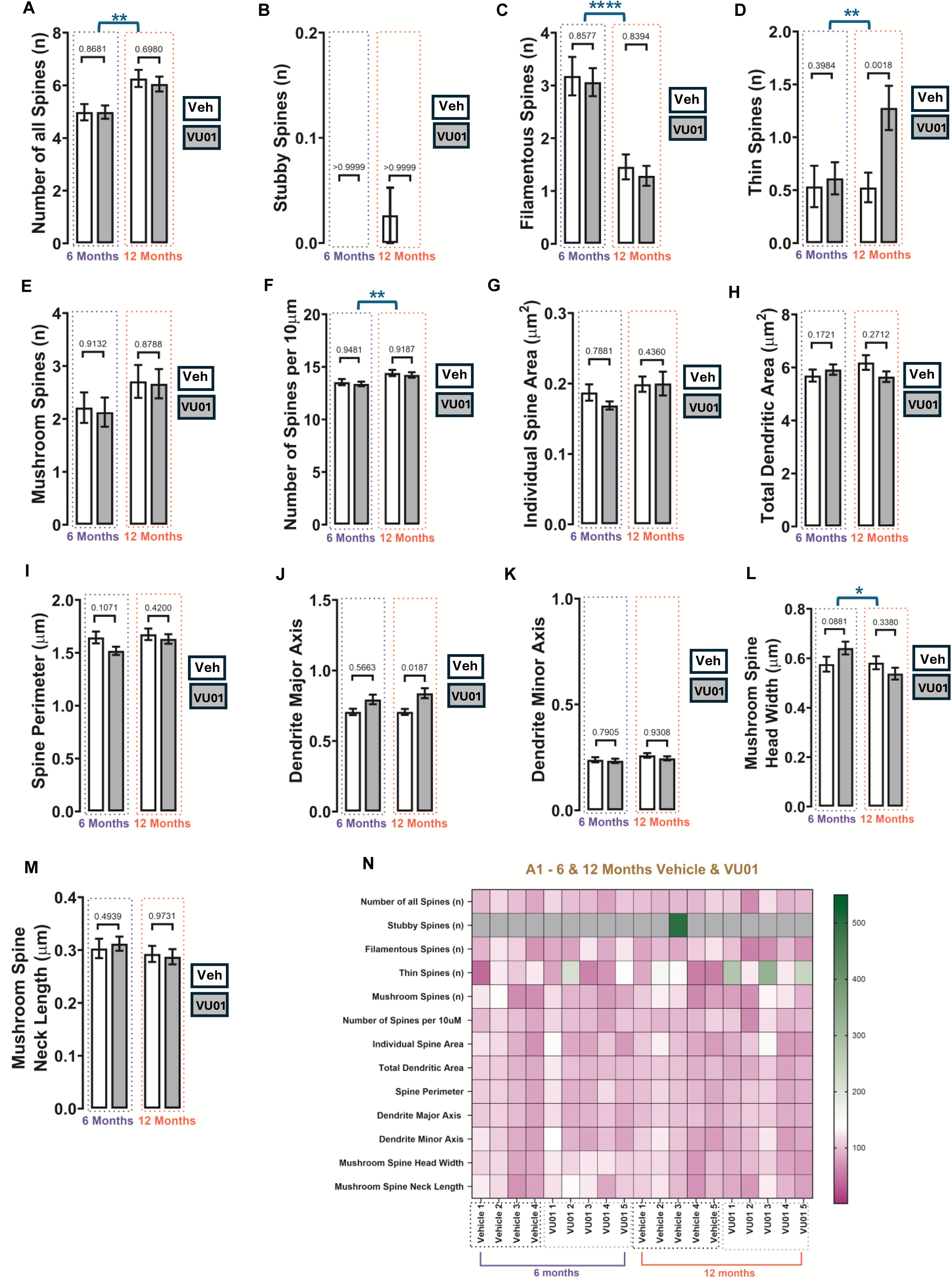
VU0155069-associated dendritic spine remodeling in the primary auditory cortex (A1) of female 3xTg-AD mice. Representative Golgi–Cox images and quantitative analyses of standardized 10-µm spiny dendritic segments from A1 are shown for 6-month vehicle, 6-month VU0155069, 12-month vehicle, and 12-month VU0155069 groups. Endpoints were: (A) total spine number; (B) stubby-spine count; (C) filamentous-spine count; (D) thin-spine count; (E) mushroom-spine count; (F) spine density per 10 µm; (G) individual spine area; (H) total dendritic area, defined as the combined two-dimensional area of the outlined dendritic shaft and associated fully visualized spines; (I) spine perimeter; (J) dendrite major axis; (K) dendrite minor axis; (L) mushroom-spine head width; and (M) mushroom-spine neck length. Data are displayed as mean ± SEM. The animal was the biological replicate, with n = 5 animals per group. Endpoint-specific n values represent pooled dendritic segments, individual spines, or mushroom spines and are provided in the Supplementary Information. Only statistically significant within-age vehicle-versus-VU0155069 comparisons are displayed above the bars. At 12 months, VU0155069 treatment was associated with increased thin-spine count (p = 0.0018; panel D) and dendrite major axis (p = 0.0187; panel J). No within-age treatment comparison was significant at 6 months. Significant Kruskal–Wallis omnibus results across the four independent age-by-treatment groups are shown separately within the corresponding panels. Four-group effects were detected for total spine number, filamentous-spine count, thin-spine count, spine density, and mushroom-spine head width. Complete Mann–Whitney U statistics and exact p values, Kruskal–Wallis H statistics, and Dunn-adjusted comparisons are provided in the Supplementary Information (**Suppl. Tables 1A** and **1B**). The heatmap displays animal-level averages normalized to the 6-month vehicle mean for the corresponding endpoint, set to 100%. Purple indicates values at, below, or near the reference; green indicates values above the reference; and grey indicates a true numerical zero rather than missing data. Each heatmap column represents one animal. The heatmap is descriptive and was not used for inferential analysis. Veh, vehicle; VU01, VU0155069.

At 6 months, VU01 treatment was not associated with a statistically significant difference in any measured A1 endpoints. These included total spine number (p = 0.8681; **Fig. 1A**); stubby (p > 0.9999; **Fig. 1B**), filamentous (p = 0.8577; **Fig. 1C**), thin (p = 0.3984; **Fig. 1D**), and mushroom spine counts (p = 0.9132; **Fig. 1E**); spine density (p = 0.9481; **Fig. 1F**); individual spine area (p = 0.7881; **Fig. 1G**); total dendritic area (p = 0.1721; **Fig. 1H**); spine perimeter (p = 0.1071; **Fig. 1I**); dendrite major axis (p = 0.5663; **Fig. 1J**); dendrite minor axis (p = 0.7905; **Fig. 1K**); mushroom-spine head width (p = 0.0881; **Fig. 1L**); and mushroom-spine neck length (p = 0.4939; **Fig. 1M**). At 12 months, VU01 treatment was associated with a selective increase in thin-spine count relative to age-matched veh treatment (p = 0.0018; **Fig. 1D**). Dendrite major axis was also increased in the VU01-treated group (p = 0. 0187; **Fig. 1J**). No statistically significant within-age treatment differences were detected for total spine number (p = 0.6980; **Fig. 1A**); stubby (p > 0.9999; **Fig. 1B**), filamentous (p = 0.8394; **Fig. 1C**), or mushroom spine counts (p = 0.8788; **Fig. 1E**); spine density (p = 0.9187; **Fig. 1F**); individual spine area (p = 0.4360; **Fig. 1G**); total dendritic area (p = 0.2712; **Fig. 1H**); spine perimeter (p = 0.4200; **Fig. 1I**); dendrite minor axis (p = 0.9308; **Fig. 1K**); mushroom-spine head width (p = 0.3380; **Fig. 1L**); or mushroom-spine neck length (p = 0.9731; **Fig. 1M**).

Four-group Kruskal–Wallis analyses identified overall distributional differences in total spine number (p = 0.0017; **Fig. 1A**), filamentous-spine count (p < 0.0001; **Fig. 1C**), thin-spine count (p = 0.0027; **Fig. 1D**), spine density (p = 0.0079; **Fig. 1F**), and mushroom-spine head width (p = 0.0231; **Fig. 1L**). No significant omnibus differences were detected for stubby-spine count (p = 0.4659; **Fig. 1B**), mushroom-spine count (p = 0.3323; **Fig. 1E**), individual spine area (p = 0.6694; **Fig. 1G**), total dendritic area (p = 0.4164; **Fig. 1H**), spine perimeter (p = 0.2922; **Fig. 1I**), dendrite major axis (p = 0.0932; **Fig. 1J**), dendrite minor axis (p = 0.8768; **Fig. 1K**), or mushroom-spine neck length (p = 0.2296; **Fig. 1M**).

The Dunn-adjusted comparisons indicated that the omnibus effects for total spine number and filamentous-spine count were driven primarily by differences between the separately processed 6-and 12-month cohorts rather than by veh-versus-VU01 differences within either age. For thin-spine count, the 12-month VU01 group showed significant differences versus the 6-month groups and the 12-month vehicle group, supporting a treatment-associated effect within the later-stage cohort. After Dunn’s adjustment, there were no significant changes for spine density parameters. Mushroom-spine head width showed a more limited difference between the 6-month and 12-month VU01 groups. Complete H statistics, mean-rank differences, and adjusted p values are provided in the Supplementary Information (**Suppl. Tables 1A** and **1B**).

Together, these findings identify A1 as a region with a selective, stage-dependent response to VU01. Although A1 spine architecture remained comparatively stable at 6 months, treatment at 12 months increased thin-spine abundance and dendrite major axis, indicating recruitment of structural plasticity during the later pathological stage. The four-group analyses further demonstrated age-associated reorganization of spine abundance and phenotype distribution. Thus, A1 retains the capacity for targeted structural remodeling, with the most prominent VU01-associated response emerging during later-stage pathology.

### Secondary Auditory Cortex (A2)

The secondary auditory cortex, A2, contributes to higher-order auditory integration and has been implicated in the consolidation and retention of auditory fear memories[54,55]. A role for PLD isoforms in fear memories was demonstrated by our group in rats[9–11], further increasing the relevance of studying this brain region for dendritic spine integrity in modulating synaptic function. Building on the VU01-associated improvements in LTP, LTD, novel-object recognition, and fear-conditioning performance reported across two prior peer-reviewed studies[13,14], the present multiregional analysis examines how this therapeutic response is reflected in dendritic spine remodeling across the cortical, hippocampal, and amygdalar circuits supporting these functions.

A2 was examined to determine whether repeated VU01 treatment was associated with dendritic spine remodeling in a higher-order auditory region at the earlier and later pathological stages represented by the 6-and 12-month cohorts (**Fig. 2**).

**Figure 2.**
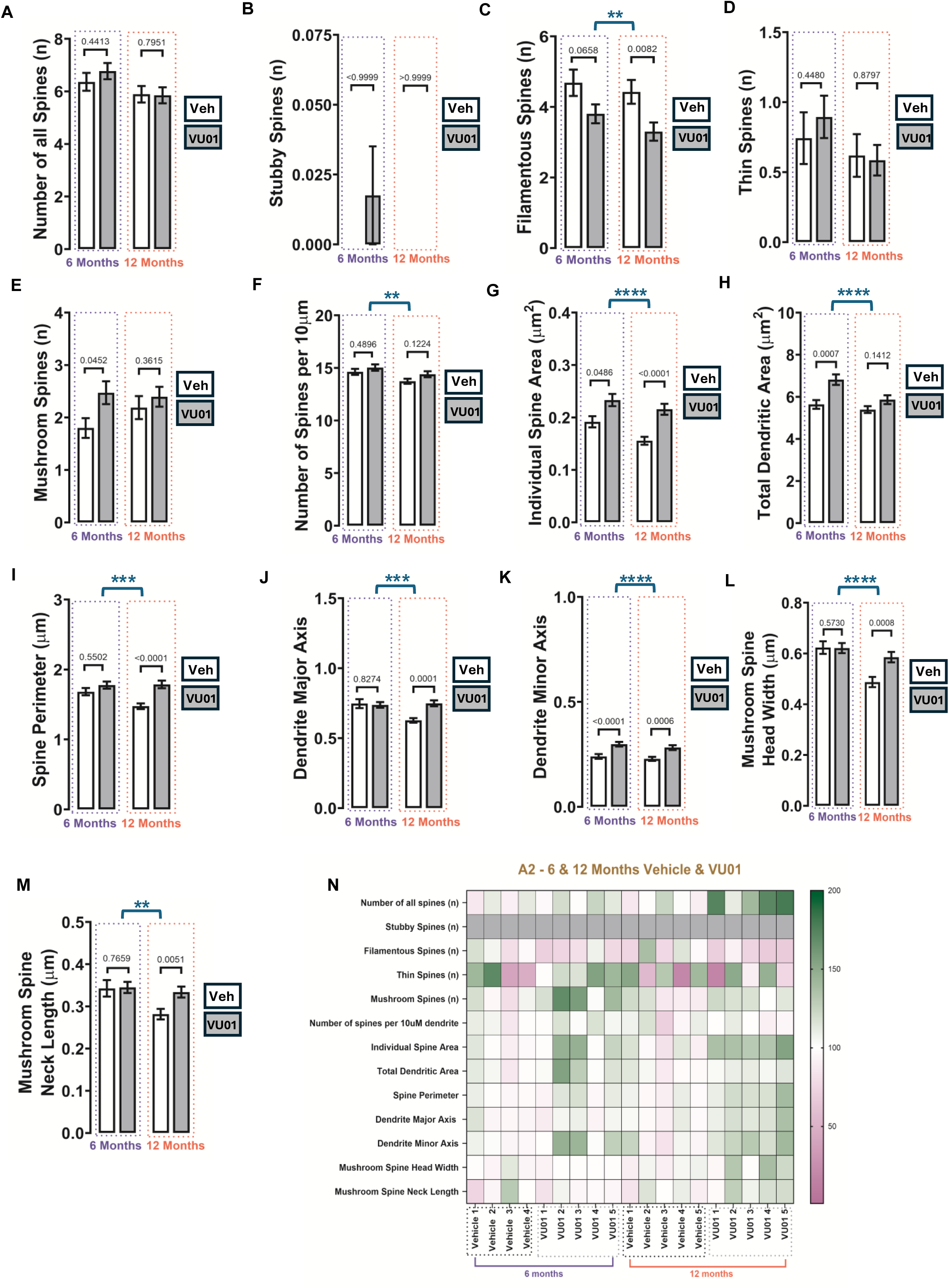
VU0155069-associated dendritic spine remodeling in the secondary auditory (A2) cortex of female 3xTg-AD mice. Representative Golgi–Cox images and quantitative analyses of standardized 10-µm spiny dendritic segments from A2 are shown for 6-month vehicle, 6-month VU0155069, 12-month vehicle, and 12-month VU0155069 groups. Endpoints were: (A) total spine number; (B) stubby-spine count; (C) filamentous-spine count; (D) thin-spine count; (E) mushroom-spine count; (F) spine density per 10 µm; (G) individual spine area; (H) total dendritic area, defined as the combined two-dimensional area of the outlined dendritic shaft and associated fully visualized spines; (I) spine perimeter; (J) dendrite major axis; (K) dendrite minor axis; (L) mushroom-spine head width; and (M) mushroom-spine neck length. Data are displayed as mean ± SEM. The animal was the biological replicate, with n = 5 animals per group. Endpoint-specific n values represent pooled dendritic segments, individual spines, or mushroom spines and are provided in the Supplementary Information. Only statistically significant within-age vehicle-versus-VU0155069 comparisons are displayed above the bars. At 6 months, VU0155069 treatment was associated with increased mushroom-spine count (p = 0.0452; panel E), individual spine area (p = 0.0486; panel G), total dendritic area (p = 0.0007; panel H), and dendrite minor axis (p < 0.0001; panel K). At 12 months, treatment was associated with reduced filamentous-spine count (p = 0.0082; panel C) and increased individual spine area (p < 0.0001; panel G), spine perimeter (p < 0.0001; panel I), dendrite major axis (p = 0.0001; panel J), dendrite minor axis (p = 0.0006; panel K), mushroom-spine head width (p = 0.0008; panel L), and mushroom-spine neck length (p = 0.0051; panel M). Significant Kruskal–Wallis omnibus results across the four independent age-by-treatment groups are shown separately within the corresponding panels. Complete Mann– Whitney U statistics and exact p values, Kruskal–Wallis statistics, and Dunn-adjusted comparisons are provided in the Supplementary Information (**Suppl. Tables 2A** and **2B**). The heatmap displays animal-level averages normalized to the 6-month vehicle mean for the corresponding endpoint, set to 100%. Purple indicates values at, below, or near the reference; green indicates values above the reference; and grey indicates a true numerical zero rather than missing data. Each heatmap column represents one animal. The heatmap is descriptive and was not used for inferential analysis. Veh, vehicle; VU01, VU0155069.

At 6 months, VU01 treatment was associated with increased mushroom-spine count (p = 0.0452; **Fig. 2E**), individual spine area (p = 0.0486; **Fig. 2G**), total dendritic area (p = 0.0007; **Fig. 2H**), and dendrite minor axis (p < 0.0001; **Fig. 2K**). No statistically significant within-age treatment differences were detected for total spine number (p = 0.4413; **Fig. 2A**), stubby-spine count (p > 0.9999; **Fig. 2B**), filamentous-spine count (p = 0.0658; **Fig. 2C**), thin-spine count (p = 0.4480; **Fig. 2D**), spine density (p = 0.4896; **Fig. 2F**), spine perimeter (p = 0.5502; **Fig. 2I**), dendrite major axis (p = 0.8274; **Fig. 2J**), mushroom-spine head width (p = 0.5730; **Fig. 2L**), or mushroom-spine neck length (p = 0.7659; **Fig. 2M**).

At 12 months, VU0155069 treatment was associated with reduced filamentous-spine count (p = 0.0082; **Fig. 2C**) and increased individual spine area (p < 0.0001; **Fig. 2G**), spine perimeter (p < 0.0001; **Fig. 2I**), dendrite major axis (p = 0.0001; **Fig. 2J**), dendrite minor axis (p = 0.0006; **Fig. 2K**), mushroom-spine head width (p = 0.0008; **Fig. 2L**), and mushroom-spine neck length (p = 0.0051; **Fig. 2M**). No statistically significant within-age differences were detected for total spine number (p = 0.7951; **Fig. 2A**), stubby-spine count (p > 0.9999; **Fig. 2B**), thin-spine count (p = 0.8797; **Fig. 2D**), mushroom-spine count (p = 0.3615; **Fig. 2E**), spine density (p = 0.1224; **Fig. 2F**), or total dendritic area (p = 0.1412; **Fig. 2H**).

Four-group Kruskal-Wallis analyses identified overall distributional differences in filamentous-spine count (p = 0.0089; **Fig. 2C**), spine density (p = 0.0095; **Fig. 2F**), individual spine area (p < 0.0001; **Fig. 2G**), total dendritic area (p < 0.0001; **Fig. 2H**), spine perimeter (p = 0.0001; **Fig. 2I**), dendrite major axis (p = 0.0003; **Fig. 2J**), dendrite minor axis (p < 0.0001; **Fig. 2K**), mushroom-spine head width (p < 0.0001; **Fig. 2L**), and mushroom-spine neck length (p = 0.0062; **Fig. 2M**). No significant omnibus differences were detected for total spine number (p = 0.0678; **Fig. 2A**), stubby-spine count (p = 0.5162; **Fig. 2B**), thin-spine count (p = 0.5097; **Fig. 2D**), or mushroom-spine count (p = 0.1395; **Fig. 2E**).

At 6 months, the difference in total dendritic area remained significant after Dunn adjustment, as did the difference in dendrite minor axis. At 12 months, Dunn-adjusted vehicle-versus-VU0155069 differences remained significant for filamentous-spine count, individual spine area, spine perimeter, dendrite major axis, dendrite minor axis, mushroom-spine head width, and mushroom-spine neck length. The significant omnibus result for spine density was driven by a difference between the 6-month VU0155069 and 12-month vehicle groups rather than a within-age contrast. Complete mean-rank differences and adjusted p values are provided in the Supplementary Information (**Suppl. Tables 2A** and **2B**).

Together, the A1 and A2 findings demonstrate that VU0155069 engages distinct, stage-dependent structural remodeling programs across the auditory cortex. A1 remained structurally stable at 6 months and developed a focused later-stage response characterized by increased thin-spine abundance and dendrite major axis. In contrast, A2 responded at both pathological stages and across a broader range of structural domains. At 6 months, VU0155069 increased mushroom-spine abundance, individual spine area, total dendritic area, and dendrite minor axis, indicating early recruitment of both subtype-specific and morphometric plasticity. By 12 months, the A2 response shifted toward reduced filamentous-spine representation together with coordinated increases in individual-spine and mushroom-spine dimensions. Thus, VU0155069 produced complementary responses within the auditory cortex: a selective later-stage remodeling program in A1 and an earlier, multidimensional, and sustained structural response in A2. These findings extend the previously demonstrated functional efficacy of VU0155069 by showing that anatomically related auditory regions contribute differently to the broader structural resilience of distributed sensory and associative networks.

### Primary Visual Cortex (V1)

The broader rationale for assessing visual cortex was presented earlier [27,42–44], but within this cortex, the primary visual cortex (V1) is the most well-understood. It is the earliest region to receive and process information [56]. Additionally, V1 has shown an increased affinity for binding to amyloid-beta (Aβ), which contributes to its deposition and eventual disruption of visual perception [57]. Significant Aβ pathology has been observed in the V1 region with increased microglial and phagocytic activity in 5xFAD mice models [58]. Thus, we examined whether repeated VU0155069 treatment was associated with dendritic spine remodeling in an early visual-processing region at the 6-and 12-month pathological stages (**Fig. 3**).

**Figure 3.**
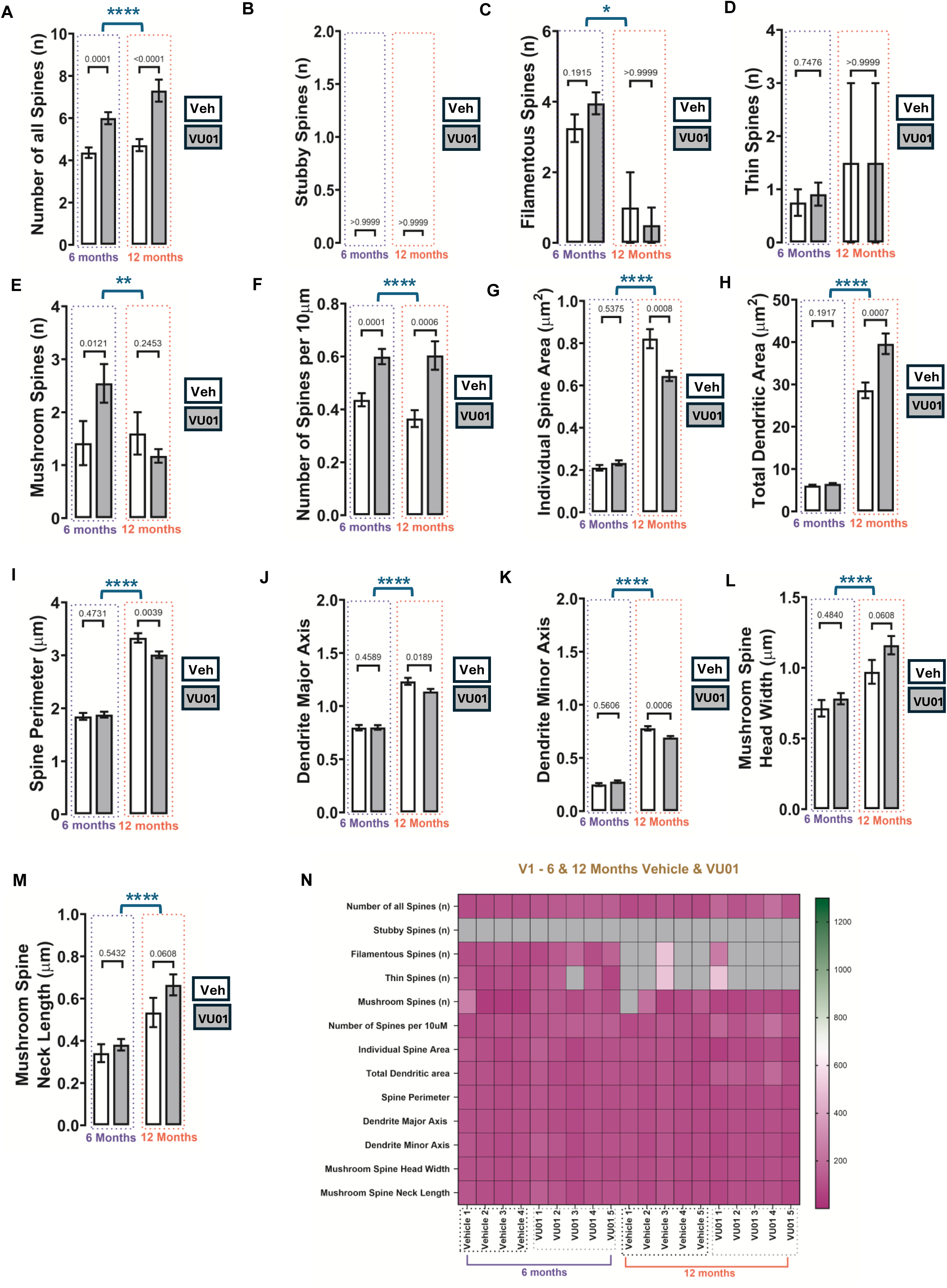
VU0155069-associated dendritic spine remodeling in the primary visual cortex (V1) of female 3xTg-AD mice. Representative Golgi–Cox images and quantitative analyses of standardized 10-µm spiny dendritic segments from V1 are shown for 6-month vehicle, 6-month VU0155069, 12-month vehicle, and 12-month VU0155069 groups. Endpoints were: (A) total spine number; (B) stubby-spine count; (C) filamentous-spine count; (D) thin-spine count; (E) mushroom-spine count; (F) spine density per 10 µm; (G) individual spine area; (H) total dendritic area, defined as the combined two-dimensional area of the outlined dendritic shaft and associated fully visualized spines; (I) spine perimeter; (J) dendrite major axis; (K) dendrite minor axis; (L) mushroom-spine head width; and (M) mushroom-spine neck length. Data are displayed as mean ± SEM. The animal was the biological replicate, with n = 5 animals per group. Endpoint-specific n values represent pooled dendritic segments, individual spines, or mushroom spines and are provided in the Supplementary Information. Only statistically significant within-age vehicle-versus-VU0155069 comparisons are displayed above the bars. At 6 months, treatment was associated with increased total spine number (p = 0.0001; panel A), mushroom-spine count (p = 0.0121; panel E), and spine density (p = 0.0001; panel F). At 12 months, treatment was associated with increased total spine number (p < 0.0001; panel A), spine density (p = 0.0006; panel F), and total dendritic area (p = 0.0007; panel H), and with reduced individual spine area (p = 0.0008; panel G), spine perimeter (p = 0.0039; panel I), dendrite major axis (p = 0.0189; panel J), and dendrite minor axis (p = 0.0006; panel K). Significant Kruskal–Wallis omnibus results across the four independent age-by-treatment groups are shown separately in the corresponding panels. Complete Mann–Whitney U statistics and exact p values, Kruskal–Wallis statistics, and Dunn-adjusted comparisons are provided in the Supplementary Information (**Suppl. Tables 3A** and **3B**). The heatmap displays animal-level averages normalized to the 6-month vehicle mean for the corresponding endpoint, set to 100%. Purple indicates values at, below, or near the reference; green indicates values above the reference; and grey indicates a true numerical zero rather than missing data. Each heatmap column represents one animal. The heatmap is descriptive and was not used for inferential analysis. Veh, vehicle; VU01, VU0155069.

At 6 months, VU0155069 treatment was associated with increased total spine number (p = 0.0001; **Fig. 3A**), mushroom-spine count (p = 0.0121; **Fig. 3E**), and spine density (p = 0.0001; **Fig. 3F**). No statistically significant within-age treatment differences were detected for stubby-spine count (p > 0.9999; **Fig. 3B**), filamentous-spine count (p = 0.1915; **Fig. 3C**), thin-spine count (p = 0.7476; **Fig. 3D**), individual spine area (p = 0.5375; **Fig. 3G**), total dendritic area (p = 0.1917; **Fig. 3H**), spine perimeter (p = 0.4731; **Fig. 3I**), dendrite major axis (p = 0.4589; **Fig. 3J**), dendrite minor axis (p = 0.5606; **Fig. 3K**), mushroom-spine head width (p = 0.4840; **Fig. 3L**), or mushroom-spine neck length (p = 0.5432; **Fig. 3M**).

At 12 months, VU0155069 treatment was associated with increased total spine number (p < 0.0001; **Fig. 3A**), spine density (p = 0.0006; **Fig. 3F**), and total dendritic area (p = 0.0007; **Fig. 3H**). In contrast, individual spine area (p = 0.0008; **Fig. 3G**), spine perimeter (p = 0.0039; **Fig. 3I**), dendrite major axis (p = 0.0189; **Fig. 3J**), and dendrite minor axis (p = 0.0006; **Fig. 3K**) were reduced relative to age-matched vehicle treatment. No statistically significant within-age differences were detected for stubby-spine count (p > 0.9999; **Fig. 3B**), filamentous-spine count (p > 0.9999; **Fig. 3C**), thin-spine count (p = 0.4831; **Fig. 3D**), mushroom-spine count (p = 0.2453; **Fig. 3E**), mushroom-spine head width (p = 0.0608; **Fig. 3L**), or mushroom-spine neck length (p = 0.0608; **Fig. 3M**).

Four-group Kruskal-Wallis analyses identified overall distributional differences in total spine number (p < 0.0001; **Fig. 3A**), filamentous-spine count (p = 0.0130; **Fig. 3C**), mushroom-spine count (p = 0.0034; **Fig. 3E**), spine density (p < 0.0001; **Fig. 3F**), individual spine area (p < 0.0001; **Fig. 3G**), total dendritic area (p < 0.0001; **Fig. 3H**), spine perimeter (p < 0.0001; **Fig. 3I**), dendrite major axis (p < 0.0001; **Fig. 3J**), dendrite minor axis (p < 0.0001; **Fig. 3K**), mushroom-spine head width (p < 0.0001; **Fig. 3L**), and mushroom-spine neck length (p < 0.0001; **Fig. 3M**). Stubby-spine count did not yield a Kruskal-Wallis statistic as all observed values were zero, resulting in no variance and making group comparisons impossible (**Fig. 3B**). No distributional differences were found in thin-spine count (p = 0.9275; **Fig. 3D**).

Dunn-adjusted comparisons supported treatment-associated changes in total spine number and spine density at both ages, and in mushroom-spine count at 6 months. For several spine-size and total-dendritic-area endpoints, the significant omnibus effects were driven predominantly by separation between the independently processed 6-and 12-month cohorts. The direct 12-month vehicle-versus-VU0155069 comparisons for individual spine area, total dendritic area, spine perimeter, and dendrite major and minor axes did not remain significant after Dunn adjustment within the four-group framework. Complete mean-rank differences and adjusted p values are provided in the Supplementary Information (**Suppl. Tables 3A** and **3B**).

Together, the V1 findings reveal a robust, stage-dependent structural response to VU0155069, characterized by increased spine abundance at both pathological stages and a distinct reorganization of spine morphology during later pathology. At 6 months, treatment increased total spine number, mushroom-spine count, and spine density. At 12 months, increased total spine number, spine density, and total dendritic area occurred together with lower values across several individual-spine dimensional measures. Importantly, this coordinated increase in spine abundance with a reduction in average individual-spine size could be identified only through the comprehensive morphometric approach used in the present study, which evaluated the number, phenotype, density, and geometry of spines within the same regional framework. The inverse relationship is consistent with a two-stage adaptive response in which enlargement of surviving spines is followed by the regeneration or recruitment of a larger population of comparatively small spines[59]. This model provides a plausible structural basis for the VU0155069-associated synaptic resilience demonstrated previously and establishes a focused hypothesis for future longitudinal imaging and regionally matched physiological studies. Such studies will complement the present findings by defining the temporal sequence and functional contribution of the V1 remodeling program.

### Secondary Visual Cortex (V2)

The secondary visual cortex (V2) integrates higher-order visual features; where we tested whether treatment effects shift from subtype-specific to global structural changes with advancing pathology. The V2 region integrates higher order complex visual features such as color and object orientation which it receives from V1. Although V2 specific neurodegenerative effects are not well known, prior research report Aβ-associated detriments in V1[27,42–44]. Golgi-Cox images acquired from archived specimens were used to quantify spine abundance, subtype distribution, density, and morphometric parameters in standardized 10-µm spiny dendritic segments (**Fig. 4**).

**Figure 4.**
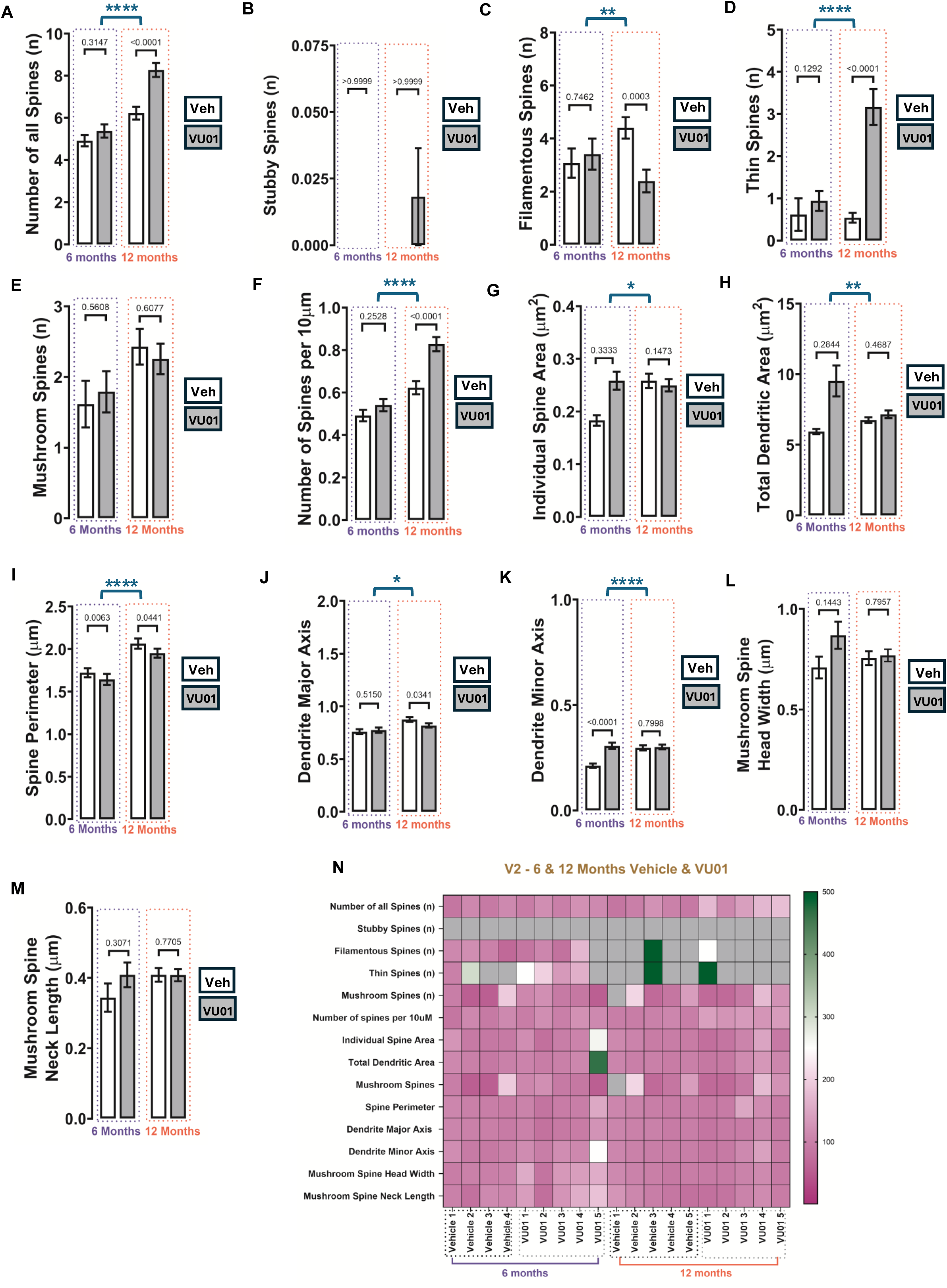
VU0155069-associated dendritic spine remodeling in the secondary visual cortex (V2) of female 3xTg-AD mice. Representative Golgi–Cox images and quantitative analyses of standardized 10-µm spiny dendritic segments from V2 are shown for 6-month vehicle, 6-month VU0155069, 12-month vehicle, and 12-month VU0155069 groups. Endpoints were: (A) total spine number; (B) stubby-spine count; (C) filamentous-spine count; (D) thin-spine count; (E) mushroom-spine count; (F) spine density per 10 µm; (G) individual spine area; (H) total dendritic area; (I) spine perimeter; (J) dendrite major axis; (K) dendrite minor axis; (L) mushroom-spine head width; and (M) mushroom-spine neck length. Data are displayed as mean ± SEM. The animal was the biological replicate, with n = 5 animals per group. Endpoint-specific observation values are provided in the Supplementary Information. Only significant within-age comparisons are displayed above bars. Significant Kruskal–Wallis omnibus results across the four independent age-by-treatment groups are shown separately within the corresponding panels. Four-group effects were detected for total spine number, filamentous-spine count, thin-spine count, spine density, and mushroom-spine head width. Complete Mann–Whitney U statistics and exact p values, Kruskal– Wallis H statistics, and Dunn-adjusted comparisons are provided in the Supplementary Information (**Suppl. Tables 4A** and **4B**). The heatmap displays animal-level averages normalized to the 6-month vehicle mean for the corresponding endpoint, set to 100%. Purple indicates values at, below, or near the reference; green indicates values above the reference; and grey indicates a true numerical zero rather than missing data. Each heatmap column represents one animal. The heatmap is descriptive and was not used for inferential analysis. Veh, vehicle; VU01, VU0155069.

At 6 months, VU0155069 treatment was associated with reduced spine perimeter (p = 0.0063; **Fig. 4I**) and increased dendrite minor axis (p < 0.0001; **Fig. 4K**). No statistically significant within-age treatment differences were detected for total spine number (p = 0.3147; **Fig. 4A**), stubby-spine count (p > 0.9999; **Fig. 4B**), filamentous-spine count (p = 0.7462; **Fig. 4C**), thin-spine count (p = 0.1292; **Fig. 4D**), mushroom-spine count (p = 0.5608; **Fig. 4E**), spine density (p = 0.2528; **Fig. 4F**), individual spine area (p = 0.3333; **Fig. 4G**), total dendritic area (p = 0.2844; **Fig. 4H**), dendrite major axis (p = 0.5150; **Fig. 4J**), mushroom-spine head width (p = 0.1443; **Fig. 4L**), or mushroom-spine neck length (p = 0.3071; **Fig. 4M**).

At 12 months, VU0155069 treatment was associated with increased total spine number (p < 0.0001; **Fig. 4A**), thin-spine count (p < 0.0001; **Fig. 4D**), and spine density (p < 0.0001; **Fig. 4F**), together with reduced filamentous-spine count (p = 0.0003; **Fig. 4C**), spine perimeter (p = 0.0441; **Fig. 4I**), and dendrite major axis (p = 0.0341; **Fig. 4J**). No statistically significant within-age differences were detected for stubby-spine count (p > 0.9999; **Fig. 4B**), mushroom-spine count (p = 0.6077; **Fig. 4E**), individual spine area (p = 0.1473; **Fig. 4G**), total dendritic area (p = 0.4687; **Fig. 4H**), dendrite minor axis (p = 0.7998; **Fig. 4K**), mushroom-spine head width (p = 0.7957; **Fig. 4L**), or mushroom-spine neck length (p = 0.7705; **Fig. 4M**).

Four-group Kruskal–Wallis analyses identified overall differences in total spine number (p < 0.0001; **Fig. 4A**), filamentous-spine count (p = 0.0015; **Fig. 4C**), thin-spine count (p < 0.0001; **Fig. 4D**), spine density (p < 0.0001; **Fig. 4F**), individual spine area (p = 0.0137; **Fig. 4G**), total dendritic area (p = 0.0094; **Fig. 4H**), spine perimeter (p < 0.0001; **Fig. 4I**), dendrite major axis (p = 0.0109; **Fig. 4J**), and dendrite minor axis (p < 0.0001; **Fig. 4K**). No significant omnibus differences were detected for stubby-spine count (p = 0.7574; **Fig. 4B**), mushroom-spine count (p = 0.1526; **Fig. 4E**), mushroom-spine head width (p = 0.4231; **Fig. 4L**), or mushroom-spine neck length (p = 0.6308; **Fig. 4M**).

Dunn-adjusted comparisons supported the 12-month vehicle-versus-VU0155069 differences in total spine number, filamentous-spine count, thin-spine count, and spine density. At 6 months, the difference in dendrite minor axis remained significant after Dunn adjustment, whereas the spine-perimeter comparison did not. Omnibus effects for individual spine area and total dendritic area primarily reflected differences between independently processed age cohorts. Complete mean-rank differences and adjusted p values are provided in the Supplementary Information (**Suppl. Tables 4A** and **4B**).

Together, the V2 findings indicate limited morphometric change at 6 months and a more pronounced subtype-and abundance-associated response at 12 months. The later-stage response comprised increased total spine number, thin-spine count, and spine density, together with reduced filamentous-spine count and modest reductions in spine perimeter and major axis. Thus, V2 exhibited a stage-dependent and selective structural profile rather than a uniform change across all measured endpoints. V2 exhibited relatively limited changes at 6 months and a more focused 12-month response characterized by increased total spine abundance, increased thin spines, decreased filamentous spines, and selective changes in perimeter-and axis-related geometry. These findings suggest that the secondary visual cortex responds to PLD1 attenuation in a stage-dependent but more subtype-selective manner than the primary visual cortex.

The distinct V1 and V2 profiles may reflect their complementary positions within the visual hierarchy. As the initial cortical recipient and distributor of visual information, adult V1 maintains a carefully regulated balance between plasticity and stability for the preservation of the organized retinotopic representation, whereas downstream visual regions integrate increasingly complex visual features and may engage broader remodeling programs[60]. Conversely, the V2 region downstream potentially exhibits greater variability, which allows for more apparent effects with attenuation of PLD1.

### Hippocampal CA2 Subfield

The CA2 region in particular, is essential in social recognition memory and has been found to be affected in AD[61]. Specifically, there is a reduction in social recognition memory in AD due to a loss of Perineuronal net (PNNs) around CA2 pyramidal neurons. Thus, we studied the hippocampal CA2 subfield to determine whether repeated VU0155069 treatment was associated with dendritic spine remodeling in a functionally specialized hippocampal region at the 6-and 12-month pathological stages (**Fig. 5**).

**Figure 5.**
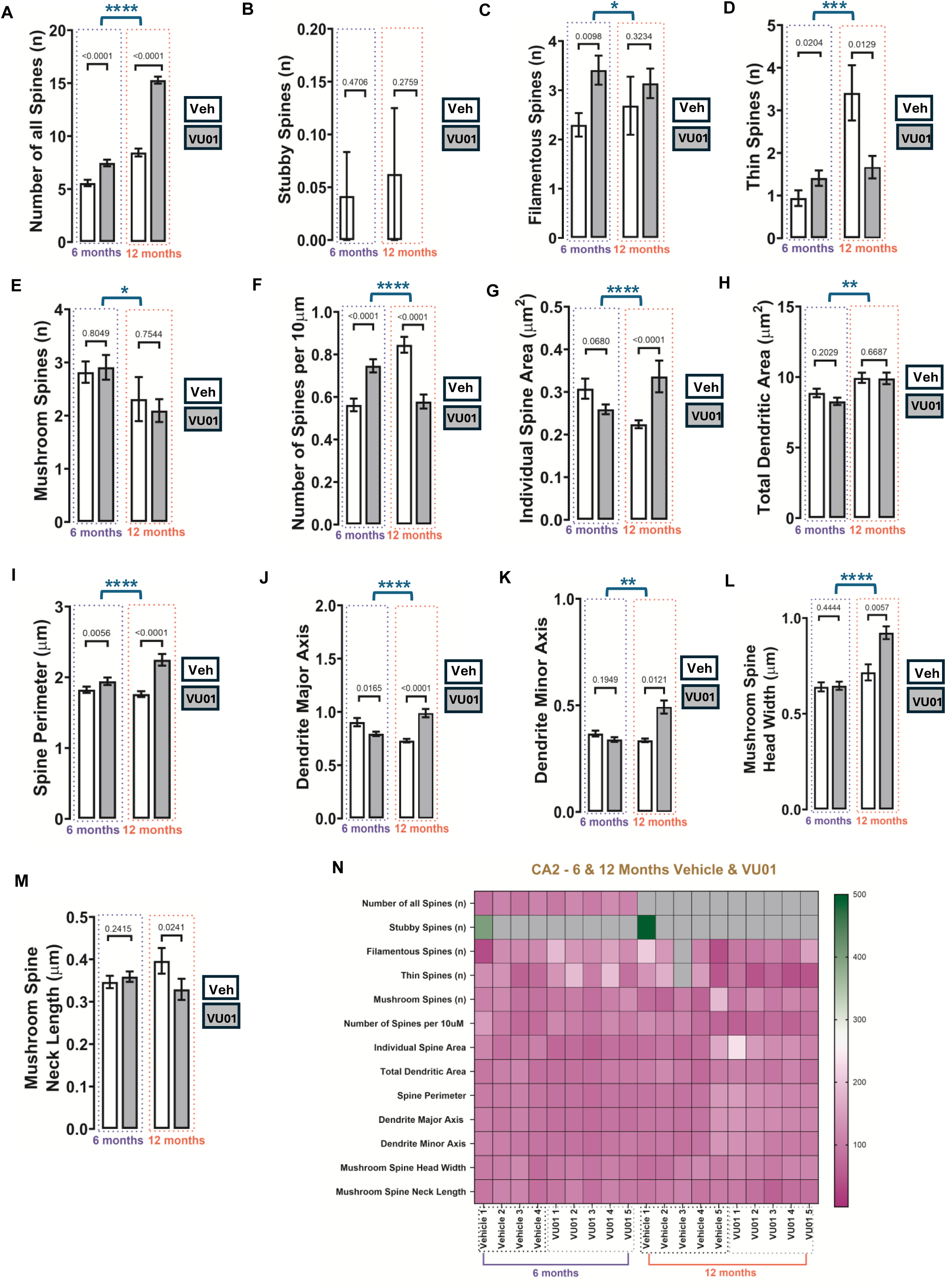
VU0155069-associated dendritic spine remodeling in the hippocampal CA2 subfield of female 3xTg-AD mice. Representative Golgi–Cox images and quantitative analyses of standardized 10-µm spiny dendritic segments from CA2 are shown for 6-month vehicle, 6-month VU0155069, 12-month vehicle, and 12-month VU0155069 groups. Endpoints were: (A) total spine number; (B) stubby-spine count; (C) filamentous-spine count; (D) thin-spine count; (E) mushroom-spine count; (F) spine density per 10 µm; (G) individual spine area; (H) total dendritic area; (I) spine perimeter; (J) dendrite major axis; (K) dendrite minor axis; (L) mushroom-spine head width; and (M) mushroom-spine neck length. Data are displayed as mean ± SEM. The animal was the biological replicate, with n = 5 animals per group. Endpoint-specific observation values are provided in the Supplementary Information. Only significant within-age comparisons are displayed above bars. Significant Kruskal–Wallis omnibus results across the four independent age-by-treatment groups are shown separately within the corresponding panels. Four-group effects were detected for total spine number, filamentous-spine count, thin-spine count, spine density, and mushroom-spine head width. Complete Mann–Whitney U statistics and exact p values, Kruskal– Wallis H statistics, and Dunn-adjusted comparisons are provided in the Supplementary Information (**Suppl. Tables 5A** and **5B**). The heatmap displays animal-level averages normalized to the 6-month vehicle mean for the corresponding endpoint, set to 100%. Purple indicates values at, below, or near the reference; green indicates values above the reference; and grey indicates a true numerical zero rather than missing data. Each heatmap column represents one animal. The heatmap is descriptive and was not used for inferential analysis. Veh, vehicle; VU01, VU0155069.

At 6 months, VU0155069 treatment was associated with increased total spine number (p < 0.0001; **Fig. 5A**), filamentous-spine count (p = 0.0098; **Fig. 5C**), thin-spine count (p = 0.0204; **Fig. 5D**), spine density (p < 0.0001; **Fig. 5F**), and spine perimeter (p = 0.0056; **Fig. 5I**), together with reduced dendrite major axis (p = 0.0165; **Fig. 5J**). No statistically significant within-age differences were detected for stubby-spine count (p = 0.4706; **Fig. 5B**), mushroom-spine count (p = 0.8049; **Fig. 5E**), individual spine area (p = 0.0680; **Fig. 5G**), total dendritic area (p = 0.2029; **Fig. 5H**), dendrite minor axis (p = 0.1949; **Fig. 5K**), mushroom-spine head width (p = 0.4444; **Fig. 5L**), or mushroom-spine neck length (p = 0.2415; **Fig. 5M**).

At 12 months, VU0155069 treatment was associated with increased total spine number (p < 0.0001; **Fig. 5A**), individual spine area (p < 0.0001; **Fig. 5G**), spine perimeter (p < 0.0001; **Fig. 5I**), dendrite major axis (p < 0.0001; **Fig. 5J**), dendrite minor axis (p = 0.0121; **Fig. 5K**), and mushroom-spine head width (p = 0.0057; **Fig. 5L**). In contrast, thin-spine count (p = 0.0129; **Fig. 5D**), spine density (p < 0.0001; **Fig. 5F**), and mushroom-spine neck length (p = 0.0241; **Fig. 5M**) were reduced. No significant within-age differences were detected for stubby-spine count (p = 0.2759; **Fig. 5B**), filamentous-spine count (p = 0.3234; **Fig. 5C**), mushroom-spine count (p = 0.7544; **Fig. 5E**), or total dendritic area (p = 0.6687; **Fig. 5H**).

Four-group Kruskal-Wallis analyses identified overall differences in total spine number (p < 0.0001; **Fig. 5A**), filamentous-spine count (p = 0.0461; **Fig. 5C**), thin-spine count (p = 0.0004; **Fig. 5D**), mushroom-spine count (p = 0.0198; **Fig. 5E**), spine density (p < 0.0001; **Fig. 5F**), individual spine area (p < 0.0001; **Fig. 5G**), total dendritic area (p = 0.0063; **Fig. 5H**), spine perimeter (p < 0.0001; **Fig. 5I**), dendrite major axis (p < 0.0001; **Fig. 5J**), dendrite minor axis (p = 0.0038; **Fig. 5K**), and mushroom-spine head width (p < 0.0001; **Fig. 5L**). No significant omnibus differences were detected for stubby-spine count (p = 0.1989; **Fig. 5B**) or mushroom-spine neck length (p = 0.0680; **Fig. 5M**).

The 12-month vehicle-versus-VU0155069 differences in individual spine area, spine perimeter, and dendrite major axis remained significant after Dunn adjustment, whereas the corresponding direct contrasts for dendrite minor axis and mushroom-spine head width did not. Complete mean-rank differences and adjusted p values are provided in the Supplementary Information (**Suppl. Tables 5A** and **5B**).

Together, the CA2 findings identify a broad, stage-dependent remodeling program that evolves from increased spine abundance and subtype diversity at 6 months to coordinated reorganization of spine number and geometry at 12 months. The later-stage combination of increased total spine number with reduced spine density is particularly informative because it suggests that the additional spines may be distributed across an expanded dendritic substrate rather than concentrated within existing segments. We therefore propose that PLD1 attenuation releases a constraint on dendritic growth, promoting greater branching or arbor complexity and thereby increasing the total synaptic substrate available to CA2 circuitry[62]. This model is supported by evidence that PLD1 negatively regulates dendritic branching in hippocampal neurons and that PLD1 knockdown or pharmacological inhibition increases dendritic complexity[63]. Given the specialized dendritic organization and strong cortical innervation of CA2 pyramidal neurons, arbor-level remodeling could substantially influence information integration within the cortico-hippocampal circuit[64,65]. Future whole-neuron reconstruction and Sholl analysis will define this component of the CA2 resilience response and complement the multidimensional spine profile revealed here.

### Hippocampal CA3 Subfield

The hippocampal CA3 subfield, with its recurrent architecture, supporting pattern completion and often shows distinct vulnerability from CA2 and CA1, and a subfield characterized by its recurrent circuitry and role in pattern completion[39,66–69] and, therefore, important for understanding the effects of how repeated VU0155069 treatment impacted dendritic spine remodeling at the 6-and 12-month pathological stages, using images acquired from archived specimens were used to quantify spine abundance, subtype distribution, density, and morphometric parameters in standardized 10-µm spiny dendritic segments (**Fig. 6**).

**Figure 6.**
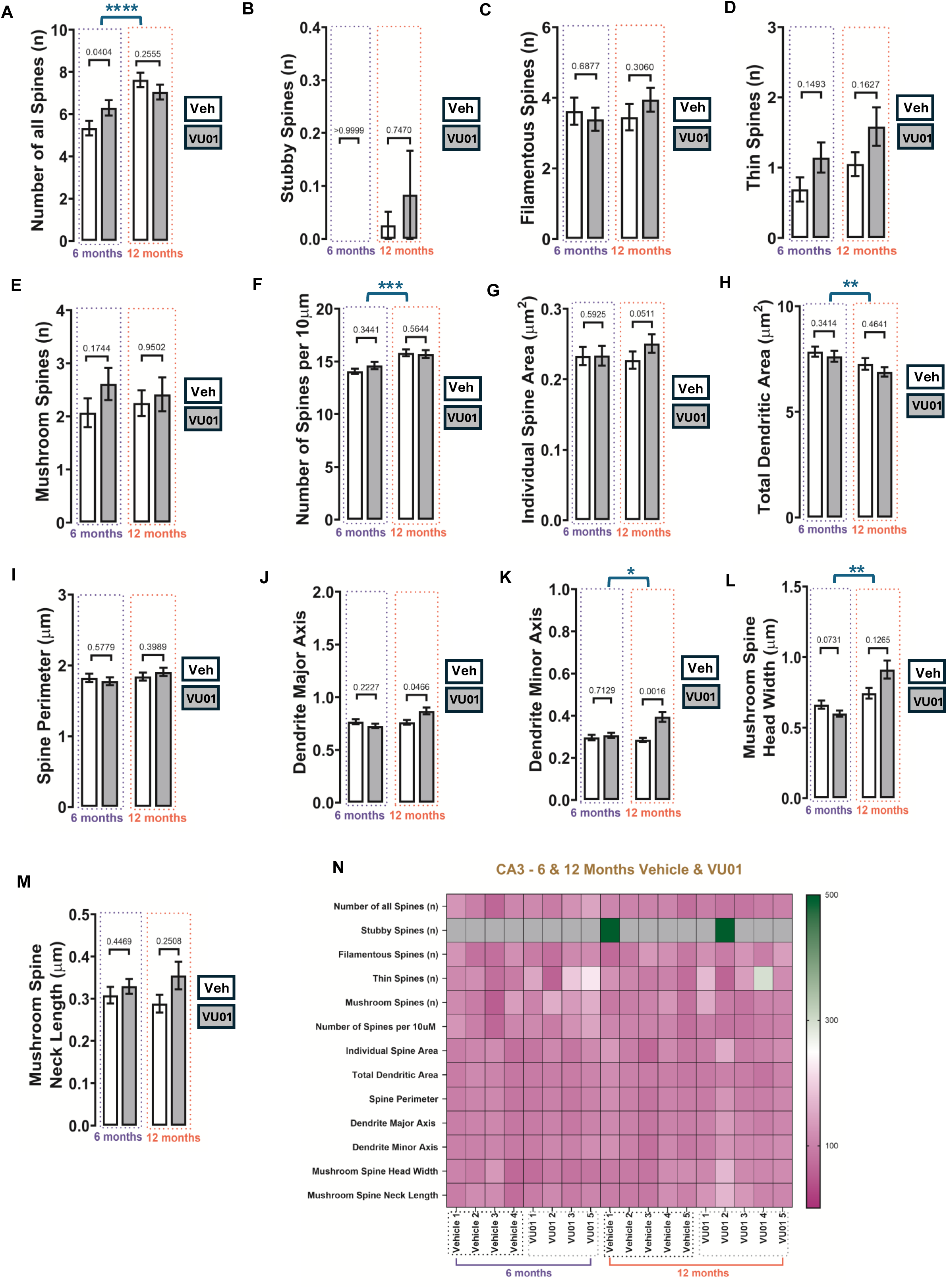
VU0155069-associated dendritic spine remodeling in the hippocampal CA3 subfield of female 3xTg-AD mice. Representative Golgi–Cox images and quantitative analyses of standardized 10-µm spiny dendritic segments from CA3 are shown for 6-month vehicle, 6-month VU0155069, 12-month vehicle, and 12-month VU0155069 groups. Endpoints were: (A) total spine number; (B) stubby-spine count; (C) filamentous-spine count; (D) thin-spine count; (E) mushroom-spine count; (F) spine density per 10 µm; (G) individual spine area; (H) total dendritic area; (I) spine perimeter; (J) dendrite major axis; (K) dendrite minor axis; (L) mushroom-spine head width; and (M) mushroom-spine neck length. Data are displayed as mean ± SEM. The animal was the biological replicate, with n = 5 animals per group. Endpoint-specific n values are provided in the Supplementary Information. At 6 months, treatment was associated with increased total spine number (p = 0.0404; panel A). At 12 months, treatment was associated with increased dendrite major axis (p = 0.0466; panel J) and dendrite minor axis (p = 0.0016; panel K). Only significant within-age comparisons are displayed above bars. Significant Kruskal–Wallis omnibus results across the four independent age-by-treatment groups are shown separately within the corresponding panels. Four-group effects were detected for total spine number, filamentous-spine count, thin-spine count, spine density, and mushroom-spine head width. Complete Mann–Whitney U statistics and exact p values, Kruskal–Wallis H statistics, and Dunn-adjusted comparisons are provided in the Supplementary Information (**Suppl. Tables 6A** and **6B**). The heatmap displays animal-level averages normalized to the 6-month vehicle mean for the corresponding endpoint, set to 100%. Purple indicates values at, below, or near the reference; green indicates values above the reference; and grey indicates a true numerical zero rather than missing data. Each heatmap column represents one animal. The heatmap is descriptive and was not used for inferential analysis. Veh, vehicle; VU01, VU0155069.

At 6 months, VU0155069 treatment was associated with increased total spine number (p = 0.0404; **Fig. 6A**). No statistically significant within-age differences were detected for stubby-spine count (p > 0.9999; **Fig. 6B**), filamentous-spine count (p = 0.6877; **Fig. 6C**), thin-spine count (p = 0.1493; **Fig. 6D**), mushroom-spine count (p = 0.1744; **Fig. 6E**), spine density (p = 0.3441; **Fig. 6F**), individual spine area (p = 0.5925; **Fig. 6G**), total dendritic area (p = 0.3414; **Fig. 6H**), spine perimeter (p = 0.5779; **Fig. 6I**), dendrite major axis (p = 0.2227; **Fig. 6J**), dendrite minor axis (p = 0.7129; **Fig. 6K**), mushroom-spine head width (p = 0.0731; **Fig. 6L**), or mushroom-spine neck length (p = 0.4469; **Fig. 6M**).

At 12 months, VU0155069 treatment was associated with increased dendrite major axis (p = 0.0466; **Fig. 6J**) and dendrite minor axis (p = 0.0016; **Fig. 6K**). No statistically significant within-age differences were detected for total spine number (p = 0.2555; **Fig. 6A**), stubby-spine count (p = 0.7470; **Fig. 6B**), filamentous-spine count (p = 0.3060; **Fig. 6C**), thin-spine count (p = 0.1627; **Fig. 6D**), mushroom-spine count (p = 0.9502; **Fig. 6E**), spine density (p = 0.5644; **Fig. 6F**), individual spine area (p = 0.0511; **Fig. 6G**), total dendritic area (p = 0.4641; **Fig. 6H**), spine perimeter (p = 0.3989; **Fig. 6I**), mushroom-spine head width (p = 0.1265; **Fig. 6L**), or mushroom-spine neck length (p = 0.2508; **Fig. 6M**).

Four-group Kruskal–Wallis analyses identified overall differences in total spine number (p < 0.0001; **Fig. 6A**), spine density (p = 0.0008; **Fig. 6F**), total dendritic area (p = 0.0052; **Fig. 6H**), dendrite minor axis (p = 0.0164; **Fig. 6K**), and mushroom-spine head width (p = 0.0012; **Fig. 6L**). No significant omnibus differences were detected for stubby-spine count (p = 0.6250; Fig. **6B**), filamentous-spine count (p = 0.7243; **Fig. 6C**), thin-spine count (p = 0.0639; **Fig. 6D**), mushroom-spine count (p = 0.6017; **Fig. 6E**), individual spine area (p = 0.1483; **Fig. 6G**), spine perimeter (p = 0.7341; **Fig. 6I**), dendrite major axis (p = 0.1374; **Fig. 6J**), or mushroom-spine neck length (p = 0.2950; **Fig. 6M**).

The 12-month vehicle-versus-VU0155069 difference in dendrite minor axis remained significant after Dunn adjustment, whereas the corresponding dendrite-major-axis comparison did not. Complete mean-rank differences and adjusted p values are provided in the Supplementary Information (**Suppl. Tables 6A** and **6B**).

The CA3 findings reveal a selective, stage-dependent structural response to VU0155069 that evolves from increased spine abundance during earlier pathology to remodeling of individual-spine geometry at the later stage. At 6 months, treatment increased total spine number, indicating expansion of the available synaptic substrate. At 12 months, the response shifted toward increased dendrite major and minor axes without corresponding changes in total spine number, subtype distribution, or spine density, suggesting that VU0155069 preferentially remodeled the geometry of existing spine populations. Thus, in contrast to the broader and more multidimensional response observed in CA2, CA3 engaged a focused structural program in which early expansion of spine number was followed by later enlargement of individual spine profiles. This stage-dependent transition highlights complementary forms of PLD1-linked structural remodeling within the hippocampus, with early expansion of the available spine population followed by later remodeling of individual-spine geometry. This interpretation is supported by evidence that PLD1 signaling regulates both dendritic arborization and spine morphogenesis and provides a plausible structural correlate of the previously demonstrated improvement in CA3–CA1 synaptic plasticity following VU0155069 treatment[13,14,19,62].

### Dentate Gyrus (DG)

The effects of PLD1 attenuation were examined in the dentate gyrus (DG) region, which is imperative for pattern separation processes from the entorhinal cortex and episodic memory formation as well as discrimination [70]. Additionally, this region is uniquely involved in adult neurogenesis, where new neurons are continuously developed from stem cells in the adult brain benefiting cognitive functions. During the late stage of differentiation during neurogenesis, formation of dendritic spines occur [71]. However, the DG is the earliest region to be impacted in AD and prior studies using transgenic mouse models have shown detriments in neurogenesis functions. APPswe/PS1^ΔE9^ displaying classic AD pathology, exhibited pronounced impairments in neurogenesis, specifically in neuronal progenitor cell proliferation and differentiation at 2 months of age before Aβ plaque deposition [72]. As a result, reduction of neurogenesis has shown to have significantly negative synaptic effects on LTP and LTD responses [73]. Thus, DG was examined to determine whether repeated VU0155069 treatment was associated with dendritic spine remodeling in the hippocampal input region at the 6-and 12-month pathological stages using Golgi–Cox images acquired from archived specimens to quantify spine abundance, subtype distribution, density, and morphometric parameters in standardized 10-µm spiny dendritic segments (**Fig. 7**).

**Figure 7.**
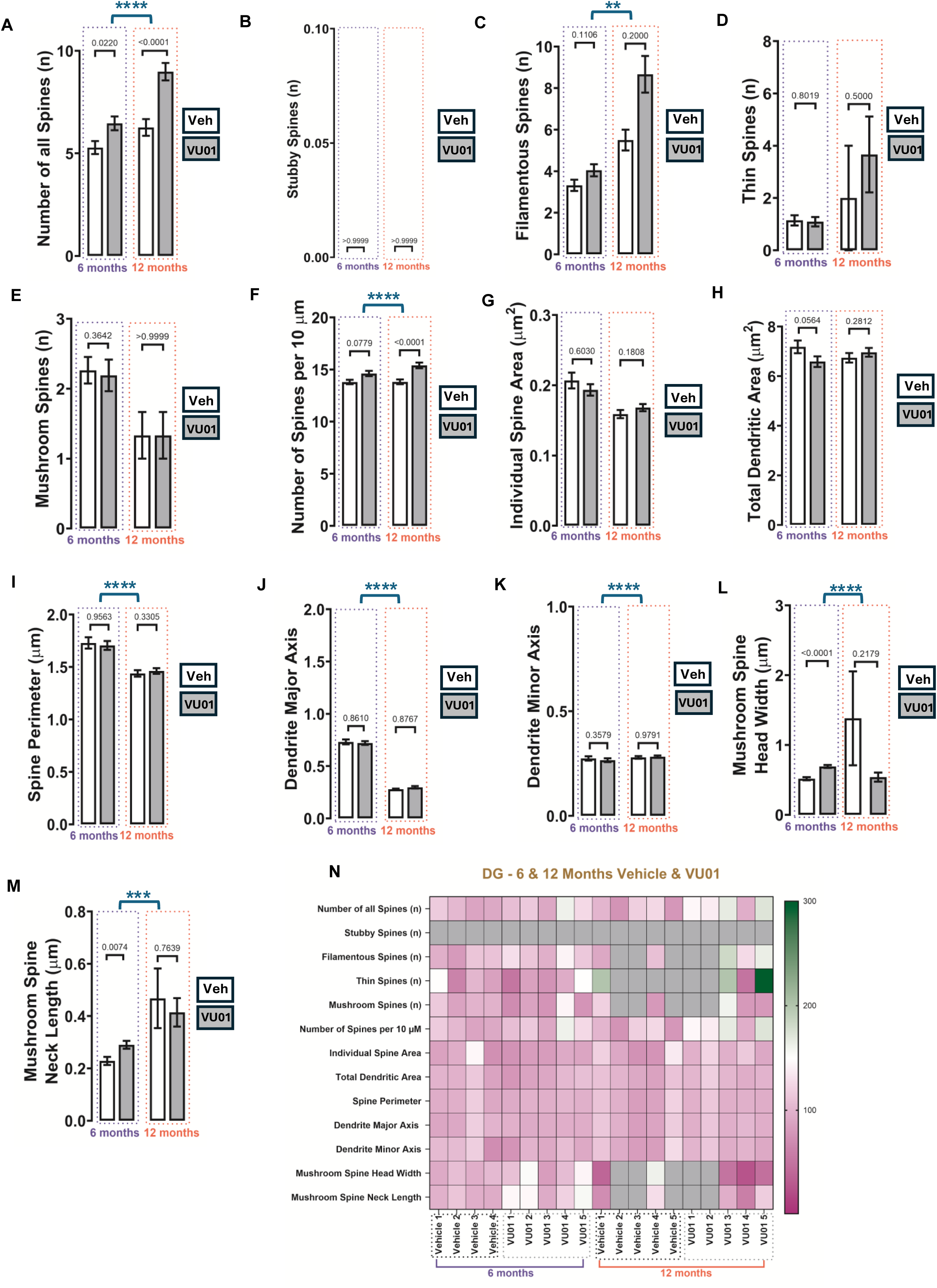
VU0155069-associated dendritic spine remodeling in the dentate gyrus (DG) of female 3xTg-AD mice. Representative Golgi–Cox images and quantitative analyses of standardized 10-µm spiny dendritic segments from DG are shown for 6-month vehicle, 6-month VU0155069, 12-month vehicle, and 12-month VU0155069 groups. Endpoints were: (A) total spine number; (B) stubby-spine count; (C) filamentous-spine count; (D) thin-spine count; (E) mushroom-spine count; (F) spine density per 10 µm; (G) individual spine area; (H) total dendritic area; (I) spine perimeter; (J) dendrite major axis; (K) dendrite minor axis; (L) mushroom-spine head width; and (M) mushroom-spine neck length. Data are displayed as mean ± SEM. n = 5 animals per group. At 6 months, treatment was associated with increased total spine number. At 12 months, treatment was associated with increased total spine number (p < 0.0001; panel A) and spine density (p < 0.0001; panel F). Only significant within-age comparisons are displayed above bars; significant Significant Kruskal–Wallis omnibus results across the four independent age-by-treatment groups are shown separately within the corresponding panels. Four-group effects were detected for total spine number, filamentous-spine count, thin-spine count, spine density, and mushroom-spine head width. Complete Mann–Whitney U statistics and exact p values, Kruskal– Wallis H statistics, and Dunn-adjusted comparisons are provided in the Supplementary Information (**Suppl. Tables 7A** and **7B**). The heatmap displays animal-level averages normalized to the 6-month vehicle mean for the corresponding endpoint, set to 100%. Purple indicates values at, below, or near the reference; green indicates values above the reference; and grey indicates a true numerical zero rather than missing data. Each heatmap column represents one animal. The heatmap is descriptive and was not used for inferential analysis. Veh, vehicle; VU01, VU0155069.

At 6 months, VU0155069 treatment was associated with increased total spine number (p = 0.0220; **Fig. 7A**), mushroom-spine head width (p < 0.0001; **Fig. 7L**), and mushroom-spine neck length (p = 0.0074; **Fig. 7M**). No statistically significant within-age differences were detected for stubby-spine count (p > 0.9999; **Fig. 7B**), filamentous-spine count (p = 0.1106; **Fig. 7C**), thin-spine count (p = 0.8019; **Fig. 7D**), mushroom-spine count (p = 0.3642; **Fig. 7E**), spine density (p = 0.0779; **Fig. 7F**), individual spine area (p = 0.6030; **Fig. 7G**), total dendritic area (p = 0.0564; **Fig. 7H**), spine perimeter (p = 0.9563; **Fig. 7I**), dendrite major axis (p = 0.8610; **Fig. 7J**), and dendrite minor axis (p = 0.3579; **Fig. 7K**).

At 12 months, VU0155069 treatment was associated with increased total spine number (p < 0.0001; **Fig. 7A**) and spine density (p < 0.0001; **Fig. 7F**). No statistically significant within-age differences were detected for stubby-spine count (p > 0.9999; **Fig. 7B**), filamentous-spine count (p = 0.2000; **Fig. 7C**), thin-spine count (p = 0.5000; **Fig. 7D**), mushroom-spine count (p > 0.9999; **Fig. 7E**), individual spine area (p = 0.1808; **Fig. 7G**), total dendritic area (p = 0.2812; **Fig. 7H**), spine perimeter (p = 0.3305; **Fig. 7I**), dendrite major axis (p = 0.8767; **Fig. 7J**), dendrite minor axis (p = 0.9791; **Fig. 7K**), mushroom-spine head width (p = 0.2179; **Fig. 7L**), or mushroom-spine neck length (p = 0.7639; **Fig. 7M**).

Four-group Kruskal–Wallis analyses identified overall differences in total spine number (p < 0.0001; **Fig. 7A**), filamentous-spine count (p = 0.0051; **Fig. 7C**), spine density (p < 0.0001; **Fig. 7F**), spine perimeter (p < 0.0001; **Fig. 7I**), dendrite major axis (p < 0.0001; **Fig. 7J**), dendrite minor axis (p < 0.0001; **Fig. 7K**), mushroom spine head width (p < 0.0001; **Fig. 7L**) and mushroom-spine neck length (p = 0.0003; **Fig. 7M**). No significant omnibus differences were detected for thin-spine count (p = 0.2147; **Fig. 7D**), mushroom-spine count (p = 0.2993; **Fig. 7E**), individual spine area (p = 0.1785; **Fig. 7G**), or total dendritic area (p = 0.0910; **Fig. 7H**). No Kruskal-Wallis statistics could be calculated for stubby-spine count as all observations were zero across group, leading to no variance (**Fig. 7B**). Complete mean-rank differences and adjusted p values are provided in the Supplementary Information (**Suppl. Tables 7A** and **7B**).

Together, the DG findings reveal a sustained and stage-amplified expansion of the structural substrate available for hippocampal input processing. VU0155069 increased total spine number, mushroom spine head width and neck length at 6 months. However, it only produced a coordinated increase in total spine number and spine density at 12 months, while maintaining a comparatively stable distribution of spine phenotypes and individual-spine dimensions. This pattern distinguishes DG from CA2 and CA3 and suggests that PLD1 attenuation strengthens DG circuitry through an early expansion in synaptic capacity and reshaping of mushroom spines, followed by more selective strengthening of specific synaptic circuits at later stages [74]. Because the DG continuously integrates adult-born granule neurons that acquire dendritic spines as they mature and enter existing hippocampal networks [75], the abundance-centered response is compatible with enhanced neuronal integration, maintenance of newly established synapses, or preservation of mature granule-cell connectivity [76]. PLD1-linked signaling has also been implicated in neural-stem-cell differentiation and neurogenesis [77], providing a mechanistic basis for examining whether modulation of this pathway contributes to the DG response. Thus, the present findings identify sustained expansion of the DG spine population as a distinctive component of VU0155069-associated hippocampal resilience, while establishing adult-born granule-cell integration [78] as a focused and biologically grounded mechanism for future investigation.

### Central Amygdala (CeA)

The central amygdala (CeA) is essential for conditional freezing after Pavlovian fear conditioning [79], where overtraining may engage a compensatory fear conditioning circuit involving the CeA in animals with damage to the basolateral amygdala (BLA). Moreover, our own studies identified PLD1 as a convergent second messenger affecting retrieval of appetitive or aversive memories in normal rats [9–11] in the amygdala. Moreover, amygdala is affected early in neurodegenerative disease [45–47,80] making our determination of how repeated VU0155069 treatment participates in dendritic spine remodeling in this amygdalar nuclei at the 6-and 12-month pathological stages using Golgi–Cox images acquired from archived specimens and were used to quantify spine abundance, subtype distribution, density, and morphometric parameters in standardized 10-µm spiny dendritic segments (**Fig. 8**).

**Figure 8.**
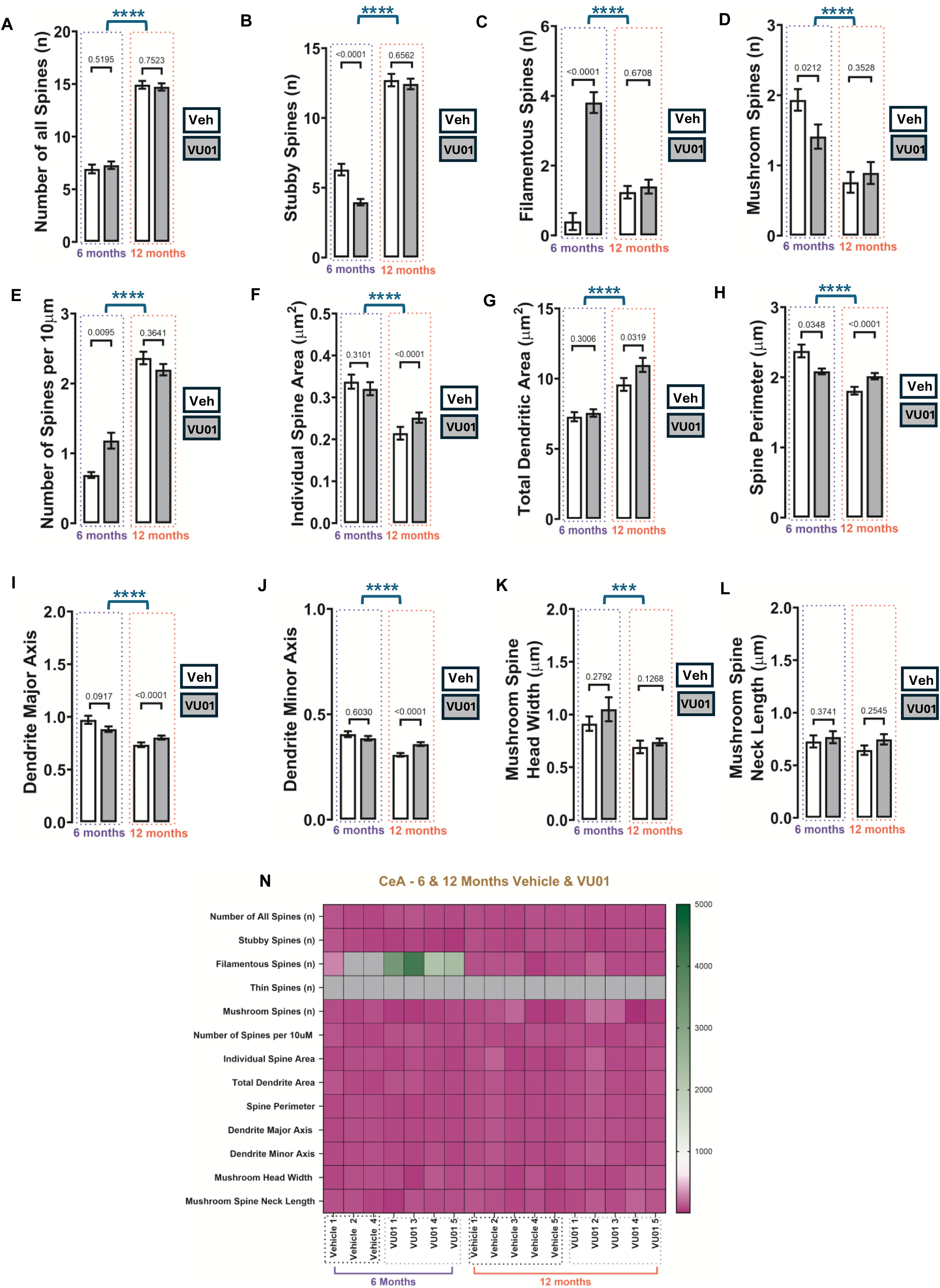
VU0155069-associated dendritic spine remodeling in the central amygdala (CeA) of female 3xTg-AD mice. Representative Golgi–Cox images and quantitative analyses of standardized 10-µm spiny dendritic segments from CeA are shown for 6-month vehicle, 6-month VU0155069, 12-month vehicle, and 12-month VU0155069 groups. Endpoints were: (A) total spine number; (B) stubby-spine count; (C) filamentous-spine count; (D) mushroom-spine count; (E) spine density per 10 µm; (F) individual spine area; (G) total dendritic area; (H) spine perimeter; (I) dendrite major axis; (J) dendrite minor axis; (K) mushroom-spine head width; and (L) mushroom-spine neck length. Thin spines were not detected and were not analyzed separately. Data are displayed as mean ± SEM and n = 5 animals per group. Only significant within-age comparisons are displayed above bars; significant Kruskal–Wallis omnibus results are shown separately. Significant Kruskal–Wallis omnibus results across the four independent age-by-treatment groups are shown separately within the corresponding panels. Four-group effects were detected for total spine number, filamentous-spine count, thin-spine count, spine density, and mushroom-spine head width. Complete Mann–Whitney U statistics and exact p values, Kruskal– Wallis H statistics, and Dunn-adjusted comparisons are provided in the Supplementary Information (**Suppl. Tables 8A** and **8B**). The heatmap displays animal-level averages normalized to the 6-month vehicle mean for the corresponding endpoint, set to 100%. Purple indicates values at, below, or near the reference; green indicates values above the reference; and grey indicates a true numerical zero rather than missing data. Each heatmap column represents one animal. The heatmap is descriptive and was not used for inferential analysis. Veh, vehicle; VU01, VU0155069.

Thin spines were not detected in the sampled CeA segments and therefore were not analyzed as a separate endpoint. Primary comparisons evaluated vehicle-and VU0155069-treated mice within each age cohort using two-sided Mann-Whitney U tests. Four-group Kruskal-Wallis analyses followed by Dunn-adjusted comparisons provided additional age-cohort context.

At 6 months, VU0155069 treatment was associated with reduced stubby-spine count (p < 0.0001; **Fig. 8B**), mushroom-spine count (p = 0.0212; **Fig. 8D**), and spine perimeter (p = 0.0348; **Fig. 8H**), together with increased filamentous-spine count (p < 0.0001; **Fig. 8C**) and spine density (p = 0.0095; **Fig. 8E**). No statistically significant within-age differences were detected for total spine number (p = 0.5195; **Fig. 8A**), individual spine area (p = 0.3101; **Fig. 8F**), total dendritic area (p = 0.3006; **Fig. 8G**), dendrite major axis (p = 0.0917; **Fig. 8I**), dendrite minor axis (p = 0.6030; **Fig. 8J**), mushroom-spine head width (p = 0.2792; **Fig. 8K**), or mushroom-spine neck length (p = 0.3741; **Fig. 8L**).

At 12 months, VU0155069 treatment was associated with increased individual spine area (p < 0.0001; **Fig. 8F**), total dendritic area (p = 0.0319; **Fig. 8G**), spine perimeter (p < 0.0001; **Fig. 8H**), dendrite major axis (p < 0.0001; **Fig. 8I**), and dendrite minor axis (p < 0.0001; **Fig. 8J**). No statistically significant within-age differences were detected for total spine number (p = 0.7523; **Fig. 8A**), stubby-spine count (p = 0.6562; **Fig. 8B**), filamentous-spine count (p = 0.6708; **Fig. 8C**), mushroom-spine count (p = 0.3528; **Fig. 8D**), spine density (p = 0.3641; **Fig. 8E**), mushroom-spine head width (p = 0.1268; **Fig. 8K**), or mushroom-spine neck length (p = 0.2545; **Fig. 8L**).

Four-group Kruskal-Wallis analyses identified overall differences in all 11 measured CeA endpoints (all p ≤ 0.0003; **Fig. 8A–K**) with the exception of mushroom spine neck length (**Fig. 8L**). Dunn-adjusted comparisons indicated extensive separation between the independently processed 6-and 12-month cohorts. The 12-month vehicle-versus-VU0155069 differences in individual spine area, spine perimeter, dendrite major axis, and dendrite minor axis remained significant after adjustment. At 6 months, the direct filamentous-spine contrast remained significant after Dunn adjustment, whereas the stubby-and mushroom-spine contrasts did not. The significant difference in spine density at 6 months was maintained following Dunn’s adjustment. Complete mean-rank differences and adjusted p values are provided in the Supplementary Information (**Suppl. Tables 8A** and **8B**).

Together, the CeA findings identify a stage-dependent remodeling program that evolves from early reorganization of spine phenotypes to later enlargement of the available synaptic structures. At 6 months, VU0155069 altered the balance among stubby, filamentous, and mushroom profiles, increased spine density, and modified spine perimeter. At 12 months, treatment increased individual-spine area, perimeter, major and minor axes, and total dendritic area while maintaining overall spine number and subtype abundance. This transition suggests that PLD1 attenuation initially increases structural flexibility within CeA and subsequently promotes consolidation of the spine population as pathology advances.

Across fear-and cocaine-conditioning paradigms, PLD emerged as a common downstream signaling node linking metabotropic glutamate, D1/D5 dopamine, and 5-HT_2C_ serotonin receptor systems to amygdalar synaptic plasticity and associative-memory expression [9–11]. Fear-associated memory recruited PLD-linked metabotropic glutamate receptor signaling within the thalamic-to-lateral amygdala pathway [10], while cocaine-conditioned appetitive memory engaged interactions among D1/D5 dopamine receptors, group I metabotropic glutamate receptors, and PLD within the BLA-to-CeA circuit [9]. Subsequent work extended this convergence to 5-HT_2C_ receptor signaling in cocaine-associated amygdala-hippocampal adaptations [11]. The present CeA remodeling profile, therefore, adds a structural dimension to a signaling framework developed across more than a decade of work, positioning PLD1 as a convergent regulator through which glutamatergic, dopaminergic, and serotonergic inputs may shape amygdalar plasticity in both associative-memory and neurodegenerative contexts.

### Lateral Amygdala (LA)

The lateral amygdala (LA) is one of the nine nuclei of the amygdala complex that functions as a sensory input capable of storing associative cued memory of learned fear is involved with learning associated with the CeA [81,82]. It is capable of a mix of excitatory and inhibitory neurotransmission [83]. Lesions or injuries in the LA region result in the incapacity to learn or acquire any fear-associated memories [84]. The LA integrates sensory inputs for fear learning and impinges on cued input to the CeA that complements the contextual input from the BLA. It serves as a key input node for emotional learning as well as memory and, together with the broader amygdalar circuitry, represents a region in which PLD1-linked signaling may contribute to long-term synaptic adaptations [85]. LA was examined to determine whether repeated VU0155069 treatment was associated with dendritic spine remodeling in an amygdalar sensory-input nucleus at the 6-and 12-month pathological stages using Golgi–Cox images acquired from archived specimens where spine abundance, subtype distribution, density, and morphometric parameters in standardized 10-µm spiny dendritic segments were quantified to assess changes (**Fig. 9**). Thin spines were not detected in the sampled LA segments and therefore were not analyzed as a separate endpoint.

**Figure 9.**
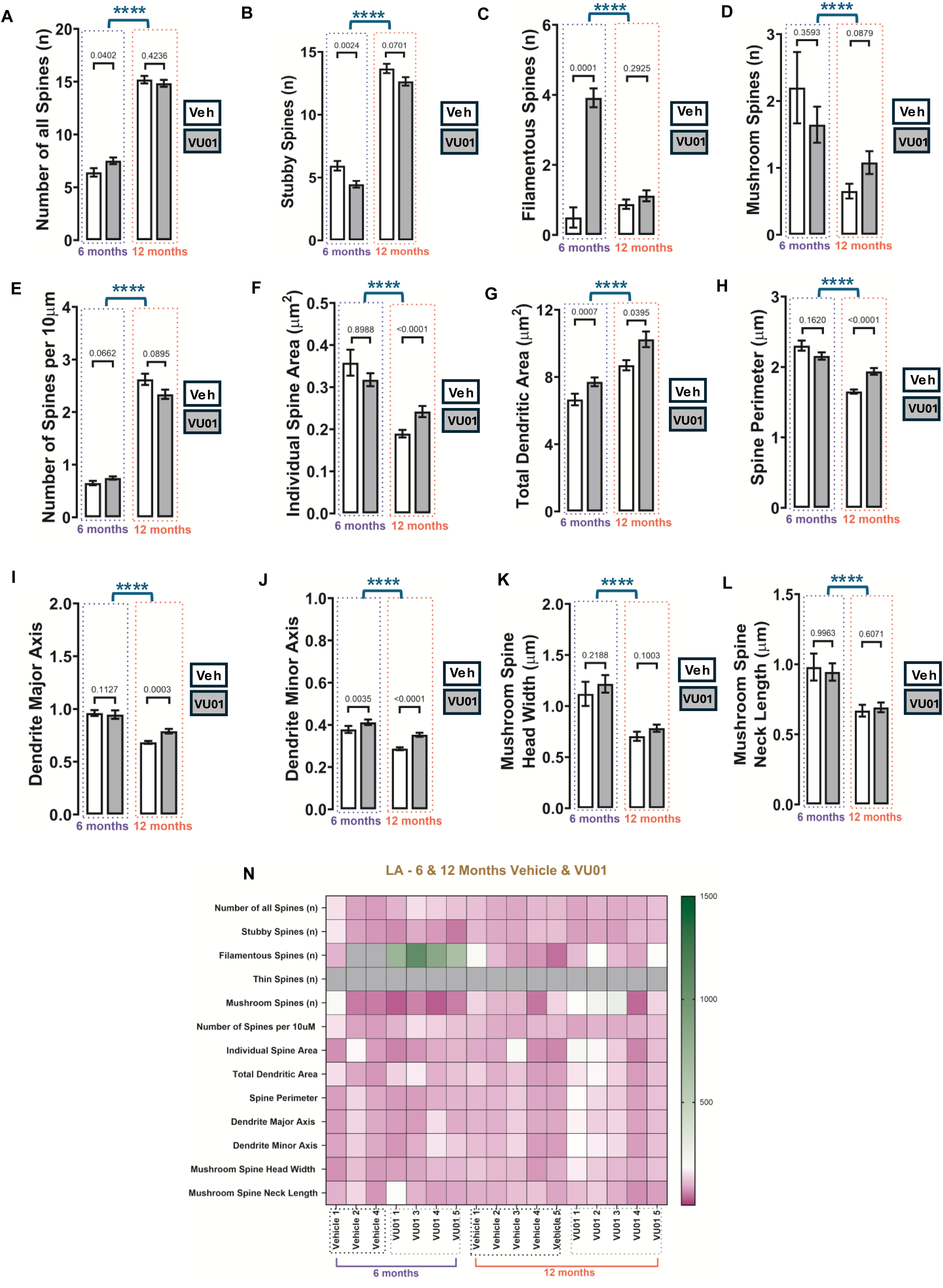
VU0155069-associated dendritic spine remodeling in the lateral amygdala (LA) of female 3xTg-AD mice. Representative Golgi–Cox images and quantitative analyses of standardized 10-µm spiny dendritic segments from LA are shown for 6-month vehicle, 6-month VU0155069, 12-month vehicle, and 12-month VU0155069 groups. Endpoints were: (A) total spine number; (B) stubby-spine count; (C) filamentous-spine count; (D) mushroom-spine count; (E) spine density per 10 µm; (F) individual spine area; (G) total dendritic area; (H) spine perimeter; (I) dendrite major axis; (J) dendrite minor axis; (K) mushroom-spine head width; and (L) mushroom-spine neck length. Thin spines were not detected and were not analyzed separately. Data are displayed as mean ± SEM. The animal was the biological replicate, with n = 5 animals per group. At 6 months, treatment was associated with increased total spine number (p = 0.0402; panel A), filamentous-spine count (p = 0.0001; panel C), total dendritic area (p = 0.0007; panel G), and dendrite minor axis (p = 0.0035; panel J) and with reduced stubby-spine count (p = 0.0024; panel B). At 12 months, treatment was associated with increased individual spine area (p < 0.0001; panel F), total dendritic area (p = 0.0395; panel G), spine perimeter (p < 0.0001; panel H), dendrite major axis (p m= 0.0003; panel I), and dendrite minor axis (p < 0.0001; panel J). Only significant within-age comparisons are displayed above bars; significant Kruskal–Wallis omnibus results are shown separately. Significant Kruskal–Wallis omnibus results across the four independent age-by-treatment groups are shown separately within the corresponding panels. Four-group effects were detected for total spine number, filamentous-spine count, thin-spine count, spine density, and mushroom-spine head width. Complete Mann–Whitney U statistics and exact p values, Kruskal– Wallis H statistics, and Dunn-adjusted comparisons are provided in the Supplementary Information (**Suppl. Tables 9A** and **9B**). The heatmap displays animal-level averages normalized to the 6-month vehicle mean for the corresponding endpoint, set to 100%. Purple indicates values at, below, or near the reference; green indicates values above the reference; and grey indicates a true numerical zero rather than missing data. Each heatmap column represents one animal. The heatmap is descriptive and was not used for inferential analysis. Veh, vehicle; VU01, VU0155069.

At 6 months, VU0155069 treatment was associated with increased total spine number (p = 0.0402; **Fig. 9A**), filamentous-spine count (p = 0.0001; **Fig. 9C**), total dendritic area (p = 0.0007; **Fig. 9G**), and dendrite minor axis (p = 0.0035; **Fig. 9J**). In contrast, stubby-spine count was reduced (p = 0.0024; **Fig. 9B**). No statistically significant within-age differences were detected for mushroom-spine count (p = 0.3593; **Fig. 9D**), spine density (p = 0.0662; **Fig. 9E**), individual spine area (p = 0.8988; **Fig. 9F**), spine perimeter (p = 0.1620; **Fig. 9H**), dendrite major axis (p = 0.1127; **Fig. 9I**), mushroom-spine head width (p = 0.2188; **Fig. 9K**), or mushroom-spine neck length (p = 0.9963; **Fig. 9L**).

At 12 months, VU0155069 treatment was associated with increased individual spine area (p < 0.0001; **Fig. 9F**), total dendritic area (p = 0.0359; **Fig. 9G**), spine perimeter (p < 0.0001; **Fig. 9H**), dendrite major axis (p = 0.0003; **Fig. 9I**), and dendrite minor axis (p < 0.0001; **Fig. 9J**). No statistically significant within-age differences were detected for total spine number (p = 0.4236; **Fig. 9A**), stubby-spine count (p = 0.0701; **Fig. 9B**), filamentous-spine count (p = 0.2925; **Fig. 9C**), mushroom-spine count (p = 0.0879; **Fig. 9D**), spine density (p = 0.0895; **Fig. 9E**), mushroom-spine head width (p = 0.1003; **Fig. 9K**), or mushroom-spine neck length (p = 0.6071; **Fig. 9L**).

Four-group Kruskal-Wallis analyses identified overall differences in all twelve measured LA endpoints (all p < 0.0001; **Fig. 9A–L**). Dunn-adjusted comparisons indicated extensive separation between the independently processed 6-and 12-month cohorts. At 6 months, the direct filamentous-spine and total-dendritic-area contrasts remained significant after adjustment, whereas the total spine, stubby spine, and dendrite-minor-axis contrasts did not. At 12 months, the vehicle-versus-VU0155069 differences in individual spine area, spine perimeter, dendrite major axis, and dendrite minor axis remained significant after adjustment; the total-dendritic-area contrast did not. Complete mean-rank differences and adjusted p values are provided in the Supplementary Information (**Suppl. Tables 9A** and **9B**).

Together, the LA findings reveal a robust, stage-dependent structural response to VU0155069 that progresses from early expansion and reorganization of the spine population to later morphometric consolidation. At 6 months, treatment increased total spine number and filamentous-spine abundance, reduced stubby-spine abundance, and increased total dendritic area and dendrite minor axis. This coordinated pattern suggests that PLD1 attenuation expands the available synaptic substrate while shifting the LA spine population toward elongated profiles associated with structural plasticity. At 12 months, the response transitioned toward broader morphometric enlargement, including increased individual-spine area, total dendritic area, spine perimeter, and dendrite major and minor axes, while total spine number and subtype abundance remained comparatively stable. The later profile therefore suggests that VU0155069 supports remodeling and consolidation of the established spine population rather than continued expansion of spine number.

This stage-dependent progression is particularly relevant to the function of the LA as a major site of sensory-associative plasticity. Previous studies have demonstrated that fear learning modifies synaptic strength, synapse size, and local dendritic translation in LA, establishing a close relationship between LA structural plasticity and the encoding of emotionally salient associations [86]. The present findings suggest that attenuation of aberrant PLD1 signaling preserves this structural adaptability during earlier pathology and supports enlargement of the available synaptic structures as pathology advances.

The LA profile also extends our earlier identification of PLD as a convergent signaling mechanism in amygdala-dependent memory [9–11]. Viewed within this established framework, the early LA expansion and redistribution of spine profiles may increase the structural repertoire available for associative plasticity, whereas the later enlargement of individual-spine profiles may strengthen or stabilize the synaptic substrate supporting previously encoded associations. Thus, the LA findings add a structural dimension to the convergent PLD-signaling model and provide a plausible anatomical basis for the preservation of fear-conditioning performance observed after VU0155069 treatment in the earlier and later 3xTg-AD cohorts[13,14]. Together with the CeA findings, the data suggests a coordinated amygdalar resilience program in which the sensory-associative input nucleus and downstream output circuitry engage complementary, stage-dependent forms of spine remodeling.

### Basolateral Amygdala (BLA)

The BLA region is an important factor in determining memory consolidation and associative memory [87] and is also highly vulnerable and critical for emotional memory components (such as frustration, anxiety) that is impinged at early stages in neurodegenerative states [88,89]. The BLA is a major integrative node for emotional learning as well as memory and, based on prior work from our group, represents a region in which PLD1-linked signaling may contribute to long-term synaptic adaptations across associative circuitry [9–11]. Therefore, we examined repeated VU0155069 treatment on dendritic spine remodeling in this amygdalar integrative nucleus at the 6-and 12-month pathological stages using Golgi-Cox images acquired from archived specimens that were used to quantify spine abundance, subtype distribution, density, and morphometric parameters in standardized 10-µm spiny dendritic segments (**Fig. 10**). Thin spines were not detected in the sampled BLA segments and therefore were not analyzed separately.

**Figure 10.**
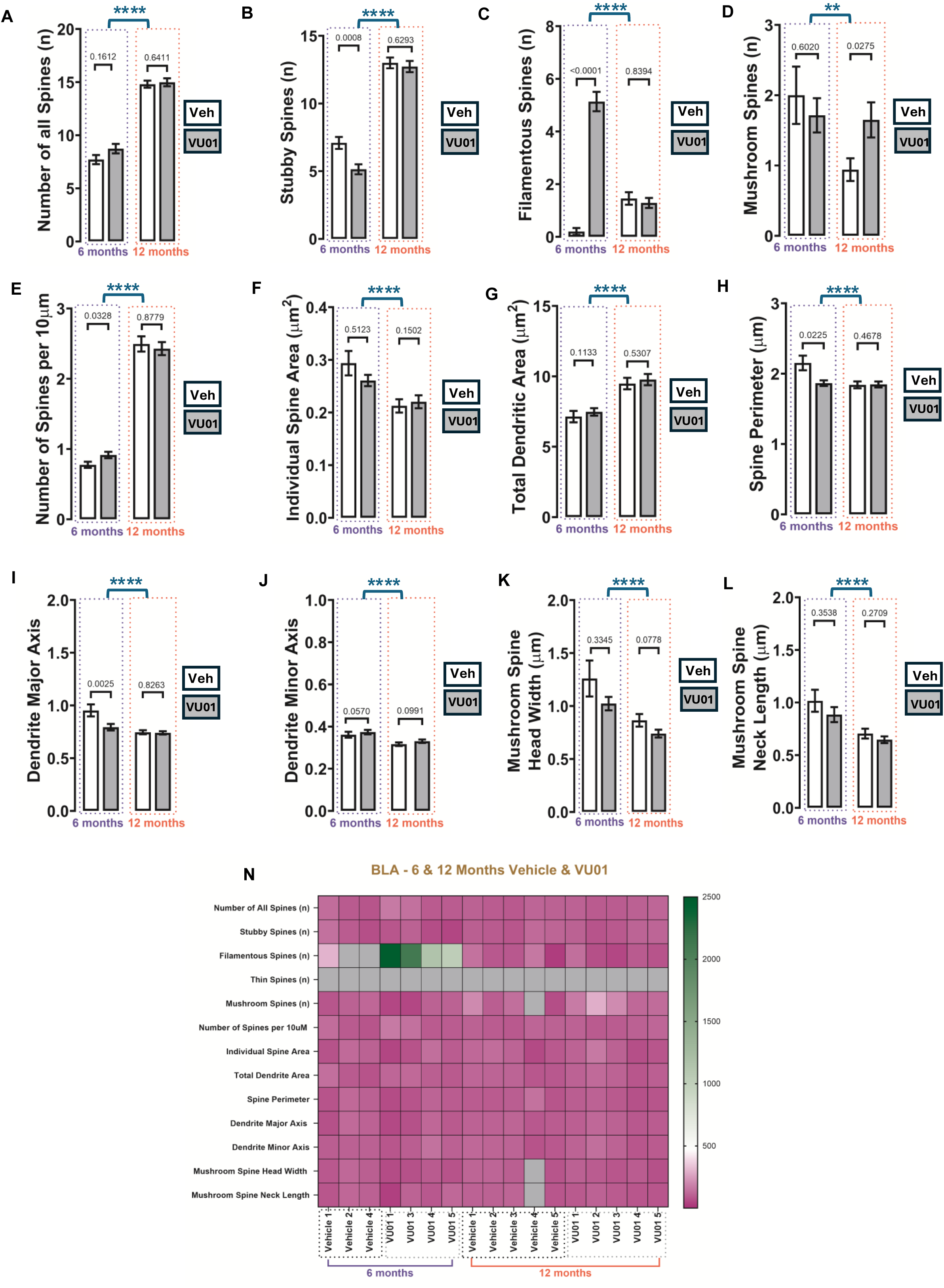
VU0155069-associated dendritic spine remodeling in the basolateral amygdala (BLA) of female 3xTg-AD mice. Representative Golgi–Cox images and quantitative analyses of standardized 10-µm spiny dendritic segments from BLA are shown for 6-month vehicle, 6-month VU0155069, 12-month vehicle, and 12-month VU0155069 groups. Endpoints were: (A) total spine number; (B) stubby-spine count; (C) filamentous-spine count; (D) mushroom-spine count; (E) spine density per 10 µm; (F) individual spine area; (G) total dendritic area; (H) spine perimeter; (I) dendrite major axis; (J) dendrite minor axis; (K) mushroom-spine head width; and (L) mushroom-spine neck length. Thin spines were not detected and were not analyzed separately. Data are displayed as mean ± SEM with n = 5 animals per group. At 6 months, treatment was associated with increased filamentous-spine count (p < 0.0001; panel C) and spine density (p = 0.0328; panel E) and with reduced stubby-spine count (p = 0.0008; panel B), spine perimeter (p = 0.0225; panel H), and dendrite major axis (p = 0.0025; panel I). At 12 months, treatment was associated only with increased mushroom-spine count (p = 0.0275; panel D). Only significant within-age comparisons are displayed above bars; significant omnibus results are shown separately. Significant Kruskal–Wallis omnibus results across the four independent age-by-treatment groups are shown separately within the corresponding panels. Four-group effects were detected for total spine number, filamentous-spine count, thin-spine count, spine density, and mushroom-spine head width. Complete Mann–Whitney U statistics and exact p values, Kruskal–Wallis H statistics, and Dunn-adjusted comparisons are provided in the Supplementary Information (**Suppl. Tables 10A** and **10B**). The heatmap displays animal-level averages normalized to the 6-month vehicle mean for the corresponding endpoint, set to 100%. Purple indicates values at, below, or near the reference; green indicates values above the reference; and grey indicates a true numerical zero rather than missing data. Each heatmap column represents one animal. The heatmap is descriptive and was not used for inferential analysis. Veh, vehicle; VU01, VU0155069.

At 6 months, VU0155069 treatment was associated with increased filamentous-spine count (p < 0.0001; **Fig. 10C**) and spine density (p = 0.0328; **Fig. 10E**), together with reduced stubby-spine count (p = 0.0008; **Fig. 10B**), spine perimeter (p = 0.0225; **Fig. 10H**), and dendrite major axis (p = 0.0025; **Fig. 10I**). No significant within-age differences were detected for total spine number (p = 0.1612; **Fig. 10A**), mushroom-spine count (p = 0.6020; **Fig. 10D**), individual spine area (p = 0.5123; **Fig. 10F**), total dendritic area (p = 0.1133; **Fig. 10G**), dendrite minor axis (p = 0.0570; **Fig. 10J**), mushroom-spine head width (p = 0.3345; **Fig. 10K**), or mushroom-spine neck length (p = 0.3538; **Fig. 10L**).

At 12 months, VU0155069 treatment was associated only with increased mushroom-spine count (p = 0.0275; **Fig. 10D**). No significant within-age differences were detected for total spine number (p = 0.6411; **Fig. 10A**), stubby-spine count (p = 0.6293; **Fig. 10B**), filamentous-spine count (p = 0.8394; **Fig. 10C**), spine density (p = 0.8779; **Fig. 10E**), individual spine area (p = 0.1502; **Fig. 10F**), total dendritic area (p = 0.5307; **Fig. 10G**), spine perimeter (p = 0.4678; **Fig. 10H**), dendrite major axis (p = 0.8263; **Fig. 10I**), dendrite minor axis (p = 0.0991; **Fig. 10J**), mushroom-spine head width (p = 0.0778; **Fig. 10K**), or mushroom-spine neck length (p = 0.2709; **Fig. 10L**).

Four-group analyses identified broad differences across BLA endpoints, but the overall structure was dominated by separation between the independently processed age cohorts. Dunn-adjusted comparisons retained the 6-month filamentous-spine difference and the 6-month spine-major-axis difference. No direct 12-month vehicle-versus-VU0155069 comparison remained significant after Dunn adjustment. Complete mean-rank differences and adjusted p values are provided in the Supplementary Information (**Suppl. Tables 10A** and **B**).

Together, the BLA findings reveal a precisely staged and phenotype-centered response to VU0155069. At 6 months, increased filamentous-spine abundance and spine density occurred together with reduced stubby-spine abundance, spine perimeter, and dendrite major axis, indicating early expansion and reorganization of a structurally adaptable spine population. At 12 months, the response shifted toward increased mushroom-spine abundance while total spine number, density, and broader morphometric measures remained stable, consistent with selective enrichment of an established spine phenotype. Thus, BLA contributes a complementary component of the amygdalar resilience program, progressing from early filamentous-spine enrichment to later mushroom-spine enrichment rather than reproducing the broader morphometric enlargement observed in CeA and LA.

## Discussion

### Principal findings and conceptual advance

The present study demonstrates that repeated VU0155069 treatment is associated with anatomically heterogeneous and pathological stage-dependent remodeling of dendritic spine architecture in female 3xTg-AD mice. The disease-matched comparator strategy is central to the therapeutic interpretation of these findings. PLD1 attenuation is intended to modify aberrant signaling within the affected state, and the direct pharmacological question is therefore whether VU0155069 changes synaptic architecture relative to age-matched vehicle-treated 3xTg-AD littermates. Wild-type cohorts provide an important physiological reference, but PLD1 is maintained near basal levels in healthy animals and the molecular organization of the wild-type brain differs substantially from that of the disease state. Consequently, a wild-type comparison would help determine whether an endpoint lies within the healthy structural range, whereas the present disease-state comparison determines whether the intervention recruits structural resilience within the population for which the therapy is intended.

Across ten anatomically defined cortical, hippocampal, and amygdalar regions, the direction and extent of the structural response varied according to brain region, age cohort, spine subtype, and morphometric endpoint. As expected, the findings, therefore, did not support a uniform structural effect of VU0155069 across the brain. Instead, the multiregional dataset identifies three recurring, nonexclusive patterns: (i) changes in spine abundance, (ii) redistribution among operational spine-morphology categories, and (iii) remodeling of two-dimensional spine or dendritic-segment morphometry.

These three complementary response patterns provide a systems-level framework for understanding how VU0155069 promotes structural resilience across anatomically and functionally specialized neural circuits. Regional differentiation was evident at both pathological stages and emerged as a principal biological finding of the study. A2 and CA2 engaged broad multidimensional remodeling programs, whereas A1, CA3, DG, and BLA recruited focused responses involving selected components of spine abundance, phenotype distribution, or geometry. The dominant form of remodeling also evolved with pathological stage. The earlier amygdalar response was characterized largely by redistribution among stubby, filamentous, and mushroom-spine profiles, while the later response in CeA and LA shifted toward enlargement of individual-spine and dendritic-segment measures. Within the visual cortex, V1 and V2 both expanded spine abundance during later-stage pathology but achieved this through distinct subtype and morphometric configurations. These coordinated differences could be resolved only through the present study approach of a comprehensive assessment of multiple spine parameters across ten anatomically defined regions.

The findings demonstrate that PLD1 attenuation supports distributed synaptic resilience by engaging the structural program most relevant to each circuit and pathological stage, thereby extending the previously established behavioral and physiological efficacy of VU0155069 into a detailed anatomical framework. Future regionally matched functional studies can now build on this framework to determine how these complementary structural programs support circuit plasticity and memory.

The inclusion of 6-and 12-month cohorts enabled the structural effects of VU0155069 to be examined across two previously characterized stages of amyloid-and tau-associated pathology in the 3xTg-AD model and in our established colony[13,14,28–30]. Within each stage, vehicle-and VU0155069-treated animals were evaluated as age-matched cohorts using consistent processing and analytical procedures, providing a rigorous basis for defining the treatment-associated response. This design revealed that PLD1 attenuation engages structural plasticity at both stages, while the predominant form of remodeling evolves with disease progression. The 6-month cohorts identified early changes in spine abundance and phenotype distribution, whereas the 12-month cohorts revealed continued structural responsiveness through region-specific changes in spine abundance and morphometry. Thus, the two-stage design extends the previously demonstrated functional efficacy of VU0155069 and shows that the therapeutic response is not restricted to an early intervention window but remains evident after AD-like pathology has progressed.

The present findings add an important structural dimension to the therapeutic efficacy of VU0155069 demonstrated previously through behavioral, electrophysiological, biochemical, and neuropathological outcomes **[13,14]**. Across the ten regions examined, PLD1 attenuation increased spine abundance, reorganized spine-phenotype distributions, and remodeled individual-spine and dendritic-segment morphometry in patterns that were specific to each circuit and pathological stage. Structural resilience was therefore not expressed as a uniform directional change across all endpoints. Instead, VU0155069 engaged complementary forms of structural plasticity, including expansion of the spine population in V1 and DG, multidimensional remodeling in A2 and CA2, and stage-dependent phenotype redistribution and morphometric consolidation across the amygdalar nuclei. This regional coordination provides a systems-level anatomical framework for the improvements in synaptic plasticity and memory reported previously and supports PLD1 attenuation as a therapeutic strategy capable of preserving and reorganizing the synaptic substrate across progressive stages of AD-like pathology. Inclusion of non-transgenic reference groups in future studies may further define how each regional response relates to the physiological range of the healthy brain. Nevertheless, because the therapeutic strategy is directed toward aberrantly elevated PLD1 signaling in the disease state, its pharmacological significance is most appropriately evaluated through comparison with the affected population receiving vehicle treatment[12].

### Relationship to previous PLD1 and VU0155069 studies

The present study extends previous PLD1 and VU0155069 findings in several ways. It evaluates multiple sensory, hippocampal, and amygdalar regions within a common treatment and imaging framework; examines spine abundance, operational phenotype distribution, and two-dimensional morphometry rather than density alone; and expands the structural analysis beyond the previously reported CA1 outcomes. This design provides a systems-level view of PLD1-sensitive structural remodeling across circuits relevant to sensory processing, contextual memory, and emotionally salient associative learning.

The current findings build directly on a body of work identifying aberrant post developmental PLD1 signaling as a contributor to AD-related synaptic vulnerability. Elevated PLD1 in synaptic fractions from human AD brain, together with experimental evidence linking PLD1 to Aβ-and tau-associated synaptic dysfunction, established the rationale for attenuating PLD1 as a strategy to preserve synaptic function[12–15,90]. In female 3xTg-AD mice, repeated VU0155069 treatment improved recognition and associative-memory performance, strengthened hippocampal synaptic plasticity, and preserved selected CA1 spine characteristics at an earlier disease stage[13]. Subsequent studies demonstrated that the intervention retained efficacy at 12 months, improving perirhinal cortex-, hippocampal-, and amygdala-dependent behaviors, HFS-LTP and LFS-LTD, selected biochemical and neuropathological measures, and CA1 spine morphology[14].

The present study advances this work from a selected CA1 analysis to a systems-level assessment of synaptic architecture. New imaging across ten additional regions demonstrates that the influence of VU0155069 extends through sensory cortices, hippocampal subfields, and amygdalar nuclei that collectively support recognition, contextual, associative, and emotional memory. The regional spine findings also complement the broader brain-region physiological heterogeneity identified by our group looking at glutamatergic synaptic integrity using fluorescence assisted single synaptosome long term potentiation (FASS-LTP) studies, which identified differential physiological effects in frontal cortex, parietal cortex, and midbrain preparations[15].

This interpretation does not require every region to exhibit the same directional change. Synaptic resilience may be achieved through different structural solutions in different circuits. In one region, resilience may be associated with expansion of spine abundance; in another, with redistribution toward spine profiles capable of further plasticity; and in another, with remodeling of spine dimensions or dendritic-segment area. The present multiregional analysis therefore moves beyond a simple “more spines is better” model and identifies PLD1-sensitive structural programs that differ across functionally specialized circuits.

The new dataset is also distinct from the previously published CA1 work. The treatment cohorts and selected CA1 findings were reported earlier, but the images, regional measurements, statistical analyses, and figures for A1, A2, V1, V2, CA2, CA3, DG, CeA, LA, and BLA are new. The use of established cohorts is an important strength because the structural observations can be interpreted within a treatment paradigm already supported by behavioral and physiological efficacy, while the new regional analysis adds anatomical resolution that was not available in the earlier studies.

Therefore, the previous therapeutic implications are strengthened by the continuity of the experimental program. VU0155069-associated resilience was established previously through improved novel-object recognition, fear-conditioning performance, hippocampal LTP, later-stage LTP and LTD, selected neuropathological outcomes, and CA1 spine integrity. The current findings now extend that evidence across visual and auditory cortices involved in processing conditioned and recognition-relevant sensory information, hippocampal subfields involved in contextual and relational memory, and amygdalar nuclei that encode and express emotionally salient associations.

The structural profiles reported here therefore provide a systems-level anatomical representation of therapeutic efficacy already demonstrated functionally at both pathological stages.

### Sensory cortices reveal circuit-dependent remodeling across disease stages

The auditory and visual cortical findings demonstrate that the structural response to VU0155069 is shaped by the functional organization of each sensory circuit. Primary sensory fields must preserve stable representations of fundamental stimulus features while remaining sufficiently plastic to adapt to changing input. Downstream and secondary fields transform those representations through broader contextual, associative, and integrative processing. Accordingly, synaptic plasticity is not expressed uniformly across the sensory cortex but is regulated according to each region within its processing network. The distinct A1, A2, V1, and V2 profiles observed here therefore support a circuit-dependent model in which PLD1 attenuation engages the form of structural remodeling most compatible with the computational requirements of each cortical field.

Within the auditory system, A1 maintains an organized representation of acoustic features but also participates directly in learning and memory[91]. Associative learning can modify A1 receptive fields and expand the cortical representation of behaviorally relevant sounds, while homeostatic synaptic scaling preserves auditory responsiveness following altered sensory input[92,93]. A1 therefore operates at the intersection of representational stability and targeted adaptive plasticity[94]. The focused VU0155069 response observed in A1 is consistent with this organization: rather than producing broad morphological reconfiguration, PLD1 attenuation preferentially recruited selected structural features that could support continued plasticity while preserving the established acoustic representation.

A2 occupies a different position within auditory processing, contributing to the integration and modulation of information that has already undergone initial cortical encoding. Experimental studies demonstrate that A1 and A2 make distinguishable contributions to auditory associative plasticity, with A1 supplying frequency-specific information and A2 modifying downstream processing through distinct synaptic pathways[95]. The broader and sustained A2 remodeling observed after VU0155069 treatment therefore dovetails with its more integrative role. Changes spanning spine phenotype, individual-spine geometry, and dendritic-segment area suggest that A2 can recruit several structural dimensions simultaneously to support the processing of behaviorally meaningful auditory information. Thus, the contrast between A1 and A2 is not one of responsiveness versus nonresponsiveness, rather it reflects complementary forms of plasticity, with A1 preserving a stable sensory representation through focused remodeling and A2 engaging a broader structural repertoire appropriate to higher-order integration.

A related distinction emerged within the visual cortex, although the structural strategy differed from that of the auditory fields. V1 is the initial cortical recipient and distributor of visual information and maintains a highly organized retinotopic representation[60]. Stable visuocortical mapping is essential because excessive reorganization could disrupt established computations for spatial correspondence, motion, depth, and object processing[96]. At the same time, V1 retains substantial experience-dependent and homeostatic plasticity. The persistent increase in V1 spine abundance following VU0155069 treatment may therefore represent a structural solution that expands the available synaptic substrate while preserving the basic organization of the primary visual representation. The accompanying reduction in average spine dimensions during later pathology further suggests that increased synaptic capacity can be achieved without generalized enlargement of the existing spine population.

V2 and other downstream visual areas integrate increasingly complex combinations of visual features and need not reproduce the plasticity program of V1[97]. Studies comparing early and higher visual cortices show that experience-dependent responses vary across the visual hierarchy and that plasticity in downstream fields is not simply inherited from V1. The V2 profile observed here is consistent with this principle. Increased spine abundance accompanied by coordinated redistribution between filamentous and thin profiles suggests that VU0155069 reorganizes the available spine population as visual information advances beyond the initial cortical stage. The difference between V1 and V2 therefore reflects complementary visual-circuit strategies: V1 predominantly expands the number of available synaptic sites while maintaining its organized cortical representation, whereas V2 combines expansion with subtype reorganization appropriate to downstream visual integration.

Viewed together, the four sensory regions reveal that PLD1 attenuation does not impose a common morphological endpoint on cortical circuits. Instead, VU0155069 appears to preserve the balance between stability and plasticity required by each field. A1 engages focused remodeling compatible with maintaining an established acoustic representation, A2 recruits broader multidimensional plasticity appropriate to auditory integration, V1 expands the synaptic substrate within a tightly organized retinotopic system, and V2 combines abundance with spine-phenotype redistribution during downstream visual processing. These anatomically differentiated responses provide a structural counterpart to the previously demonstrated functional efficacy of VU0155069 and support a model in which PLD1 attenuation promotes sensory-cortical resilience by allowing each circuit to engage its characteristic mode of synaptic adaptation.

### Associative memory regions engage complementary structural programs

The hippocampus and amygdala contribute complementary operations to associative memory, with the hippocampus organizing contextual and relational information and the amygdala assigning emotional and motivational significance to sensory experiences. Hippocampal and amygdalar interactions are therefore essential for linking what occurred, where it occurred, and whether the experience carried aversive or appetitive significance.

These memory systems also differ fundamentally in anatomical organization. The hippocampus is arranged as a highly ordered, laminar circuit in which entorhinal inputs are distributed across the dentate gyrus and CA subfields through anatomically segregated pathways, including the perforant path, mossy fibers, Schaffer collaterals, and direct entorhinal projections[98]. The amygdala is organized as a collection of interconnected nuclei in which sensory information enters predominantly through LA and the basolateral complex, is integrated with cortical, hippocampal, and neuromodulatory signals, and is transmitted through CeA and related pathways to coordinate learned behavioral and autonomic responses[99]. These contrasting anatomical arrangements provide a biologically grounded explanation for why VU0155069 produced complementary rather than uniform structural responses across associative-memory regions.

### Hippocampal subfields distribute associative-memory processing across specialized laminar pathways

The classical hippocampal circuit routes entorhinal information through DG granule cells to CA3 pyramidal neurons and then through Schaffer collaterals to CA1, while direct entorhinal projections and additional recurrent connections create parallel routes for memory encoding and retrieval[100]. Within this organization, DG and proximal CA3 contribute strongly to pattern separation, the recurrent CA3 network supports pattern completion, CA2 integrates unusually strong cortical input and coordinates hippocampal network dynamics, and CA1 serves as a major integrative output stage[64,101].

The previously published CA1 findings provide the functional and structural anchor for interpreting the present hippocampal profile. In the earlier 3xTg-AD cohort, repeated VU0155069 treatment preserved novel-object recognition and fear-conditioning performance, improved hippocampal synaptic plasticity, and supported CA1 dendritic spine integrity[13]. In the later cohort, the same therapeutic strategy preserved perirhinal cortex-, hippocampal-, and amygdala-dependent behaviors, improved HFS-LTP and LFS-LTD, and maintained selected mushroom-and filamentous-spine characteristics in CA1[14]. Together, these two studies established that attenuation of aberrant PLD1 signaling supports hippocampal function and CA1 structural resilience during both earlier and later pathology.

The present analysis extends that established CA1 profile upstream through CA2, CA3, and DG, revealing that the hippocampal circuit does not rely on one common form of structural remodeling. This interpretation is supported by the known specialization of hippocampal dendrites, because the number and location of synaptic inputs, dendritic morphology, and compartment-specific integrative properties differ substantially across hippocampal subfields.

CA2 displayed the broadest multidimensional response, consistent with its position as a computational hub receiving strong direct cortical input and participating in the coordination of hippocampal network activity[101]. CA2 pyramidal neurons receive substantially greater distal entorhinal excitation than CA1 pyramidal neurons, and their distinctive dendritic morphology and integrative properties enable efficient propagation of this cortical drive to the soma[64]. The combination of changes in spine abundance, phenotype distribution, and individual-spine geometry observed after VU0155069 treatment, therefore, suggests that CA2 can recruit several structural dimensions to preserve its role in integrating cortical information and coordinating hippocampal output.

The later CA2 combination of increased total spine number with lower local spine density also raises a biologically compelling arbor-level model. Experimental studies in hippocampal neurons have shown that PLD1 can restrict dendritic branching and that PLD1 knockdown or pharmacological inhibition increases dendritic complexity through a RhoA–PLD1–phosphatidic-acid pathway[62]. PLD1 attenuation may therefore release a constraint on CA2 dendritic growth, distributing a larger spine population across an expanded dendritic arbor and increasing the structural substrate available for strong cortical integration. Whole-neuron reconstruction and Sholl analysis can now test this focused model and complement the multidimensional spine pattern revealed by the present study.

CA3 engaged a more focused program that transitioned from increased spine number during earlier pathology to enlargement of individual-spine profiles at the later stage[66]. CA3 occupies a central position in hippocampal associative processing because DG mossy-fiber input converges with entorhinal and recurrent collateral input onto a network capable of retrieving complete representations from partial cues. The early expansion of spine abundance may increase the structural substrate available for convergent and recurrent input, whereas the later increase in spine major and minor axes may support remodeling of established synaptic contacts as pathology advances. This progression provides a plausible structural correlate of the improved CA3–CA1 Schaffer collateral plasticity reported after VU0155069 treatment in both the earlier and later 3xTg-AD cohorts[13,14].

DG exhibited a third hippocampal strategy centered on sustained expansion of spine abundance[74]. The DG is the principal cortical input gate to the hippocampus, and its sparse granule-cell activity contributes to separation of overlapping input patterns before transmission to CA3 through mossy fibers. The DG is also distinguished by the continuing addition of adult-born granule neurons, which acquire synaptic inputs and outputs as they mature and become incorporated into the existing hippocampal circuit. Adult-born granule cells provide a source of heightened structural and physiological plasticity and have been linked to pattern separation, contextual discrimination, memory encoding, and cognitive flexibility.

The progressive increase in DG spine abundance after VU0155069 treatment is therefore compatible with strengthened connectivity among mature granule neurons, enhanced maintenance of newly established synapses, or greater integration of adult-born granule-cell populations. PLD1 signaling has independently been implicated in neural-stem-cell differentiation and neurogenesis through Rho-family GTPase-, calcium-, and SHP-1/STAT3-associated pathways, providing a mechanistic basis for examining whether neuronal-lineage effects contribute to the DG response[62]. The present DG findings do not require neurogenesis to explain the abundance-centered profile, but they identify adult-born granule-cell integration as a biologically grounded mechanism that can be tested using birth dating and lineage-resolved imaging.

Viewed across the hippocampal circuit, the structural findings suggest a coordinated division of labor: DG expands the spine substrate available for cortical input, CA3 transitions from increased synaptic opportunity to remodeling of existing spine geometry, CA2 engages broad multidimensional plasticity, and CA1 preserves the output-stage spine integrity previously associated with improved hippocampal LTP and memory. The resulting sequence places the current CA2, CA3, and DG findings within the complete hippocampal profile established across the present and earlier VU0155069 studies.

### Amygdalar nuclei distribute associative processing across input, integration, and output nodes

Whereas the hippocampus partitions information through laminar pathways and ordered subfield connections, the amygdala distributes associative-memory processing across interconnected nuclei with distinct input-output relationships. LA is a major sensory-input and associative-plasticity site, the basolateral complex integrates sensory information with cortical, hippocampal, and neuromodulatory signals, and CeA transforms this processed information into coordinated behavioral and autonomic responses[81,82]. CeA also contributes directly to acquisition and consolidation of conditioned fear, indicating that CeA is not simply a passive output relay[102].

The distinct LA, BLA, and CeA spine profiles observed after VU0155069 treatment dovetail with this nucleus-based organization. LA progressed from early expansion and redistribution of the available spine population to later enlargement of individual-spine profiles, a pattern compatible with its role in forming and maintaining sensory associations. Learning-related studies have shown that fear and safety conditioning bidirectionally alter synapse size and local dendritic translation in LA, establishing that LA synaptic structure is closely coupled to the encoding of associative significance[86].

BLA followed a more phenotype-centered trajectory, with an early filamentous-spine-enriched profile followed by greater mushroom-spine representation during later pathology[103]. The basolateral complex is extensively interconnected with higher-order cortical and hippocampal regions and contains diverse excitatory principal neurons and inhibitory microcircuits that regulate the encoding and expression of emotionally salient experiences. Experience-dependent changes in BLA spine density and dendritic organization accompany fear acquisition and extinction, supporting the biological relevance of a staged structural response within this integrative nucleus. CeA progressed from early redistribution among operational spine phenotypes to later enlargement across several individual-spine and dendritic-segment measures[79]. CeA integrates sensory and associative information and coordinates adaptive responses through molecularly and functionally diverse neuronal populations with extensive subcortical outputs. The later CeA morphometric profile may therefore reflect consolidation of the structural substrate through which processed associative information is translated into the behavioral and autonomic components of learned responses.

This nucleus-specific organization aligns with our group publications of PLD as a convergent signaling mechanism in amygdala-dependent associative plasticity[9–11]. The current LA, BLA, and CeA findings add a structural dimension to that established signaling framework. PLD1 attenuation may preserve early structural flexibility in sensory-associative and integrative amygdalar nuclei and support later consolidation of the synaptic substrate through nucleus-appropriate remodeling. The differing LA, BLA, and CeA profiles therefore do not represent inconsistent treatment effects; they reflect complementary structural solutions aligned with the input, integration, and output functions of the amygdalar network.

### A distributed structural framework for VU0155069-associated associative-memory resilience

The hippocampal and amygdalar findings together support a distributed model of VU0155069-associated associative-memory resilience. The hippocampus engaged subfield-specific structural programs within a laminar network specialized for contextual representation, pattern separation, pattern completion, and integrated memory output. The amygdala engaged nucleus-specific programs across sensory input, associative integration, and behavioral-output pathways involved in assigning emotional significance to those representations.

This systems-level interpretation is strengthened by the prior VU0155069 studies[13,14]. Rather than requiring one morphology to explain every functional benefit, the combined findings suggest that PLD1 attenuation preserves network performance through complementary structural adaptations. DG may increase the synaptic substrate available for cortical input, CA3 may adjust spine abundance and geometry to support recurrent processing, CA2 may engage broad dendritic and spine remodeling to integrate cortical information, and CA1 may preserve the final hippocampal representation transmitted to downstream targets. Within the amygdala, LA may preserve sensory-associative adaptability, BLA may reorganize spine phenotypes within an integrative network, and CeA may consolidate the structures that translate associative information into coordinated responses.

Thus, the hippocampal and amygdalar findings extend the VU0155069 therapeutic profile from isolated regional outcomes to an integrated associative-memory network. The multiregional approach reveals that PLD1 attenuation supports synaptic resilience not through a single morphological endpoint, but by engaging the structural requirements of each laminar hippocampal subfield and each functionally specialized amygdalar nucleus. These complementary structural programs provide a systems-level anatomical correlate of the previously demonstrated preservation of recognition memory, fear conditioning, LTP, and LTD and establish a focused foundation for regionally matched physiological and molecular studies.

### A PLD1-dependent lipid-signaling framework for regional spine remodeling

The multiregional findings can be integrated within a PLD1-dependent lipid-signaling framework. PLD1 hydrolyzes phosphatidylcholine to generate phosphatidic acid, a bioactive lipid that regulates membrane curvature, vesicle trafficking, protein recruitment, and cytoskeletal organization[104–106]. Each of these processes is directly relevant to dendritic spine formation and remodeling. Excessive post developmental PLD1 signaling in AD may disturb the balance between membrane supply, receptor trafficking, and actin-dependent spine structure[12,21,107]. VU0155069 may restore this balance by reducing aberrant PLD1 activity and allowing regionally appropriate structural plasticity to re-emerge[108,109].

The prior behavioral, electrophysiological, biochemical, and selected CA1 studies support this model[13,14]. VU0155069 improved synaptic plasticity and memory at both early and later stages, while the present work demonstrates that the same treatment paradigm reorganizes spine architecture across distributed circuits. The convergence of functional and structural effects is consistent with PLD1 acting upstream of processes that govern both synaptic transmission and morphology. The region-specific response patterns may reflect differences in basal PLD1 tone, local phosphatidic-acid metabolism, disease burden, circuit activity, and the cellular sources of PLD1 signaling.

Astrocytic PLD1 and phosphatidic acid provide an additional mechanistic dimension. Astrocyte-derived phosphatidic acid can influence neuronal dendritic branching in culture, supporting the concept that PLD1-linked lipid signaling can act across cell types[20]. In the AD brain, astrocytes may shape synaptic structure through lipid release, inflammatory signaling, extracellular-matrix regulation, and metabolic support. Neuronal and astrocytic PLD1 may,therefore, contribute in complementary ways to the regional structural phenotypes observed here. This possibility is especially relevant to regions such as CA2, CeA, and LA, where broad remodeling involves both spine and dendritic-segment measures.

PLD1-generated phosphatidic acid also participates in vesicle docking and fusion, providing a direct link between lipid signaling and synaptic membrane trafficking[110]. Reduced aberrant PLD1 activity could alter the availability or localization of phosphatidic acid at synaptic membranes, thereby influencing receptor trafficking, membrane expansion, and actin-associated spine geometry. The present results do not resolve the relative contributions of these processes, but they identify the regional structural outcomes that a mechanistic study must explain.

One reported PLD1-independent action of VU0155069 is inhibition of inflammasome-associated signaling in macrophage and sepsis models[111]. This observation does not weaken the PLD1 framework. Rather, it identifies an additional pharmacological property that may contribute to the net in vivo response and should be tested directly in brain tissue. Genetic attenuation of PLD1, catalytic activity measurements, regional phosphatidic-acid profiling, and inflammatory readouts will allow the PLD1-dependent and PLD1-independent components of VU0155069 action to be separated. The strongest mechanistic model will be the one that explains both pharmacological and genetic effects across the most responsive regions.

### Structural remodeling supports the therapeutic concept of synaptic resilience

The therapeutic significance of the present findings lies in the preservation of structural plasticity across both early and later disease stages. Current AD treatments are often evaluated primarily by their effects on pathological protein burden. The PLD1 strategy addresses a complementary objective: maintaining the structural and functional capacity of synapses within circuits already exposed to Aβ-and tau-associated stress. The earlier studies demonstrated behavioral and physiological resilience, and the current data show that this resilience is accompanied by widespread but anatomically organized structural remodeling.

Regional selectivity should be viewed as a therapeutic feature rather than a shortcoming. Neural circuits differ in baseline architecture and computational role, so an intervention that permits each region to engage an appropriate structural program may be preferable to one that forces a uniform morphological response. V1 and DG primarily expanded spine abundance, A2 and CA2 engaged multidimensional remodeling, and the amygdala shifted from early phenotype redistribution to later structural consolidation. These distinct programs converge on the broader outcome of maintaining a responsive synaptic substrate across distributed memory networks.

The persistence of structural responsiveness at 12 months is especially encouraging. By this stage, 3xTg-AD mice display a more advanced combination of amyloid-and tau-associated abnormalities, yet VU0155069 continued to modify spine architecture in sensory, hippocampal, and amygdalar regions. The findings therefore support continued investigation of PLD1 attenuation not only as an early preventive strategy but also as an intervention capable of engaging residual synaptic plasticity after pathology is established.

This therapeutic model is also compatible with the distributed nature of cognition. Recognition memory, contextual learning, associative learning, and sensory integration emerge from interactions among cortical, hippocampal, and amygdalar circuits. The present study shows that PLD1 attenuation reaches structural components across these networks. The combination of regional specificity and network breadth provides a strong rationale for evaluating PLD1-linked interventions using multimodal endpoints that integrate molecular signaling, spine architecture, circuit physiology, and behavior.

### Strengths and limitations

The major strength of this study is its systems-level anatomical design. Ten regions were analyzed using the same overall treatment paradigm, acquisition platform, segment-selection framework, and morphometric definitions. The study included paired primary and secondary sensory cortices, three hippocampal subfields, and three amygdalar nuclei, allowing the structural response to be compared across functionally related circuits. The extensive parameter set captured spine abundance, phenotype distribution, and geometry rather than relying on density alone. New regional imaging was generated from archived specimens, and none of the ten-region measurements or figures duplicated the previously published CA1 data.

Several additional features support the internal consistency of the work. Littermates were assigned to vehicle or VU0155069 treatment, treatment groups were represented across cages and dosing batches, and vehicle-and VU0155069-treated samples were processed together within each age cohort. Regional sampling used three sections and five nonoverlapping fields per section, with one standardized 10-µm segment selected per field. Analysts used common classification criteria, coded specimens, spatial calibration, and cross-review during training. All 20 animals completed treatment and contributed usable measurements from all ten regions.

The main limitations concern biological sample size and analytical hierarchy. Each group contained five animals, while multiple segments and spines were sampled within each animal. These nested observations provide valuable measurement depth but do not increase the number of independent biological replicates. The established analyses used pooled measurement-level distributions, and no global correction was applied across every region and endpoint. The most persuasive findings are, therefore, the recurring, directionally coherent patterns supported across related measures and, where applicable, by Dunn-adjusted comparisons. Independent replication with larger animal-level samples and hierarchical models will be important for confirmation.

The 6-and 12-month groups were separate cohorts processed in different age-specific Golgi-Cox batches, and imaging was completed over multiple periods. Accordingly, the study is strongest for vehicle-versus-VU0155069 comparisons within each age cohort. The age-associated patterns remain biologically informative, but should not be treated as longitudinal observations from the same animals. In addition, sampled dendrites were localized anatomically, but were not traced to parent somata or assigned to a neuronal class, cortical layer, or apical versus basal compartment. Future cell-type-and compartment-resolved studies will refine the regional patterns reported here.

Golgi-Cox staining and full-focus reconstruction also impose method-specific boundaries. Golgi methods may detect fewer spines and yield larger estimates of selected dimensions than intracellular fluorescent fills[112,113]. Morphological categories simplify a continuum of spine forms, and zero-coded mushroom dimensions combine subtype absence with dimensions among present spines. These considerations affect the absolute interpretation of individual measurements but do not negate the matched within-method comparisons that define the regional treatment responses.

Finally, only female 3xTg-AD mice were included, and no non-transgenic reference groups were analyzed. The female-focused design is relevant to the higher burden of AD in women and to reported female vulnerability in the 3xTg-AD model, but both sex replication is needed to assess generalizability[1,49]. Age-matched nontransgenic groups would provide an additional physiological reference for determining whether individual morphometric endpoints approach the range observed in the healthy brain. Such comparisons would address normalization, whereas the present age-matched disease-vehicle comparison addresses the direct pharmacological question of whether PLD1 attenuation modifies synaptic architecture within the affected population. Regional PLD1 activity, phosphatidic-acid species, pathology, and target engagement were not measured concurrently in these specimens. These are focused mechanistic priorities for the next phase rather than reasons to diminish the present systems-level structural advance.

## Conclusions

Repeated VU0155069 treatment reorganized dendritic spine architecture across distributed cortical, hippocampal, and amygdalar circuits in female 3xTg-AD mice. The response was regionally and stage differentiated, with three recurring structural programs: expansion of spine abundance, redistribution among spine phenotypes, and morphometric reorganization of individual spines and dendritic segments. Primary and secondary cortical fields, hippocampal subregions, and amygdalar nuclei engaged these programs in distinct combinations, demonstrating that PLD1 attenuation promotes synaptic structural resilience through circuit-appropriate rather than uniform remodeling.

These findings extend the PLD1 story from human AD synaptic fractions and selected CA1 outcomes to a systems-level map of structural responsiveness across ten additional regions. In combination with our prior behavioral, electrophysiological, biochemical, and neuropathological studies, the present data support a coherent model in which attenuation of aberrant PLD1 signaling preserves the capacity of distributed neural networks to remodel under Aβ-and tau-associated stress.

The persistence of remodeling at both 6 and 12 months broadens the therapeutic relevance of PLD1 attenuation. The data support continued development of PLD1-linked interventions as a strategy to preserve synaptic resilience during early pathology and to engage residual structural plasticity after disease progression. Regionally matched PLD1 activity, phosphatidic-acid profiling, pathology, physiology, and cell-type-specific genetic studies will now be essential for defining the mechanisms that generate each structural response mode.

Overall, the study establishes regional identity and pathological stage as central determinants of the structural response to VU0155069. This systems-level perspective provides a stronger foundation for understanding PLD1 biology in AD and identifies the cortical, hippocampal, and amygdalar circuits most likely to yield mechanistic and translational insight in the next generation of studies.

## Acknowledgments

The authors acknowledge Batbayar Tumurbaatar for maintaining the 3xTg-AD breeding colony, monitoring breeding-cohort transgene integrity, and assisting with animal provision. The authors thank Giulio Taglialatela and the Mitchell Center for Neurodegenerative Diseases for supporting the breeding program and shared research resources and acknowledge the UTMB Department of Neurology for institutional support. The authors thank all current and past Krishnan Lab members for the editing and revision process as well as their support.

## Author Contributions

Conceptualization: Balaji Krishnan. Methodology: Balaji Krishnan, Chandramouli Natarajan, and Karthik Ramaswamy. Investigation and formal analysis: Balaji Krishnan, Karthik Ramaswamy, Sarah Bosworth, and Sanjana Mohanty. Data curation: Sanjana Mohanty and Balaji Krishnan. Visualization: Balaji Krishnan and Sanjana Mohanty. Resources, supervision, project administration, and funding acquisition: Balaji Krishnan. Writing -original draft: Balaji Krishnan and Sanjana Mohanty. Writing-review and editing: Balaji Krishnan, Sanjana Mohanty, Sarah Bosworth, Karthik Ramaswamy, and Chandramouli Natarajan. Sanjana Mohanty and Sarah Bosworth contributed equally to this work.

## Statements and Declarations

### Ethical Considerations

All animal procedures were approved by the University of Texas Medical Branch Institutional Animal Care and Use Committee under protocol 1710061, active from October 2018 through both experimental cohorts. Procedures were performed by trained personnel in accordance with the NIH

Guide for the Care and Use of Laboratory Animals. Predefined humane endpoints were incorporated into the approved protocol; no animal reached a humane endpoint or required premature euthanasia.

### Consent to participate

Not applicable.

### Consent for publication

Not applicable.

### Declaration of conflicting interest

The authors declared no potential conflicts of interest with respect to the research, authorship, and/or publication of this article.

### Funding

This work was supported by Alzheimer’s Association Research Grant AARG-17-533363 and by the National Institute on Aging under award numbers R21 AG059223 and R01 AG063945 to Balaji Krishnan. Sanjana Mohanty and Chandramouli Natarajan received training support from NIA T32 AG067952, Predoctoral and Postdoctoral Training in Alzheimer’s Pathophysiology. Additional support was provided by the Don and Nancy Mafrige Professorship in Neurodegenerative Diseases, the Mitchell Center for Neurodegenerative Diseases, and the UTMB Department of Neurology. The content is solely the responsibility of the authors and does not necessarily represent the official views of the funding organizations.

### Data Availability

Animal-level summary data and complete statistical outputs are provided in the Supplementary Material. Underlying Keyence image stacks, full-focus TIFF images, ImageJ measurement files, GraphPad Prism files, and associated source data are available from the corresponding author upon reasonable request and, where appropriate, through controlled access to the institutional data repository.

### Conflict of Interest

There are no conflicts of interest to report.

---

**Supplementary Table S1A.** Descriptive statistics and within-age Mann–Whitney results for A1. Values represent the mean ± standard error of mean (SEM) for each dendritic spine parameter analyzed using the Mann-Whitney test, with the total number of pooled measurement-level observation values indicated in the appropriate labeled column with the total number of animals denoted by n (n =5). Spine parameters were compared between ages (6-months and 12-months) and across treatment groups (vehicle and VU01) which are separated by columns. Endpoints refer to the specific dendritic spine parameters evaluated in the study including the observation level (Panels A-M). Median values were reported, including Mann-Whitney U-values which indicate the degree of rank separation between groups and the p-value to describe the statistically significant differences relative to α = 0.05.

| Panel | Endpoint | Observation level | 6-mo Veh n | 6-mo Veh mean ± SEM | 6-mo Veh median | 6-mo VU01 n | 6-mo VU01 mean ± SEM | 6-mo VU01 median | MW U, 6 mo | Exact p, 6 mo | 12-mo Veh n | 12-mo Veh mean ± SEM | 12-mo Veh median | 12-mo VU01 n | 12-mo VU01 mean ± SEM | 12-mo VU01 median | MW U; p, 12 mo |
| --- | --- | --- | --- | --- | --- | --- | --- | --- | --- | --- | --- | --- | --- | --- | --- | --- | --- |
| A | Number of all Spines | Dendritic segment | 60 | 4.983 ± 0.3085 | 4 | 75 | 4.987 ± 0.2515 | 5 | 2213 | 0.8681 | 75 | 6.267 ± 0.3235 | 6 | 75 | 6.053 ± 0.2815 | 6 | 2710; 0.6980 |
| B | Stubby Spines | Dendritic segment | 28 | 0 ± 0 | 0 | 34 | 0 ± 0 | 0 | 476 | >0.9999 | 38 | 0.02632 ± 0.2632 | 0 | 35 | 0 ± 0 | 0 | 647.5; >0.9999 |
| C | Filamentous Spines | Dendritic segment | 28 | 3.179 ± 0.3639 | 3 | 31 | 3.065 ± 0.2661 | 3 | 422 | 0.8577 | 59 | 1.458 ± 0.2373 | 1 | 59 | 1.288 ± 0.1889 | 1 | 1704; 0.8394 |
| D | Thin Spines | Dendritic segment | 28 | 0.5357 ± 0.1957 | 0 | 31 | 0.6129 ± 0.1515 | 0 | 385.5 | 0.3984 | 38 | 0.5263 ± 0.1398 | 0 | 36 | 1.278 ± 0.2093 | 1 | 419; 0.0018 |
| E | Mushroom Spines | Mushroom spine | 28 | 2.214 ± 0.2880 | 1.5 | 31 | 2.129 ± 0.2772 | 2 | 427 | 0.9132 | 38 | 2.711 ± 0.3108 | 2.5 | 36 | 2.667 ± 0.2760 | 2 | 670; 0.8788 |
| F | Spine Density per 10 μm | Dendritic segment | 60 | 13.55 ± 0.2924 | 12.66 | 75 | 13.38 ± 0.2199 | 13.05 | 2235 | 0.9481 | 75 | 14.42 ± 0.2954 | 13.99 | 75 | 14.24 ± 0.2560 | 13.72 | 2785; 0.9187 |
| G | Individual Spine Area | Individual spine | 303 | 0.1873 ± 0.01158 | 0.1030 | 792 | 0.1688 ± 0.005720 | 0.1140 | 118730 | 0.7881 | 469 | 0.1991 ± 0.01081 | 0.1030 | 455 | 0.2001 ± 0.01694 | 0.1140 | 103540; 0.4360 |
| H | Total Dendritic Area | Dendritic segment | 60 | 5.697 ± 0.2267 | 5.166 | 75 | 5.922 ± 0.1967 | 5.505 | 1941 | 0.1721 | 75 | 6.189 ± 0.2748 | 5.784 | 75 | 5.655 ± 0.2003 | 5.123 | 2519; 0.2712 |
| I | Spine Perimeter | Individual spine | 299 | 1.646 ± 0.05624 | 1.377 | 801 | 1.519 ± 0.03845 | 1.270 | 112199 | 0.1071 | 470 | 1.676 ± 0.0545 | 1.270 | 454 | 1.632 ± 0.04534 | 1.421 | 103422; 0.4200 |
| J | Dendrite Major Axis | Dendritic segment | 299 | 0.7065 ± 0.02271 | 0.6040 | 405 | 0.7952 ± 0.03404 | 0.6040 | 59019 | 0.5663 | 470 | 0.7066 ± 0.02131 | 0.5595 | 490 | 0.8393 ± 0.03519 | 0.6090 | 105061; 0.0187 |
| K | Dendrite Minor Axis | Dendritic segment | 299 | 0.2391 ± 0.01210 | 0.1860 | 374 | 0.2335 ± 0.01024 | 0.1750 | 55264 | 0.7905 | 470 | 0.2592 ± 0.01118 | 0.1665 | 454 | 0.2456 ± 0.009786 | 0.1910 | 106346; 0.9308 |
| L | Mushroom spine Head Width | Mushroom spine | 135 | 0.5763 ± 0.02999 | 0.4770 | 198 | 0.6417 ± 0.02576 | 0.5595 | 11894 | 0.0881 | 208 | 0.5820 ± 0.02635 | 0.4830 | 192 | 0.5382 ± 0.02407 | 0.4590 | 18861; 0.3380 |
| M | Mushroom Spine Neck Length | Mushroom spine | 124 | 0.3033 ± 0.01832 | 0.2765 | 124 | 0.3121 ± 0.01344 | 0.2780 | 11720 | 0.4939 | 208 | 0.2928 ± 0.01532 | 0.2600 | 192 | 0.2875 ± 0.01437 | 0.2460 | 19929; 0.9731 |

**Supplementary Table S1B.** Kruskal–Wallis and Dunn-adjusted comparisons for A1. Statistical differences obtained after conducting the Kruskal-Wallis test for all endpoint dendritic spine parameters depicted in each panel (Panels A-M). Dunn’s post-hoc test was applied for comparison across all combinations of treatment (vehicle and VU01) and ages (6-months and 12-months) to isolate specifically significant group pairs. Omnibus p-values reflect overall significance while specific pairwise differences between treatment and age groups were determined and adjusted via Dunn’s post-hoc test, with an α = 0.05. Mean ranks were calculated for each comparison to indicate the relative distribution of data within groups.

| Panel | Endpoint | KW omnibus p | 6 Veh vs 6 VU01 rank diff. | adj. p | 6 Veh vs 12 Veh rank diff. | adj. p | 6 Veh vs 12 VU01 rank diff. | adj. p | 6 VU01 vs 12 Veh rank diff. | adj. p | 6 VU01 vs 12 VU01 rank diff. | adj. p | 12 Veh vs 12 VU01 rank diff. | adj. p |
| --- | --- | --- | --- | --- | --- | --- | --- | --- | --- | --- | --- | --- | --- | --- |
| A | Number of all Spines | 0.0017 | -1.998 | >0.9999 | -40.00 | 0.0286 | -37.77 | 0.0461 | -38.00 | 0.0267 | -35.77 | 0.0445 | 2.227 | >0.9999 |
| B | Stubby Spines | 0.4659 | 0 | >0.9999 | -1.776 | >0.9999 | 0 | >0.9999 | -1.776 | >0.9999 | 0 | >0.9999 | 1.776 | >0.9999 |
| C | Filamentous Spines | <0.0001 | -2.300 | >0.9999 | 48.60 | 0.0001 | 50.81 | <0.0001 | 50.90 | <0.0001 | 53.11 | <0.0001 | 2.212 | >0.9999 |
| D | Thin Spines | 0.0027 | -6.889 | >0.9999 | -2.331 | >0.9999 | -28.02 | 0.0082 | 4.559 | >0.9999 | -21.13 | 0.0779 | -25.69 | 0.0088 |
| E | Mushroom Spines | 0.3323 | 1.982 | >0.9999 | -10.07 | >0.9999 | -11.46 | >0.9999 | -12.05 | >0.9999 | -13.44 | 0.8246 | -1.392 | >0.9999 |
| F | Spine Density per 10 μm | 0.0079 | 2.670 | >0.9999 | -32.26 | 0.1429 | -32.03 | 0.1491 | -34.93 | 0.0567 | -34.70 | 0.0596 | 0.2333 | >0.9999 |
| G | Individual Spine Area | 0.6694 | -7.267 | >0.9999 | 34.00 | >0.9999 | 1.009 | >0.9999 | 41.27 | >0.9999 | 8.276 | >0.9999 | -33.00 | >0.9999 |
| H | Total Dendritic Area | 0.4164 | -16.98 | >0.9999 | -14.77 | >0.9999 | 0.9383 | >0.9999 | 2.213 | >0.9999 | 17.92 | >0.9999 | 15.71 | >0.9999 |
| I | Spine Perimeter | 0.2922 | 63.43 | 0.6545 | 49.51 | >0.9999 | 13.79 | >0.9999 | -13.92 | >0.9999 | -49.64 | 0.8880 | -35.72 | >0.9999 |
| J | Dendrite Major Axis | 0.0932 | -21.54 | >0.9999 | 40.17 | >0.9999 | -33.70 | >0.9999 | 61.71 | 0.3482 | -12.16 | >0.9999 | -73.87 | 0.1031 |
| K | Dendrite Minor axis | 0.8768 | 8.628 | >0.9999 | -11.60 | >0.9999 | -14.59 | >0.9999 | -20.23 | >0.9999 | -23.22 | >0.9999 | -2.990 | >0.9999 |
| L | Spine Head Width | 0.0231 | -39.14 | 0.5858 | 21.51 | >0.9999 | 25.56 | >0.9999 | 43.19 | 0.2393 | 64.70 | 0.0153 | 21.51 | >0.9999 |
| M | Spine Neck Length | 0.2296 | -17.30 | >0.9999 | 18.61 | >0.9999 | 20.88 | >0.9999 | 35.91 | 0.4970 | 38.18 | 0.4240 | 2.268 | >0.9999 |

**Supplementary note.** n denotes animals; number of observations denotes pooled measurement-level observations. The endpoint-specific unit is a dendritic segment, individual spine, or mushroom spine. Multiple observations were nested within n = 5 animals per group. Zero subtype counts indicate that no spine of the corresponding subtype was detected. For mushroom-spine dimensions, zero indicates absence of a mushroom spine rather than a physically measured dimension of zero. Dunn adjustment was applied to six comparisons within each endpoint-specific four-group family; no global correction was applied across regions and endpoints.

**Supplementary Table S2A.** Descriptive statistics and within-age Mann–Whitney results for A2. Mann-Whitney statistics, median and mean with SEM values for the A2 region are reported in this table. Statistical metrics, abbreviations and parameters presented here use the same reporting conventions as described in Supplementary Table S1A which can be referred for a detailed explanation of these terms and parameters.

| Panel | Endpoint | Observation level | 6-mo Veh Observation<br>s | 6-mo Veh mean ± SEM | 6-mo Veh median | 6-mo VU01 n | 6-mo VU01 mean ± SEM | 6-mo VU01 median | MW U, 6 mo | Exact p, 6 mo | 12-mo Veh n | 12-mo Veh mean ± SEM | 12-mo Veh median | 12-mo VU01 n | 12-mo VU01 mean ± SEM | 12-mo VU01 median | MW U; p, 12 mo |
| --- | --- | --- | --- | --- | --- | --- | --- | --- | --- | --- | --- | --- | --- | --- | --- | --- | --- |
| Panel | Endpoint | Observation level | 6 mo Sal n | 6 mo Sal mean ± SEM | 6 mo Sal median | 6 mo Inh n | 6 mo Inh mean ± SEM | 6 mo Inh median | MW U (6 mo) | MW p (6 mo) | 12 mo Sal n | 12 mo Sal mean ± SEM | 12 mo Sal median | 12 mo Inh n | 12 mo Inh mean ± SEM | 12 mo Inh median | MW U; p (12 mo) |
| A | Number of all Spines | Dendritic segment | 60 | 6.367 ± 0.3386 | 6 | 75 | 6.773 ± 0.3076 | 7 | 2077 | 0.4413 | 75 | 5.893 ± 0.3163 | 6 | 75 | 5.853 ± 0.3047 | 5 | 2744; 0.7951 |
| B | Stubby Spines | Dendritic segment | 35 | 0 ± 0 | 0 | 57 | 0.0175 ± 0.0175 | 0 | 980 | >0.9999 | 42 | 0 ± 0 | 0 | 53 | 0 ± 0 | 0 | 1113; >0.9999 |
| C | Filamentous Spines | Dendritic segment | 35 | 4.686 ± 0.373 | 4 | 57 | 3.807 ± 0.2678 | 4 | 772 | 0.0658 | 42 | 4.429 ± 0.3376 | 4 | 53 | 3.302 ± 0.2592 | 3 | 767.5; 0.0082 |
| D | Thin Spines | Dendritic segment | 35 | 0.7429 ± 0.1849 | 0 | 57 | 0.8947 ± 0.1516 | 1 | 910.5 | 0.448 | 42 | 0.619 ± 0.1522 | 0 | 53 | 0.5849 ± 0.1092 | 0 | 1095; 0.8797 |
| E | Mushroom Spines | Mushroom spine | 35 | 1.8 ± 0.187 | 1 | 57 | 2.474 ± 0.2184 | 2 | 762 | 0.0452 | 42 | 2.19 ± 0.2189 | 1.5 | 53 | 2.396 ± 0.1892 | 2 | 996; 0.3615 |
| F | Spine Density per 10 µm | Dendritic segment | 60 | 14.63 ± 0.2812 | 14.34 | 75 | 15.04 ± 0.3023 | 14.7 | 2093 | 0.4896 | 75 | 13.73 ± 0.2384 | 13.49 | 75 | 14.4 ± 0.2795 | 13.76 | 2401; 0.1224 |
| G | Individual Spine Area | Individual spine | 380 | 0.1917 ± 0.0107 | 0.111 | 508 | 0.2332 ± 0.0117 | 0.125 | 89064 | 0.0486 | 442 | 0.1555 ± 0.0075 | 0.097 | 438 | 0.2156 ± 0.0105 | 0.131 | 80298; <0.0001 |
| H | Total Dendritic Area | Dendritic segment | 60 | 5.642 ± 0.2059 | 5.22 | 75 | 6.82 ± 0.2502 | 6.303 | 1495 | 0.0007 | 75 | 5.398 ± 0.1641 | 4.878 | 75 | 5.867 ± 0.2089 | 5.14 | 2421; 0.1412 |
| I | Spine Perimeter | Individual spine | 382 | 1.684 ± 0.0517 | 1.421 | 508 | 1.777 ± 0.0506 | 1.395 | 94762 | 0.5502 | 442 | 1.475 ± 0.0409 | 1.27 | 439 | 1.787 ± 0.0544 | 1.44 | 82015; <0.0001 |
| J | Dendrite Major Axis | Dendritic segment | 382 | 0.7469 ± 0.0322 | 0.604 | 508 | 0.738 ± 0.0199 | 0.604 | 96201 | 0.8274 | 441 | 0.6272 ± 0.0164 | 0.534 | 439 | 0.7492 ± 0.0213 | 0.609 | 82166; 0.0001 |
| K | Dendrite Minor Axis | Dendritic segment | 382 | 0.2403 ± 0.0112 | 0.151 | 508 | 0.2982 ± 0.011 | 0.224 | 81394 | <0.0001 | 441 | 0.2285 ± 0.0094 | 0.151 | 440 | 0.2821 ± 0.0116 | 0.202 | 84302; 0.0006 |
| L | Mushroom Spine Head Width | Mushroom spine | 126 | 0.6232 ± 0.02481 | 0.5310 | 286 | 0.6214 ± 0.01976 | 0.5390 | 17390 | 0.5730 | 185 | 0.4876 ± 0.02022 | 0.4070 | 252 | 0.5857 ± 0.02057 | 0.5060 | 18935; 0.0008 |
| M | Mushroom Spine Neck Length | Mushroom spine | 126 | 0.3428 ± 0.0195 | 0.3285 | 286 | 0.345 ± 0.0133 | 0.3185 | 17686 | 0.7659 | 185 | 0.2818 ± 0.0125 | 0.27 | 252 | 0.3341 ± 0.0129 | 0.304 | 19660; 0.0051 |

**Supplementary Table S2B.** Kruskal–Wallis and Dunn-adjusted comparisons for A2. Differences among groups were assessed using the Kruskal-Wallis test. Statistical measures, abbreviations and parameters are defined in greater detail in Supplementary Table S1B which can be referred to.

| Panel | Endpoint | KW omnibus p | 6 Veh vs 6 VU01 rank diff. | adj. p | 6 Veh vs 12 Veh rank diff. | adj. p | 6 Veh vs 12 VU01 rank diff. | adj. p | 6 VU01 vs 12 Veh rank diff. | adj. p | 6 VU01 vs 12 VU01 rank diff. | adj. p | 12 Veh vs 12 VU01 rank diff. | adj. p |
| --- | --- | --- | --- | --- | --- | --- | --- | --- | --- | --- | --- | --- | --- | --- |
| A | Number of all Spines | 0.0678 | -11.54 | >0.9999 | 16.02 | >0.9999 | 20.19 | 0.9267 | 27.56 | 0.2358 | 31.73 | 0.1059 | 4.173 | >0.9999 |
| B | Stubby Spines | 0.5162 | -1.64 | >0.9999 | 0 | >0.9999 | 0 | >0.9999 | 1.64 | >0.9999 | 1.64 | >0.9999 | 0 | >0.9999 |
| C | Filamentous Spines | 0.0089 | 20.61 | 0.435 | 5.376 | >0.9999 | 34.72 | 0.0171 | -15.23 | 0.9657 | 14.11 | 0.9992 | 29.34 | 0.0472 |
| D | Thin Spines | 0.5097 | -8.517 | >0.9999 | 4.521 | >0.9999 | 3.258 | >0.9999 | 13.04 | >0.9999 | 11.77 | >0.9999 | -1.263 | >0.9999 |
| E | Mushroom Spines | 0.1395 | -22.18 | 0.2709 | -13.46 | >0.9999 | -23.8 | 0.2041 | 8.721 | >0.9999 | -1.624 | >0.9999 | -10.35 | >0.9999 |
| F | Spine Density per 10 μm | 0.0095 | -8.265 | >0.9999 | 34.65 | 0.0913 | 14.18 | >0.9999 | 42.91 | 0.0086 | 22.45 | 0.5721 | -20.47 | 0.7700 |
| G | Individual Spine Area | <0.0001 | -67.35 | 0.3100 | 87.52 | 0.0853 | -62.94 | 0.4709 | 154.9 | <0.0001 | 4.403 | >0.9999 | -150.5 | <0.0001 |
| H | Total Dendritic Area | <0.0001 | -48.33 | 0.0043 | 11.47 | >0.9999 | -8.142 | >0.9999 | 59.8 | <0.0001 | 40.19 | 0.017 | -19.61 | 0.8702 |
| I | Spine Perimeter | 0.0001 | -19.77 | >0.9999 | 106 | 0.0179 | -31.26 | >0.9999 | 125.8 | 0.0009 | -11.48 | >0.9999 | -137.3 | 0.0004 |
| J | Dendrite Major Axis | 0.0003 | 8.331 | >0.9999 | 118.7 | 0.0053 | -15.27 | >0.9999 | 110.4 | 0.0054 | -23.6 | >0.9999 | -134 | 0.0006 |
| K | Dendrite Minor Axis | <0.0001 | -145.1 | 0.0001 | 5.397 | >0.9999 | -109.7 | 0.0106 | 150.5 | <0.0001 | 35.37 | >0.9999 | -115.1 | 0.0040 |
| L | Mushroom Spine Head Width | <0.0001 | 20.37 | >0.9999 | 130.2 | <0.0001 | 49.56 | 0.3834 | 109.8 | <0.0001 | 29.19 | >0.9999 | -80.65 | 0.0041 |
| M | Mushroom Spine Neck Length | 0.0062 | 7.837 | >0.9999 | 79.76 | 0.0292 | 14.59 | >0.9999 | 71.92 | 0.0113 | 6.750 | >0.9999 | -65.17 | 0.0363 |

**Supplementary note.** n denotes animals; number of observations denotes pooled measurement-level observations. The endpoint-specific unit is a dendritic segment, individual spine, or mushroom spine. Multiple observations were nested within n = 5 animals per group. Zero subtype counts indicate that no spine of the corresponding subtype was detected. For mushroom-spine dimensions, zero indicates absence of a mushroom spine rather than a physically measured dimension of zero. Dunn adjustment was applied to six comparisons within each endpoint-specific four-group family; no global correction was applied across regions and endpoints.

**Supplementary Table S3A.** Descriptive statistics and within-age Mann–Whitney results for V1. Mann-Whitney U-test statistics, medians and means ± SEM for the V1 region are shown in this table. Statistical measures, abbreviations and parameters follow similar conventions as shown in Supplementary Table S1A.

| Panel | Endpoint | Observation level | 6-mo Veh n | 6-mo Veh mean ± SEM | 6-mo Veh median | 6-mo VU01 n | 6-mo VU01 mean ± SEM | 6-mo VU01 median | MW U, 6 mo | Exact p, 6 mo | 12-mo Veh n | 12-mo Veh mean ± SEM | 12-mo Veh median | 12-mo VU01 n | 12-mo VU01 mean ± SEM | 12-mo VU01 median | MW U; p, 12 mo |
| --- | --- | --- | --- | --- | --- | --- | --- | --- | --- | --- | --- | --- | --- | --- | --- | --- | --- |
| A | Number of All Spines | Dendritic segment | 60 | 4.367 ± 0.2481 | 4 | 75 | 6 ± 0.2873 | 5 | 1409 | 0.0001 | 58 | 4.724 ± 0.2866 | 4 | 62 | 7.306 ± 0.5211 | 6.5 | 1047; <0.0001 |
| B | Stubby Spines | Dendritic segment | 12 | 0 ± 0 | 0 | 22 | 0 ± 0 | 0 | 132 | >0.9999 | 2 | 0 ± 0 | 0 | 2 | 0 ± 0 | 0 | 2; >0.9999 |
| C | Filamentous Spines | Dendritic segment | 12 | 3.25 ± 0.3917 | 3 | 22 | 3.955 ± 0.3119 | 4 | 96 | 0.1915 | 2 | 1 ± 1 | 1 | 2 | 0.5 ± 0.5 | 0.5 | 1.5; >0.9999 |
| D | Thin Spines | Dendritic segment | 12 | 0.75 ± 0.25 | 0.5 | 22 | 0.9091 ± 0.2173 | 1 | 123 | 0.7476 | 2 | 1.500 ± 1.500 | 1.5 | 2 | 1.500 ± 1.500 | 1.5 | 2; >0.9999 |
| E | Mushroom Spines | Mushroom spine | 12 | 1.417 ± 0.4167 | 1 | 22 | 2.545 ± 0.3647 | 2 | 70.5 | 0.0121 | 10 | 1.6 ± 0.4 | 1 | 17 | 1.176 ± 0.1282 | 1 | 69.5; 0.2453 |
| F | Spine Density per 10 μm | Dendritic segment | 60 | 0.4367 ± 0.02481 | 0.4 | 75 | 0.6 ± 0.02873 | 0.5 | 1409 | 0.0001 | 75 | 0.3653 ± 0.0319 | 0.4 | 75 | 0.604 ± 0.0537 | 0.5 | 1911; 0.0006 |
| G | Individual Spine Area | Individual spine | 261 | 0.2109 ± 0.0131 | 0.137 | 458 | 0.2341 ± 0.01184 | 0.134 | 58118 | 0.5375 | 274 | 0.8218 ± 0.0451 | 0.618 | 452 | 0.6455 ± 0.0241 | 0.49 | 52747; 0.0008 |
| H | Total Dendritic Area | Dendritic segment | 60 | 6.098 ± 0.2112 | 5.813 | 75 | 6.495 ± 0.2061 | 5.852 | 1955 | 0.1917 | 58 | 28.61 ± 1.842 | 27.37 | 62 | 39.6 ± 2.428 | 35.29 | 1158; 0.0007 |
| I | Spine Perimeter | Individual spine | 262 | 1.849 ± 0.0643 | 1.572 | 450 | 1.882 ± 0.0543 | 1.502 | 57052 | 0.4731 | 274 | 3.331 ± 0.0882 | 3.049 | 453 | 3.014 ± 0.0624 | 2.728 | 54150; 0.0039 |
| J | Dendrite Major Axis | Dendritic segment | 262 | 0.7978 ± 0.0255 | 0.684 | 450 | 0.7999 ± 0.0215 | 0.643 | 56990 | 0.4589 | 274 | 1.234 ± 0.0326 | 1.122 | 453 | 1.138 ± 0.0238 | 1.027 | 55617; 0.0189 |
| K | Dendrite Minor Axis | Dendritic segment | 262 | 0.2504 ± 0.0138 | 0.1995 | 450 | 0.277 ± 0.012 | 0.2005 | 57427 | 0.5606 | 274 | 0.7791 ± 0.0203 | 0.732 | 453 | 0.693 ± 0.0136 | 0.651 | 52591; 0.0006 |
| L | Mushroom Spine Head Width | Mushroom spine | 36 | 0.7136 ± 0.05836 | 0.6955 | 108 | 0.7820 ± 0.03974 | 0.6840 | 1792 | 0.4840 | 34 | 0.972 ± 0.08392 | 0.9755 | 42 | 1.161 ± 0.06455 | 1.099 | 534.5; 0.0608 |
| M | Mushroom Spine Neck Length | Mushroom Spine | 36 | 0.3419 ± 0.0426 | 0.304 | 108 | 0.3818 ± 0.0273 | 0.299 | 1812 | 0.5432 | 34 | 0.5346 ± 0.0694 | 0.398 | 42 | 0.6653 ± 0.0497 | 0.6185 | 534.5; 0.0608 |

**Supplementary Table S3B.**
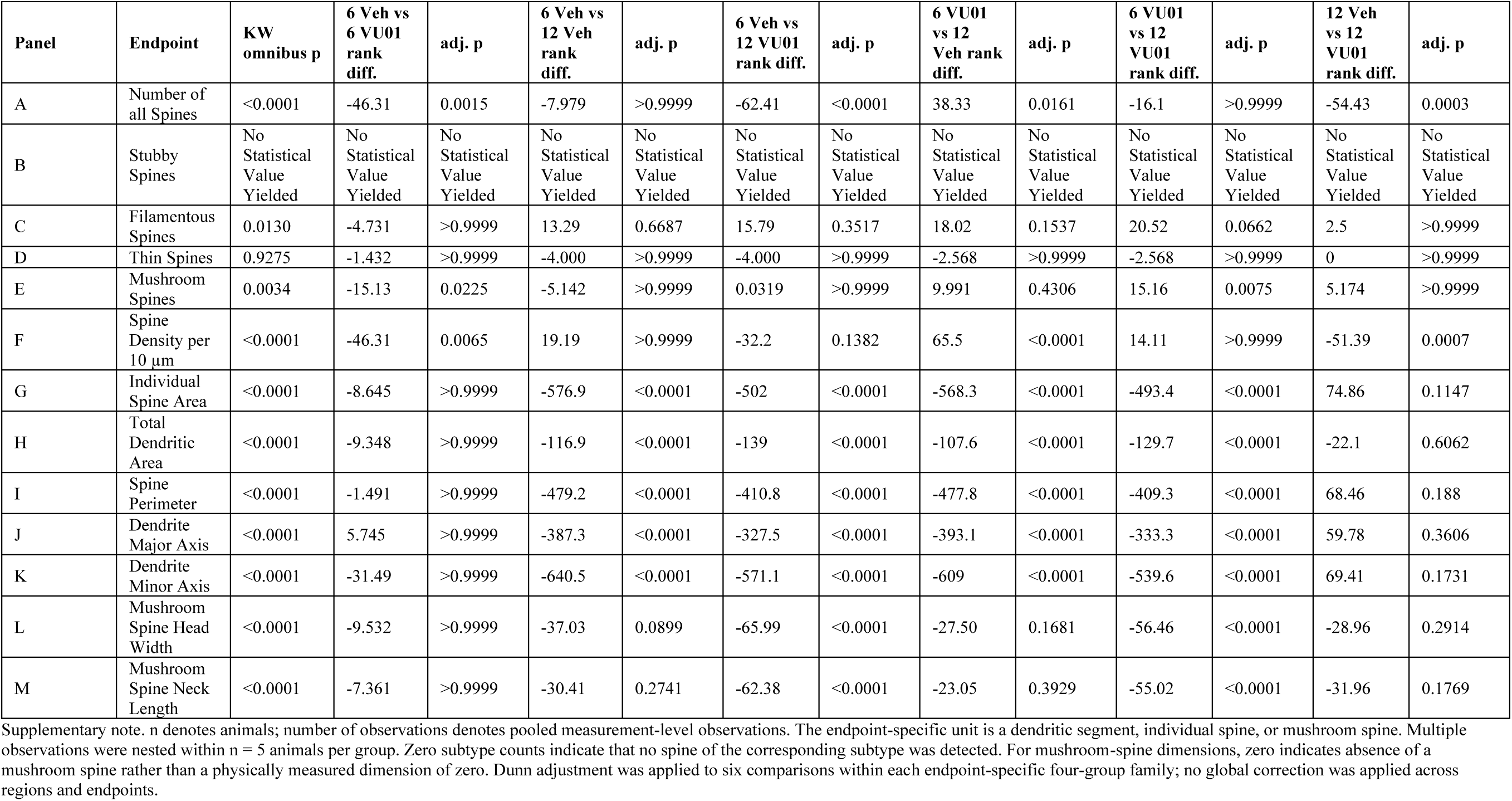
Kruskal–Wallis and Dunn-adjusted comparisons for V1. Group differences were evaluated using the Kruskal-Wallis test. Statistical measures, abbreviations, and parameters are described in Supplementary Table S1B.

| Panel | Endpoint | KW omnibus p | 6 Veh vs 6 VU01 rank diff. | adj. p | 6 Veh vs 12 Veh rank diff. | adj. p | 6 Veh vs 12 VU01 rank diff. | adj. p | 6 VU01 vs 12 Veh rank diff. | adj. p | 6 VU01 vs 12 VU01 rank diff. | adj. p | 12 Veh vs 12 VU01 rank diff. | adj. p |
| --- | --- | --- | --- | --- | --- | --- | --- | --- | --- | --- | --- | --- | --- | --- |
| A | Number of all Spines | <0.0001 | -46.31 | 0.0015 | -7.979 | >0.9999 | -62.41 | <0.0001 | 38.33 | 0.0161 | -16.1 | >0.9999 | -54.43 | 0.0003 |
| B | Stubby Spines | No Statistical Value Yielded | No Statistical Value Yielded | No Statistical Value Yielded | No Statistical Value Yielded | No Statistical Value Yielded | No Statistical Value Yielded | No Statistical Value Yielded | No Statistical Value Yielded | No Statistical Value Yielded | No Statistical Value Yielded | No Statistical Value Yielded | No Statistical Value Yielded | No Statistical Value Yielded |
| C | Filamentous Spines | 0.0130 | -4.731 | >0.9999 | 13.29 | 0.6687 | 15.79 | 0.3517 | 18.02 | 0.1537 | 20.52 | 0.0662 | 2.5 | >0.9999 |
| D | Thin Spines | 0.9275 | -1.432 | >0.9999 | -4.000 | >0.9999 | -4.000 | >0.9999 | -2.568 | >0.9999 | -2.568 | >0.9999 | 0 | >0.9999 |
| E | Mushroom Spines | 0.0034 | -15.13 | 0.0225 | -5.142 | >0.9999 | 0.0319 | >0.9999 | 9.991 | 0.4306 | 15.16 | 0.0075 | 5.174 | >0.9999 |
| F | Spine Density per 10 μm | <0.0001 | -46.31 | 0.0065 | 19.19 | >0.9999 | -32.2 | 0.1382 | 65.5 | <0.0001 | 14.11 | >0.9999 | -51.39 | 0.0007 |
| G | Individual Spine Area | <0.0001 | -8.645 | >0.9999 | -576.9 | <0.0001 | -502 | <0.0001 | -568.3 | <0.0001 | -493.4 | <0.0001 | 74.86 | 0.1147 |
| H | Total Dendritic Area | <0.0001 | -9.348 | >0.9999 | -116.9 | <0.0001 | -139 | <0.0001 | -107.6 | <0.0001 | -129.7 | <0.0001 | -22.1 | 0.6062 |
| I | Spine Perimeter | <0.0001 | -1.491 | >0.9999 | -479.2 | <0.0001 | -410.8 | <0.0001 | -477.8 | <0.0001 | -409.3 | <0.0001 | 68.46 | 0.188 |
| J | Dendrite Major Axis | <0.0001 | 5.745 | >0.9999 | -387.3 | <0.0001 | -327.5 | <0.0001 | -393.1 | <0.0001 | -333.3 | <0.0001 | 59.78 | 0.3606 |
| K | Dendrite Minor Axis | <0.0001 | -31.49 | >0.9999 | -640.5 | <0.0001 | -571.1 | <0.0001 | -609 | <0.0001 | -539.6 | <0.0001 | 69.41 | 0.1731 |
| L | Mushroom Spine Head Width | <0.0001 | -9.532 | >0.9999 | -37.03 | 0.0899 | -65.99 | <0.0001 | -27.50 | 0.1681 | -56.46 | <0.0001 | -28.96 | 0.2914 |
| M | Mushroom Spine Neck Length | <0.0001 | -7.361 | >0.9999 | -30.41 | 0.2741 | -62.38 | <0.0001 | -23.05 | 0.3929 | -55.02 | <0.0001 | -31.96 | 0.1769 |

Supplementary note. n denotes animals; number of observations denotes pooled measurement-level observations. The endpoint-specific unit is a dendritic segment, individual spine, or mushroom spine. Multiple observations were nested within n = 5 animals per group. Zero subtype counts indicate that no spine of the corresponding subtype was detected. For mushroom-spine dimensions, zero indicates absence of a mushroom spine rather than a physically measured dimension of zero. Dunn adjustment was applied to six comparisons within each endpoint-specific four-group family; no global correction was applied across regions and endpoints.

**Supplementary Table S4A.** Descriptive statistics and within-age Mann–Whitney results for V2. This table summarizes the Mann- Whitney U-test statistics, medians and means ± SEM for the V2 region. All statistical measures, abbreviations and parameters follow the same conventions as Supplementary Table S1A.

| Panel | Endpoint | Level | 6 Veh<br>n | 6 Veh<br>mean ±<br>SEM | 6 Veh<br>median | 6 VU01<br>n | 6 VU01<br>mean ±<br>SEM | 6 VU01<br>median | U 6<br>mo | p 6 mo | 12 Veh<br>n | 12 Veh<br>mean ±<br>SEM | 12 Veh<br>median | 12 VU01<br>n | 12 VU01<br>mean ±<br>SEM | 12 VU01<br>median | U; p 12 mo |
| --- | --- | --- | --- | --- | --- | --- | --- | --- | --- | --- | --- | --- | --- | --- | --- | --- | --- |
| A | Number of all Spines | Dendritic segment | 60 | 4.917 ± 0.2692 | 5 | 60 | 5.383 ± 0.3127 | 5 | 1610 | 0.3147 | 75 | 6.227 ± 0.3064 | 6 | 75 | 8.28 ± 0.3345 | 8 | 1692; <0.0001 |
| B | Stubby Spines | Dendritic segment | 13 | 0 ± 0 | 0 | 17 | 0 ± 0 | 0 | 110.5 | >0.9999 | 35 | 0 ± 0 | 0 | 55 | 0.0182 ± 0.0182 | 0 | 945; >0.9999 |
| C | Filamentous Spines | Dendritic segment | 13 | 3.077 ± 0.5484 | 3 | 17 | 3.412 ± 0.582 | 3 | 102.5 | 0.7462 | 35 | 4.4 ± 0.4042 | 4 | 55 | 2.4 ± 0.4277 | 0 | 541; 0.0003 |
| D | Thin Spines | Dendritic segment | 13 | 0.6154 ± 0.3846 | 0 | 17 | 0.9412 ± 0.2344 | 1 | 77 | 0.1292 | 35 | 0.5429 ± 0.1184 | 0 | 55 | 3.164 ± 0.427 | 2 | 445; <0.0001 |
| E | Mushroom Spines | Mushroom spine | 13 | 1.615 ± 0.3309 | 1 | 19 | 1.789 ± 0.292 | 1 | 109.5 | 0.5608 | 35 | 2.429 ± 0.2537 | 2 | 55 | 2.255 ± 0.2159 | 2 | 901.5; 0.6077 |
| F | Spine Density per 10 μm | Dendritic segment | 60 | 0.4917 ± 0.0269 | 0.5 | 71 | 0.5408 ± 0.0285 | 0.5 | 1884 | 0.2528 | 75 | 0.6227 ± 0.0306 | 0.6 | 75 | 0.828 ± 0.0335 | 0.8 | 1692; <0.0001 |
| G | Individual Spine Area | Individual spine | 295 | 0.183 ± 0.0101 | 0.125 | 384 | 0.2584 ± 0.0169 | 0.125 | 54190 | 0.3333 | 466 | 0.2583 ± 0.0136 | 0.148 | 621 | 0.2496 ± 0.0116 | 0.142 | 137272; 0.1473 |
| H | Total Dendritic Area | Dendritic segment | 60 | 5.943 ± 0.1796 | 5.631 | 71 | 9.523 ± 1.101 | 5.624 | 1898 | 0.2844 | 75 | 6.756 ± 0.1952 | 6.348 | 75 | 7.156 ± 0.273 | 7.083 | 2619; 0.4687 |
| I | Spine Perimeter | Individual spine | 295 | 1.723 ± 0.052 | 1.528 | 323 | 1.645 ± 0.0628 | 1.27 | 41593 | 0.0063 | 467 | 2.065 ± 0.06 | 1.697 | 621 | 1.953 ± 0.0524 | 1.528 | 134679; 0.0441 |
| J | Dendrite Major Axis | Dendritic segment | 295 | 0.7615 ± 0.0225 | 0.679 | 384 | 0.7755 ± 0.0244 | 0.643 | 54991 | 0.515 | 467 | 0.8766 ± 0.0237 | 0.755 | 629 | 0.8199 ± 0.0202 | 0.679 | 135893; 0.0341 |
| K | Dendrite Minor Axis | Dendritic segment | 295 | 0.2122 ± 0.0104 | 0.151 | 384 | 0.3066 ± 0.0146 | 0.201 | 46788 | <0.0001 | 467 | 0.2974 ± 0.012 | 0.213 | 629 | 0.3014 ± 0.0108 | 0.209 | 145566; 0.7998 |
| L | Mushroom Spine Head Width | Mushroom spine | 42 | 0.7088 ± 0.05383 | 0.647 | 64 | 0.8699 ± 0.06773 | 0.7880 | 1118 | 0.1443 | 170 | 0.7556 ± 0.03412 | 0.649 | 246 | 0.7694 ± 0.03024 | 0.649 | 20598; 0.7957 |
| M | Mushroom Spine Neck Length | Mushroom spine | 42 | 0.3439 ± 0.0401 | 0.3345 | 64 | 0.4087 ± 0.0352 | 0.4135 | 1186 | 0.3071 | 170 | 0.4086 ± 0.0196 | 0.4025 | 246 | 0.408 ± 0.0174 | 0.368 | 20558; 0.7705 |

**Supplementary Table S4B.** Kruskal–Wallis and Dunn-adjusted comparisons for V2. The Kruskal-Wallis test was used to observe differences among groups. For comprehensive definitions of statistical measures, abbreviations and parameters, see Supplementary Table S1B.

| Panel | Endpoint | KW p | 6 Veh vs<br>6 VU01<br>rank<br>diff. | adj. p | 6 Veh vs<br>12 Veh<br>rank<br>diff. | adj. p | 6 Veh vs<br>12 VU01<br>rank<br>diff. | adj. p | 6 VU01<br>vs 12<br>Veh<br>rank<br>diff. | adj. p | 6 VU01<br>vs 12<br>VU01<br>rank<br>diff. | adj. p | 12 Veh<br>vs 12<br>VU01<br>rank<br>diff. | adj. p |
| --- | --- | --- | --- | --- | --- | --- | --- | --- | --- | --- | --- | --- | --- | --- |
| A | Number of<br>all Spines | <0.0001 | -14.3 | >0.9999 | -37.95 | 0.0286 | -89.81 | <0.0001 | -23.65 | 0.4712 | -75.51 | <0.0001 | -51.85 | 0.0003 |
| B | Stubby<br>Spines | 0.7574 | 0 | >0.9999 | 0 | >0.9999 | -1.091 | >0.9999 | 0 | >0.9999 | -1.091 | >0.9999 | -1.091 | >0.9999 |
| C | Filamentous<br>Spines | 0.0015 | -2.219 | >0.9999 | -14.22 | >0.9999 | 14.52 | >0.9999 | -12 | >0.9999 | 16.74 | 0.4708 | 28.74 | 0.0006 |
| D | Thin Spines | <0.0001 | -14.83 | >0.9999 | -4.197 | >0.9999 | -36.72 | 0.0021 | 10.63 | >0.9999 | -21.89 | 0.1061 | -32.52 | <0.0001 |
| E | Mushroom<br>Spines | 0.1526 | -5.358 | >0.9999 | -21 | 0.3388 | -16.69 | 0.6616 | -15.64 | 0.6323 | -11.34 | >0.9999 | 4.305 | >0.9999 |
| F | Spine<br>Density per<br>10 μm | <0.0001 | -16.01 | >0.9999 | -39.59 | 0.0279 | -93.59 | <0.0001 | -23.58 | 0.4676 | -77.57 | <0.0001 | -53.99 | 0.0003 |
| G | Individual<br>Spine Area | 0.0137 | -45.17 | >0.9999 | -118.9 | 0.0104 | -71.32 | 0.2873 | -73.69 | 0.2158 | -26.15 | >0.9999 | 47.54 | 0.7688 |
| H | Total<br>Dendritic<br>Area | 0.0094 | -18.46 | >0.9999 | -37.43 | 0.047 | -42.89 | 0.0139 | -18.97 | 0.9519 | -24.43 | 0.4168 | -5.46 | >0.9999 |
| I | Spine<br>Perimeter | <0.0001 | 97.92 | 0.0812 | -102.7 | 0.0302 | -39.46 | >0.9999 | -200.6 | <0.0001 | -137.4 | 0.0003 | 63.25 | 0.2158 |
| J | Dendrite<br>Major Axis | 0.0109 | 22.83 | >0.9999 | -88.02 | 0.1253 | -20.51 | >0.9999 | -110.8 | 0.0101 | -43.34 | >0.9999 | 67.51 | 0.1858 |
| K | Dendrite<br>Minor Axis | <0.0001 | -160 | 0.0003 | -193.1 | <0.0001 | -183.8 | <0.0001 | -33.11 | >0.9999 | -23.84 | >0.9999 | 9.267 | >0.9999 |
| L | Mushroom<br>Spine Head<br>Width | 0.4231 | -38.03 | >0.9999 | -3.292 | >0.9999 | -7.130 | >0.9999 | 34.74 | 0.6974 | 30.90 | 0.8652 | -3.837 | >0.9999 |
| M | Mushroom<br>Spine Neck<br>Length | 0.6308 | -32.65 | >0.9999 | -33.34 | >0.9999 | -28.92 | >0.9999 | -0.6859 | >0.9999 | 3.735 | >0.9999 | 4.421 | >0.9999 |

Supplementary note. n denotes animals; number of observations denotes pooled measurement-level observations. Multiple observations were nested within n = 5 animals per group. Dunn adjustment was applied within each endpoint-specific six-comparison family; no global correction was applied across regions and endpoints.

**Supplementary Table S5A.** Descriptive statistics and within-age Mann–Whitney results for CA2. Mann-Whitney U-test statistics, median and mean ± SEM values are outlined within this table. As mentioned previously, all statistical measures, abbreviations, and parameters mirror the formatting of Supplementary Table S1A.

| Panel | Endpoint | Level | 6 Veh<br>n | 6 Veh<br>mean ±<br>SEM | 6 Veh<br>median | 6<br>VU01<br>n | 6<br>VU01<br>mean ±<br>SEM | 6 VU01<br>median | U 6 mo | p 6 mo | 12<br>Veh<br>n | 12 Veh<br>mean ±<br>SEM | 12 Veh<br>median | 12<br>VU01<br>n | 12<br>VU01<br>mean ±<br>SEM | 12<br>VU01<br>median | U; p 12<br>mo |
| --- | --- | --- | --- | --- | --- | --- | --- | --- | --- | --- | --- | --- | --- | --- | --- | --- | --- |
| A | Number of<br>all Spines | Dendritic<br>segment | 71 | 5.577 ±<br>0.2987 | 5 | 75 | 7.467 ±<br>0.311 | 7 | 1614 | <0.0001 | 72 | 8.458 ±<br>0.3716 | 8 | 75 | 15.29 ±<br>0.3263 | 14.82 | 268;<br><0.0001 |
| B | Stubby<br>Spines | Dendritic<br>segment | 48 | 0.0417<br>±<br>0.0417 | 0 | 54 | 0 ± 0 | 0 | 1269 | 0.4706 | 16 | 0.0625<br>±<br>0.0625 | 0 | 42 | 0 ± 0 | 0 | 315;<br>0.2759 |
| C | Filamentous<br>Spines | Dendritic<br>segment | 50 | 2.3 ±<br>0.2395 | 2 | 56 | 3.411 ±<br>0.2947 | 3 | 999 | 0.0098 | 16 | 2.688 ±<br>0.5896 | 2.5 | 42 | 3.143 ±<br>0.3002 | 3 | 279.5;<br>0.3234 |
| D | Thin Spines | Dendritic<br>segment | 50 | 0.94 ±<br>0.1839 | 0.5 | 56 | 1.411 ±<br>0.1798 | 1 | 1051 | 0.0204 | 17 | 3.412 ±<br>0.6477 | 3 | 42 | 1.667 ±<br>0.2632 | 1 | 212.5;<br>0.0129 |
| E | Mushroom<br>Spines | Mushroom<br>spine | 50 | 2.82 ±<br>0.2014 | 3 | 56 | 2.911 ±<br>0.2325 | 2 | 1362 | 0.8049 | 16 | 2.313 ±<br>0.4155 | 2 | 42 | 2.095 ±<br>0.215 | 2 | 318.5;<br>0.7544 |
| F | Spine<br>Density per<br>10 µm | Dendritic<br>segment | 70 | 0.5629<br>±<br>0.0299 | 0.5 | 75 | 0.7467<br>±<br>0.0311 | 0.7 | 1613 | <0.0001 | 72 | 0.8458<br>±<br>0.0372 | 0.8 | 75 | 0.5787<br>±<br>0.0332 | 0.6 | 1429;<br><0.0001 |
| G | Individual<br>Spine Area | Individual<br>spine | 405 | 0.3078<br>±<br>0.0235 | 0.188 | 560 | 0.2594<br>±<br>0.0116 | 0.16 | 105604 | 0.068 | 609 | 0.2243<br>±<br>0.0092 | 0.154 | 435 | 0.3365<br>±<br>0.0371 | 0.199 | 107696;<br><0.0001 |
| H | Total<br>Dendritic<br>Area | Dendritic<br>segment | 75 | 8.876 ±<br>0.3023 | 8.667 | 75 | 8.281 ±<br>0.2595 | 8.001 | 2473 | 0.2029 | 72 | 9.949 ±<br>0.3605 | 9.368 | 75 | 9.906 ±<br>0.41 | 9.055 | 2589;<br>0.6687 |
| I | Spine<br>Perimeter | Individual<br>spine | 1084 | 1.825 ±<br>0.0443 | 1.454 | 550 | 1.945 ±<br>0.0526 | 1.635 | 273137 | 0.0056 | 602 | 1.764 ±<br>0.0409 | 1.484 | 433 | 2.249 ±<br>0.0825 | 1.786 | 104212;<br><0.0001 |
| J | Dendrite<br>Major Axis | Dendritic<br>segment | 433 | 0.9044<br>±<br>0.0386 | 0.759 | 550 | 0.7945<br>±<br>0.0203 | 0.679 | 108477 | 0.0165 | 602 | 0.7305<br>±<br>0.0169 | 0.609 | 477 | 0.9888<br>±<br>0.0395 | 0.759 | 114450;<br><0.0001 |
| K | Dendrite<br>Minor Axis | Dendritic<br>segment | 433 | 0.3678<br>±<br>0.0135 | 0.283 | 550 | 0.3397<br>±<br>0.0108 | 0.277 | 113360 | 0.1949 | 602 | 0.3362<br>±<br>0.0083 | 0.28 | 477 | 0.4924<br>±<br>0.0308 | 0.323 | 130817;<br>0.0121 |
| L | Mushroom<br>Spine Head<br>Width | Mushroom<br>spine | 310 | 0.6404<br>±<br>0.0234 | 0.539 | 318 | 0.6465<br>±<br>0.0212 | 0.55 | 47552 | 0.4444 | 78 | 0.7161<br>±<br>0.0424 | 0.6335 | 435 | 0.9234<br>±<br>0.0336 | 0.755 | 13643;<br>0.0057 |
| M | Mushroom<br>Spine Neck<br>Length | Mushroom<br>spine | 276 | 0.3468<br>±<br>0.0145 | 0.311 | 318 | 0.3593<br>±<br>0.0123 | 0.338 | 41442 | 0.2415 | 78 | 0.3966<br>±<br>0.03025 | 0.334 | 182 | 0.3295<br>±<br>0.02491 | 0.279 | 5855;<br>0.0241 |

**Supplementary Table S5B.** Kruskal–Wallis and Dunn-adjusted comparisons for CA2. For determination of significant variations across groups, Kruskal-Wallis test was applied. All statistical measurements and parametric details can be referenced in Supplementary Table S1B.

| Panel | Endpoint | KW p | 6 Veh vs<br>6 VU01<br>rank<br>diff. | adj. p | 6 Veh vs<br>12 Veh<br>rank<br>diff. | adj. p | 6 Veh vs<br>12 VU01<br>rank<br>diff. | adj. p | 6 VU01<br>vs 12<br>Veh<br>rank<br>diff. | adj. p | 6 VU01<br>vs 12<br>VU01<br>rank<br>diff. | adj. p | 12 Veh<br>vs 12<br>VU01<br>rank<br>diff. | adj. p |
| --- | --- | --- | --- | --- | --- | --- | --- | --- | --- | --- | --- | --- | --- | --- |
| A | Number of<br>all Spines | <0.0001 | -41.89 | 0.0166 | -62.74 | <0.0001 | -174.3 | <0.0001 | -20.85 | 0.8102 | -132.4 | <0.0001 | -111.5 | <0.0001 |
| B | Stubby<br>Spines | 0.1989 | 1.677 | >0.9999 | -3.292 | >0.9999 | 1.677 | >0.9999 | -4.969 | 0.3015 | 0 | >0.9999 | 4.969 | 0.3471 |
| C | Filamentous<br>Spines | 0.0461 | -23.79 | 0.0546 | -5.906 | >0.9999 | -19.71 | 0.2674 | 17.89 | >0.9999 | 4.08 | >0.9999 | -13.81 | >0.9999 |
| D | Thin Spines | 0.0004 | -18.68 | 0.2261 | -53.68 | 0.0002 | -22.23 | 0.129 | -35 | 0.0374 | -3.551 | >0.9999 | 31.44 | 0.1074 |
| E | Mushroom<br>Spines | 0.0198 | 1.162 | >0.9999 | 20.62 | 0.7207 | 25.47 | 0.0506 | 19.46 | 0.8235 | 24.3 | 0.0596 | 4.848 | >0.9999 |
| F | Spine<br>Density per<br>10 μm | <0.0001 | -53.47 | 0.0008 | -77.52 | <0.0001 | -6.759 | >0.9999 | -24.05 | 0.4961 | 46.71 | 0.004 | 70.76 | <0.0001 |
| G | Individual<br>Spine Area | <0.0001 | 71.11 | 0.361 | 111 | 0.017 | -70.59 | 0.4678 | 39.91 | >0.9999 | -141.7 | 0.0008 | -181.6 | <0.0001 |
| H | Total<br>Dendritic<br>Area | 0.0063 | 17.16 | >0.9999 | -28 | 0.2887 | -19.63 | 0.9692 | -45.16 | 0.0086 | -36.79 | 0.0522 | 8.37 | >0.9999 |
| I | Spine<br>Perimeter | <0.0001 | -114.1 | 0.0281 | -25.3 | >0.9999 | -285.7 | <0.0001 | 88.78 | 0.3048 | -171.6 | 0.0032 | -260.4 | <0.0001 |
| J | Dendrite<br>Major Axis | <0.0001 | 94.08 | 0.0834 | 159.3 | 0.0001 | -49.16 | >0.9999 | 65.26 | 0.3785 | -143.2 | 0.0007 | -208.5 | <0.0001 |
| K | Dendrite<br>Minor Axis | 0.0038 | 51.99 | >0.9999 | 1.734 | >0.9999 | -83.46 | 0.2076 | -50.26 | 0.9129 | -135.5 | 0.0016 | -85.19 | 0.117 |
| L | Mushroom<br>Spine Head<br>Width | <0.0001 | -16.36 | >0.9999 | -91.77 | 0.1673 | -198.1 | <0.0001 | -75.41 | 0.4204 | -181.7 | <0.0001 | -106.3 | 0.0522 |
| M | Mushroom<br>Spine Neck<br>Length | 0.0680 | -20.75 | >0.9999 | -34.47 | >0.9999 | 34.27 | 0.8707 | -13.72 | >0.9999 | 55.01 | 0.0976 | 68.74 | 0.2352 |

Supplementary note. n denotes animals; number of observations denotes pooled measurement-level observations. Multiple observations were nested within n = 5 animals per group. NR means not reported or not recoverable without invention. Dunn adjustment was applied within each endpoint-specific six-comparison family; no global correction was applied across regions and endpoints.

**Supplementary Table S6A.**
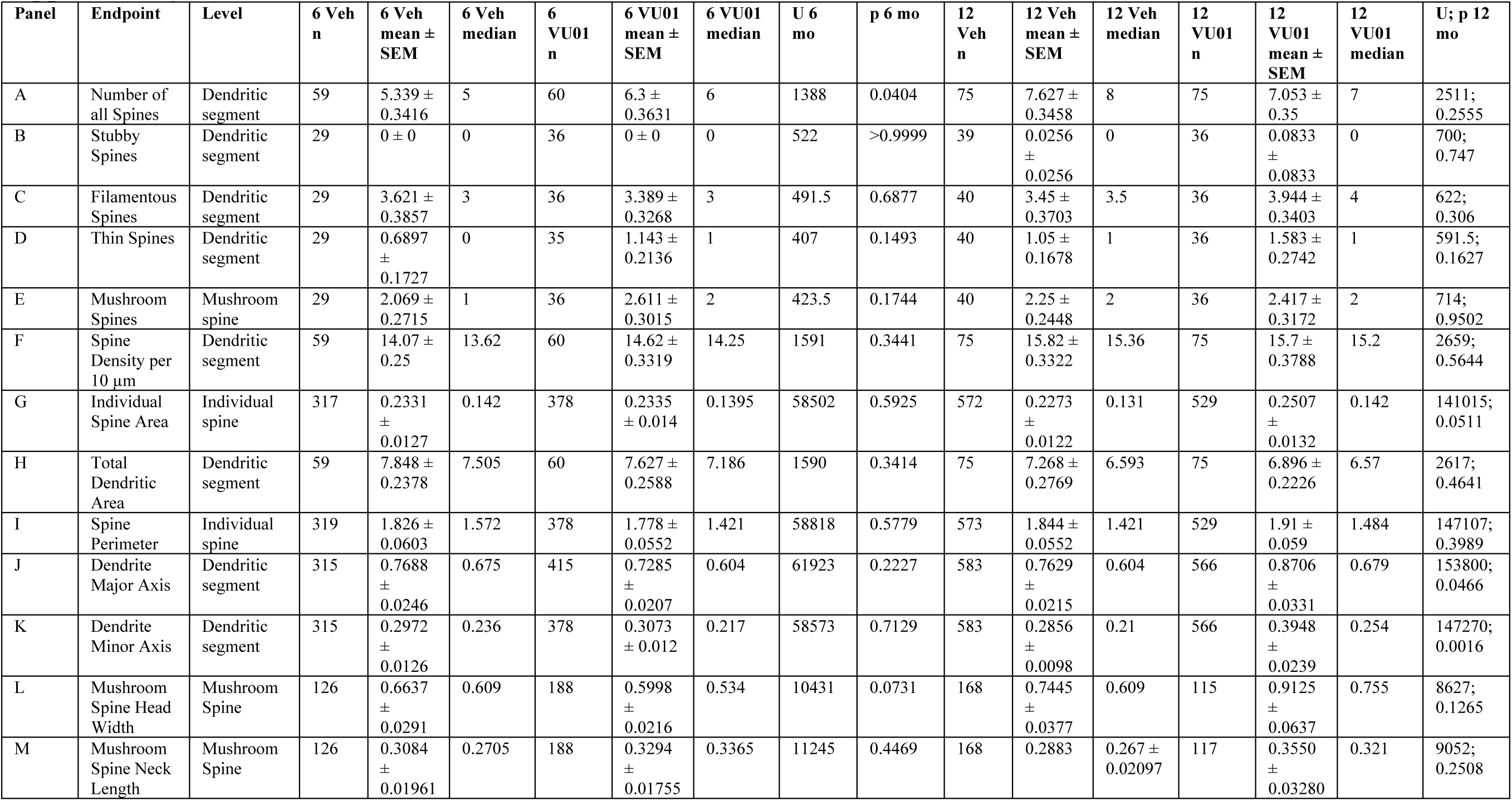
Descriptive statistics and within-age Mann–Whitney results for CA3. Statistical analysis in the CA3 region was done using the Mann-Whitney test yielding U-test statistics, medians, and means ± SEM. All details of statistical measures, abbreviations and parameters can be found in Supplementary Table S1A.

**Supplementary Table S6B.**
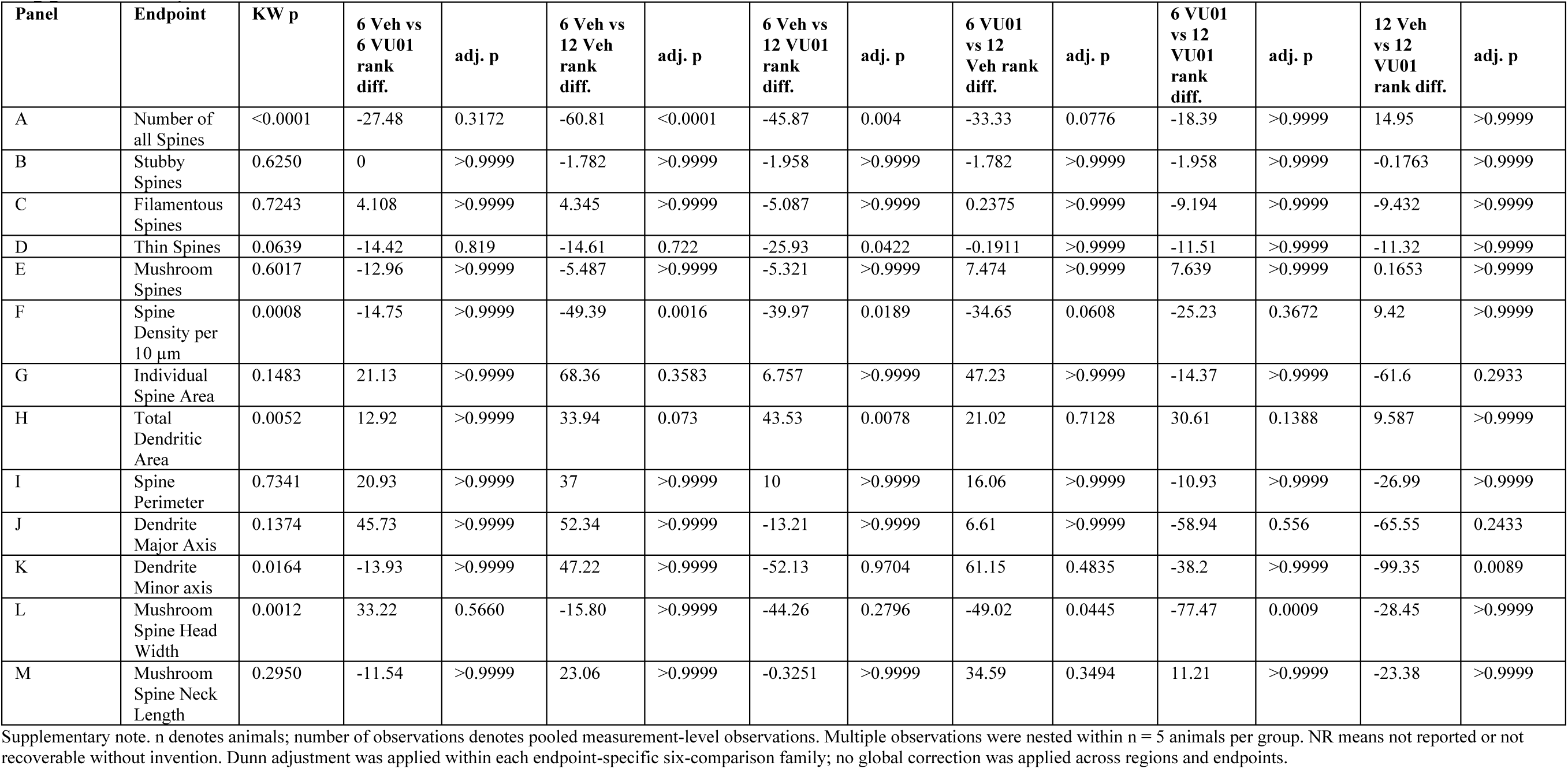
Kruskal–Wallis and Dunn-adjusted comparisons for CA3. Regional Kruskal-Wallis analysis of the CA3 region was conducted to observe for inter-group differences with all similar statistical notations and parameters as Supplementary Table S1B.

**Supplementary Table S7A.** Descriptive statistics and within-age Mann–Whitney results for DG. Significant pairwise differences within the V2 region were determined using Mann-Whitney U-tests alongside reporting of medians, means ± SEM. All statistical notations and parameters follow the Supplementary Table S1A schema.

| Panel | Endpoint | Level | 6 Veh<br>n | 6 Veh<br>mean ±<br>SEM | 6 Veh<br>median | 6 VU01<br>n | 6 VU01<br>mean ±<br>SEM | 6 VU01<br>median | U 6<br>mo | p 6 mo | 12 Veh<br>n | 12 Veh<br>mean ±<br>SEM | 12 Veh<br>median | 12 VU01<br>n | 12 VU01<br>mean ±<br>SEM | 12 VU01<br>median | U; p 12 mo |
| --- | --- | --- | --- | --- | --- | --- | --- | --- | --- | --- | --- | --- | --- | --- | --- | --- | --- |
| A | Number of all Spines | Dendritic segment | 70 | 5.286 ± 0.3173 | 5 | 75 | 6.467 ± 0.3339 | 6 | 2051 | 0.022 | 75 | 6.267 ± 0.4055 | 6 | 75 | 8.987 ± 0.4243 | 9 | 1650; <0.0001 |
| B | Stubby Spines | Dendritic segment | 34 | 0 ± 0 | 0 | 42 | 0 ± 0 | 0 | 1309 | >0.9999 | 2 | 0 ± 0 | 0 | 3 | 0 ± 0 | 0 | 3; >0.9999 |
| C | Filamentous Spines | Dendritic segment | 34 | 3.324 ± 0.2694 | 3 | 42 | 4.048 ± 0.2931 | 4 | 563.5 | 0.1106 | 2 | 5.5 ± 0.5 | 5.5 | 3 | 8.667 ± 0.8819 | 9 | 0; 0.2 |
| D | Thin Spines | Dendritic segment | 34 | 1.147 ± 0.1941 | 1 | 42 | 1.095 ± 0.1797 | 1 | 690 | 0.8019 | 2 | 2 ± 2 | 2 | 3 | 3.667 ± 1.453 | 4 | 1.5; 0.5 |
| E | Mushroom Spines | Mushroom spine | 34 | 2.265 ± 0.1903 | 2 | 42 | 2.19 ± 0.2267 | 1.5 | 630.5 | 0.3642 | 3 | 1.333 ± 0.3333 | 1 | 3 | 1.333 ± 0.3333 | 1 | 4.5; >0.9999 |
| F | Spine Density per 10 µm | Dendritic segment | 70 | 13.81 ± 0.2143 | 13.64 | 75 | 14.62 ± 0.2545 | 14.69 | 2179 | 0.0779 | 75 | 13.82 ± 0.2295 | 13.82 | 75 | 15.4 ± 0.2676 | 15.26 | 1715; <0.0001 |
| G | Individual Spine Area | Individual spine | 370 | 0.2068 ± 0.0113 | 0.1395 | 483 | 0.1934 ± 0.0081 | 0.125 | 87501 | 0.603 | 470 | 0.159 ± 0.0058 | 0.1225 | 674 | 0.1682 ± 0.0051 | 0.131 | 151035; 0.1808 |
| H | Total Dendritic Area | Dendritic segment | 69 | 7.184 ± 0.255 | 6.593 | 75 | 6.587 ± 0.2158 | 5.972 | 2111 | 0.0564 | 75 | 6.742 ± 0.192 | 6.473 | 75 | 6.965 ± 0.1775 | 6.713 | 2525; 0.2812 |
| I | Spine Perimeter | Individual spine | 360 | 1.73 ± 0.0538 | 1.484 | 507 | 1.706 ± 0.0421 | 1.421 | 91061 | 0.9563 | 470 | 1.439 ± 0.031 | 1.289 | 673 | 1.462 ± 0.0267 | 1.333 | 152812; 0.3305 |
| J | Dendrite Major Axis | Dendritic segment | 360 | 0.7323 ± 0.0234 | 0.609 | 484 | 0.7216 ± 0.0181 | 0.609 | 86507 | 0.861 | 470 | 0.2794 ± 0.0062 | 0.256 | 667 | 0.2982 ± 0.0117 | 0.252 | 155899; 0.8767 |
| K | Dendrite Minor Axis | Dendritic segment | 360 | 0.2746 ± 0.0104 | 0.226 | 484 | 0.2662 ± 0.009 | 0.2105 | 83922 | 0.3579 | 470 | 0.2794 ± 0.0062 | 0.256 | 664 | 0.283 ± 0.0055 | 0.252 | 155898; 0.9791 |
| L | Mushroom Spine Head Width | Mushroom spine | 152 | 0.5172 ± 0.02229 | 0.459 | 668 | 0.6924 ± 0.02059 | 0.5900 | 37216 | <0.0001 | 14 | 1.383 ± 0.6735 | 0.6825 | 8 | 0.5401 ± 0.0657 | 0.5290 | 37.5; 0.2179 |
| M | Mushroom Spine Neck Length | Mushroom spine | 152 | 0.2288 ± 0.01552 | 0.2090 | 184 | 0.2905 ± 0.01496 | 0.2670 | 11614 | 0.0074 | 14 | 0.4681 ± 0.1140 | 0.3360 | 8 | 0.4145 ± 0.05420 | 0.4310 | 51; 0.7639 |

**Supplementary Table S7B.** Kruskal–Wallis and Dunn-adjusted comparisons for DG. Group variations were quantified using the Kruskal-Wallis test, with statistical parameters that follow Supplementary Table S1A.

| Panel | Endpoint | KW p | 6 Veh vs 6 VU01 rank diff. | adj. p | 6 Veh vs 12 Veh rank diff. | adj. p | 6 Veh vs 12 VU01 rank diff. | adj. p | 6 VU01 vs 12 Veh rank diff. | adj. p | 6 VU01 vs 12 VU01 rank diff. | adj. p | 12 Veh vs 12 VU01 rank diff. | adj. p |
| --- | --- | --- | --- | --- | --- | --- | --- | --- | --- | --- | --- | --- | --- | --- |
| A | Number of all Spines | <0.0001 | -29.66 | 0.2136 | -21.83 | 0.7319 | -85.34 | <0.0001 | 7.833 | >0.9999 | -55.68 | 0.0004 | -63.51 | <0.0001 |
| B | Stubby Spines | No Statistical Value Yielded | No Statistical Value Yielded | No Statistical Value Yielded | No Statistical Value Yielded | No Statistical Value Yielded | No Statistical Value Yielded | No Statistical Value Yielded | No Statistical Value Yielded | No Statistical Value Yielded | No Statistical Value Yielded | No Statistical Value Yielded | No Statistical Value Yielded | No Statistical Value Yielded |
| C | Filamentous Spines | 0.0051 | -8.331 | 0.7152 | -28.93 | 0.5178 | -44.34 | 0.0089 | -20.6 | >0.9999 | -36.01 | 0.0559 | -15.42 | >0.9999 |
| D | Thin Spines | 0.2147 | 1.342 | >0.9999 | -6.706 | >0.9999 | -26.62 | 0.2999 | -8.048 | >0.9999 | -27.96 | 0.2279 | -19.92 | >0.9999 |
| E | Mushroom Spines | 0.2993 | 5.116 | >0.9999 | 18.65 | >0.9999 | 18.65 | >0.9999 | 13.54 | >0.9999 | 13.54 | >0.9999 | 0 | >0.9999 |
| F | Spine Density per 10 μm | <0.0001 | -27.83 | 0.2976 | 0.939 | >0.9999 | -56.94 | 0.0004 | 28.77 | 0.2332 | -29.11 | 0.22 | -57.88 | 0.0002 |
| G | Individual Spine Area | 0.1785 | 24.96 | >0.9999 | 84.47 | 0.21 | 41.24 | >0.9999 | 59.5 | 0.6667 | 16.28 | >0.9999 | -43.23 | >0.9999 |
| H | Total Dendritic Area | 0.0910 | 29.94 | 0.2086 | 14.06 | >0.9999 | -0.9957 | >0.9999 | -15.87 | >0.9999 | -30.93 | 0.1552 | -15.06 | >0.9999 |
| I | Spine Perimeter | <0.0001 | -0.4399 | >0.9999 | 151.2 | 0.0012 | 119.3 | 0.0098 | 151.6 | 0.0003 | 119.7 | 0.0027 | -31.88 | >0.9999 |
| J | Dendrite Major Axis | <0.0001 | -5.271 | >0.9999 | 813.8 | <0.0001 | 804.1 | <0.0001 | 819.1 | <0.0001 | 809.4 | <0.0001 | -9.751 | >0.9999 |
| K | Dendrite Minor Axis | <0.0001 | 45.69 | >0.9999 | -103.4 | 0.058 | -105.9 | 0.0274 | -149.1 | 0.0003 | -151.6 | <0.0001 | -2.509 | >0.9999 |
| L | Mushroom Spine Head Width | <0.0001 | -112.6 | <0.0001 | -186.0 | 0.0370 | -55.98 | >0.9999 | -73.35 | >0.9999 | 56.65 | >0.9999 | 130.0 | >0.9999 |
| M | Mushroom Spine Neck Length | 0.0003 | -30.09 | 0.0472 | -87.56 | 0.0144 | -103.6 | 0.0340 | -57.47 | 0.2686 | -73.55 | 0.2920 | -16.08 | >0.9999 |

Supplementary note. n denotes animals; number of observations denotes pooled measurement-level observations. Multiple observations were nested within n = 5 animals per group. Dunn adjustment was applied within each endpoint-specific six-comparison family; no global correction was applied across regions and endpoints.

**Supplementary Table S8A.** Descriptive statistics and within-age Mann–Whitney results for CeA. Summary metrics for CeA region yielded Mann-Whitney U-test outputs, medians and means ± SEM. All underlying statistical parameters follow the conventions utilized in Supplementary Table S1A.

| Panel | Endpoint | Level | 6 Veh<br>n | 6 Veh<br>mean ±<br>SEM | 6 Veh<br>median | 6 VU01<br>n | 6 VU01<br>mean ±<br>SEM | 6 VU01<br>median | U 6<br>mo | p 6 mo | 12 Veh n | 12 Veh<br>mean ±<br>SEM | 12 Veh<br>median | 12 VU01<br>n | 12 VU01<br>mean ±<br>SEM | 12 VU01<br>median | U; p 12<br>mo |
| --- | --- | --- | --- | --- | --- | --- | --- | --- | --- | --- | --- | --- | --- | --- | --- | --- | --- |
| A | Number of<br>all Spines | Dendritic<br>segment | 46 | 6.935 ±<br>0.4175 | 6 | 55 | 7.291 ±<br>0.3574 | 7 | 1171 | 0.5195 | 75 | 14.93 ±<br>0.3683 | 15 | 75 | 14.73 ±<br>0.3481 | 15 | 2729;<br>0.7523 |
| B | Stubby<br>Spines | Dendritic<br>segment | 46 | 6.304 ±<br>0.4134 | 6 | 55 | 3.964 ±<br>0.2321 | 4 | 620.5 | <0.0001 | 75 | 12.72 ±<br>0.4478 | 13 | 75 | 12.44 ±<br>0.3793 | 12 | 2694;<br>0.6562 |
| C | Filamentous<br>Spines | Dendritic<br>segment | 5 | 0.4 ±<br>0.2449 | 0 | 42 | 3.81 ±<br>0.2976 | 3.5 | 4 | <0.0001 | 75 | 1.24 ±<br>0.1779 | 1 | 75 | 1.4 ±<br>0.2 | 1 | 2705;<br>0.6708 |
| D | Mushroom<br>Spines | Mushroom<br>spine | 15 | 1.933 ±<br>0.1533 | 2 | 17 | 1.412 ±<br>0.1728 | 1 | 70 | 0.0212 | 75 | 0.76 ±<br>0.1481 | 0 | 75 | 0.8933<br>± 0.156 | 0 | 2594;<br>0.3528 |
| E | Spine<br>Density per<br>10 µm | Dendritic<br>segment | 46 | 0.6909<br>±<br>0.0421 | 0.6058 | 70 | 1.183 ±<br>0.113 | 0.8031 | 1153 | 0.0095 | 75 | 2.366 ±<br>0.09 | 2.264 | 75 | 2.198 ±<br>0.0812 | 2.181 | 2570;<br>0.3641 |
| F | Individual<br>Spine Area | Individual<br>spine | 317 | 0.3376<br>±<br>0.0168 | 0.251 | 401 | 0.3207<br>±<br>0.0155 | 0.222 | 60757 | 0.3101 | 1125 | 0.2145<br>±<br>0.0151 | 0.085 | 1184 | 0.2517<br>±<br>0.0121 | 0.117 | 571917;<br><0.0001 |
| G | Total<br>Dendritic<br>Area | Dendritic<br>segment | 46 | 7.292 ±<br>0.3258 | 6.69 | 55 | 7.557 ±<br>0.2679 | 7.368 | 1113 | 0.3006 | 75 | 9.592 ±<br>0.4569 | 8.457 | 75 | 10.98 ±<br>0.5032 | 9.867 | 2242;<br>0.0319 |
| H | Spine<br>Perimeter | Individual<br>spine | 321 | 2.376 ±<br>0.091 | 2.062 | 440 | 2.085 ±<br>0.0406 | 1.826 | 64301 | 0.0348 | 1125 | 1.81 ±<br>0.0548 | 1.368 | 1172 | 2.016 ±<br>0.0451 | 1.576 | 564784;<br><0.0001 |
| I | Dendrite<br>Major Axis | Dendritic<br>segment | 316 | 0.973 ±<br>0.0382 | 0.849 | 443 | 0.8842<br>±<br>0.0259 | 0.77 | 64974 | 0.0917 | 1107 | 0.7361<br>±<br>0.0222 | 0.584 | 1104 | 0.8043<br>± 0.019 | 0.6595 | 530100;<br><0.0001 |
| J | Dendrite<br>Minor Axis | Dendritic<br>segment | 316 | 0.4059<br>±<br>0.0139 | 0.346 | 441 | 0.3865<br>±<br>0.0106 | 0.334 | 68135 | 0.603 | 1107 | 0.3078<br>±<br>0.0085 | 0.206 | 1104 | 0.359 ±<br>0.0092 | 0.2455 | 534071;<br><0.0001 |
| K | Mushroom<br>Spine Head<br>Width | Mushroom<br>spine | 33 | 0.9138<br>±<br>0.0685 | 0.844 | 24 | 1.05 ±<br>0.1139 | 0.9165 | 328.5 | 0.2792 | 57 | 0.6929<br>±<br>0.0601 | 0.637 | 69 | 0.7401<br>±<br>0.0328 | 0.702 | 1655;<br>0.1268 |
| L | Mushroom<br>Spine Neck<br>Length | Mushroom<br>spine | 33 | 0.7268<br>±<br>0.05760 | 0.684 | 24 | 0.7677<br>±<br>0.05717 | 0.6900 | 340.5 | 0.3741 | 57 | 0.6444<br>±<br>0.04464 | 0.581 | 67 | 0.7470<br>±<br>0.04898 | 0.629 | 1682;<br>0.2545 |

**Supplementary Table S8B.** Kruskal–Wallis and Dunn-adjusted comparisons for CeA. Distributional variations across the studied cohorts within the CeA region of the amygdala were evaluated using the Kruskal-Wallis test. All statistical metrics and parameters are detailed in Supplementary S1B for reference.

| Panel | Endpoint | KW p | 6 Veh vs<br>6 VU01<br>rank<br>diff. | adj. p | 6 Veh vs<br>12 Veh<br>rank<br>diff. | adj. p | 6 Veh vs<br>12 VU01<br>rank<br>diff. | adj. p | 6 VU01<br>vs 12<br>Veh<br>rank<br>diff. | adj. p | 6 VU01<br>vs 12<br>VU01<br>rank<br>diff. | adj. p | 12 Veh<br>vs 12<br>VU01<br>rank<br>diff. | adj. p |
| --- | --- | --- | --- | --- | --- | --- | --- | --- | --- | --- | --- | --- | --- | --- |
| A | Number of<br>all Spines | <0.0001 | -4.252 | >0.9999 | -120.5 | <0.0001 | -118.2 | <0.0001 | -116.3 | <0.0001 | -113.9 | <0.0001 | 2.327 | >0.9999 |
| B | Stubby<br>Spines | <0.0001 | 35.35 | 0.0876 | -94.96 | <0.0001 | -92.41 | <0.0001 | -130.3 | <0.0001 | -127.8 | <0.0001 | 2.553 | >0.9999 |
| C | Filamentous<br>Spines | <0.0001 | -97.9 | 0.0011 | -26.84 | >0.9999 | -30.76 | >0.9999 | 71.06 | <0.0001 | 67.14 | <0.0001 | -3.92 | >0.9999 |
| D | Mushroom<br>Spines | <0.0001 | 18.49 | >0.9999 | 64.58 | <0.0001 | 57.9 | 0.0002 | 46.09 | 0.0028 | 39.41 | 0.0164 | -6.68 | >0.9999 |
| E | Spine<br>Density per<br>10 μm | <0.0001 | -39.60 | 0.0401 | -133.8 | <0.0001 | -125.0 | <0.0001 | -94.19 | <0.0001 | -85.37 | <0.0001 | 8.813 | >0.9999 |
| F | Individual<br>Spine Area | <0.0001 | 21.55 | >0.9999 | 841.2 | <0.0001 | 630.2 | <0.0001 | 819.7 | <0.0001 | 608.7 | <0.0001 | -211 | <0.0001 |
| G | Total<br>Dendritic<br>Area | <0.0001 | -10.47 | >0.9999 | -51.97 | 0.0008 | -75.46 | <0.0001 | -41.5 | 0.0077 | -64.99 | <0.0001 | -23.49 | 0.2855 |
| H | Spine<br>Perimeter | <0.0001 | 78.52 | >0.9999 | 636.2 | <0.0001 | 414.2 | <0.0001 | 557.7 | <0.0001 | 335.7 | <0.0001 | -222 | <0.0001 |
| I | Dendrite<br>Major Axis | <0.0001 | 59.76 | >0.9999 | 623.5 | <0.0001 | 428.9 | <0.0001 | 563.8 | <0.0001 | 369.1 | <0.0001 | -194.6 | <0.0001 |
| J | Dendrite<br>Minor Axis | 0.0001 | 3.076 | >0.9999 | 552 | <0.0001 | 359.3 | <0.0001 | 548.9 | <0.0001 | 356.3 | <0.0001 | -192.6 | <0.0001 |
| K | Mushroom<br>Spine Head<br>Width | 0.0003 | -14.31 | >0.9999 | 34.60 | 0.0169 | 22.52 | 0.2672 | 48.92 | 0.0009 | 36.84 | 0.0200 | -12.08 | >0.9999 |
| L | Mushroom<br>Spine Neck<br>Length | 0.2709 | -10.51 | >0.9999 | 13.39 | >0.9999 | 2.526 | >0.9999 | 23.90 | 0.3648 | 13.04 | >0.9999 | -10.86 | >0.9999 |

Supplementary note. n denotes animals; number of observations denotes pooled measurement-level observations. Multiple observations were nested within n = 5 animals per group. Dunn adjustment was applied within each endpoint-specific six-comparison family; no global correction was applied across regions and endpoints.

**Supplementary Table S9A.** Descriptive statistics and within-age Mann–Whitney results for LA. Mann-Whitney U-test statistics, medians and means ± SEM were evaluated for the LA region of the amygdala. All statistical measures and parameters follow the schema of Supplementary Table S1A.

| Panel | Endpoint | Level | 6 Veh<br>n | 6 Veh<br>mean ±<br>SEM | 6 Veh<br>median | 6<br>VU01<br>n | 6 VU01<br>mean ±<br>SEM | 6 VU01<br>median | U 6 mo | p 6 mo | 12<br>Veh n | 12 Veh<br>mean ±<br>SEM | 12 Veh<br>median | 12<br>VU01<br>n | 12<br>VU01<br>mean ±<br>SEM | 12<br>VU01<br>median | U; p 12<br>mo |
| --- | --- | --- | --- | --- | --- | --- | --- | --- | --- | --- | --- | --- | --- | --- | --- | --- | --- |
| A | Number of<br>all Spines | Dendritic<br>segment | 45 | 6.422 ±<br>0.3976 | 6 | 54 | 7.519 ±<br>0.3114 | 8 | 926 | 0.0402 | 75 | 15.2 ±<br>0.3582 | 16 | 75 | 14.85 ±<br>0.3225 | 15 | 2600;<br>0.4236 |
| B | Stubby<br>Spines | Dendritic<br>segment | 45 | 5.933 ±<br>0.3819 | 6 | 54 | 4.463 ±<br>0.264 | 4 | 791 | 0.0024 | 75 | 13.68 ±<br>0.3716 | 14 | 75 | 12.65 ±<br>0.3366 | 12 | 2333;<br>0.0701 |
| C | Filamentous<br>Spines | Dendritic<br>segment | 4 | 0.5 ±<br>0.2887 | 0.5 | 36 | 3.917 ±<br>0.2684 | 4 | 3 | 0.0001 | 75 | 0.88 ±<br>0.1328 | 0 | 75 | 1.12 ±<br>0.1553 | 1 | 2550;<br>0.2925 |
| D | Mushroom<br>Spines | Mushroom<br>spine | 10 | 2.2 ±<br>0.5333 | 1.5 | 17 | 1.647 ±<br>0.2704 | 1 | 68 | 0.3593 | 75 | 0.6533<br>±<br>0.1099 | 0 | 75 | 1.08 ±<br>0.1706 | 1 | 2400;<br>0.0879 |
| E | Spine<br>Density per<br>10 µm | Dendritic<br>segment | 45 | 0.6515<br>± 0.041 | 0.6001 | 55 | 0.7465<br>±<br>0.0314 | 0.7128 | 972 | 0.0662 | 75 | 2.624 ±<br>0.1071 | 2.552 | 75 | 2.34 ±<br>0.0871 | 2.32 | 2361;<br>0.0895 |
| F | Individual<br>Spine Area | Individual<br>spine | 288 | 0.3582<br>±<br>0.0309 | 0.234 | 409 | 0.3179<br>±<br>0.0154 | 0.222 | 58563 | 0.8988 | 1141 | 0.1895<br>± 0.009 | 0.086 | 1116 | 0.2421<br>±<br>0.0133 | 0.1045 | 572448;<br><0.0001 |
| G | Total<br>Dendritic<br>Area | Dendritic<br>segment | 45 | 6.668 ±<br>0.3454 | 6.28 | 55 | 7.717 ±<br>0.2614 | 7.055 | 757 | 0.0007 | 75 | 8.704 ±<br>0.3018 | 8.41 | 75 | 10.25 ±<br>0.4634 | 8.89 | 2255;<br>0.0359 |
| H | Spine<br>Perimeter | Individual<br>spine | 287 | 2.304 ±<br>0.0721 | 1.955 | 409 | 2.156 ±<br>0.0523 | 1.893 | 55040 | 0.162 | 1124 | 1.651 ±<br>0.0299 | 1.385 | 1119 | 1.938 ±<br>0.0479 | 1.481 | 566446;<br><0.0001 |
| I | Dendrite<br>Major Axis | Dendritic<br>segment | 286 | 0.962 ±<br>0.0274 | 0.844 | 439 | 0.9472<br>± 0.04 | 0.788 | 58406 | 0.1127 | 1142 | 0.6842<br>±<br>0.0134 | 0.587 | 1118 | 0.7907<br>±<br>0.0217 | 0.629 | 581774;<br>0.0003 |
| J | Dendrite<br>Minor Axis | Dendritic<br>segment | 286 | 0.3789<br>±<br>0.0165 | 0.311 | 439 | 0.4124<br>±<br>0.0132 | 0.342 | 54722 | 0.0035 | 1142 | 0.2871<br>± 0.007 | 0.202 | 1118 | 0.3533<br>±<br>0.0093 | 0.233 | 557309;<br><0.0001 |
| K | Mushroom<br>Spine Head<br>Width | Mushroom<br>spine | 23 | 1.119 ±<br>0.1184 | 1.027 | 28 | 1.218 ±<br>0.0856 | 1.157 | 256.5 | 0.2188 | 49 | 0.7048<br>±<br>0.0449 | 0.642 | 81 | 0.7837<br>± 0.035 | 0.767 | 1642;<br>0.1003 |
| L | Mushroom<br>Spine Neck<br>Length | Mushroom<br>spine | 23 | 0.9804<br>±<br>0.09614 | 0.880 | 28 | 0.9453<br>±<br>0.06217 | 0.8820 | 321.5 | 0.9963 | 53 | 0.6690<br>±<br>0.04196 | 0.600 | 80 | 0.6924<br>±<br>0.03479 | 0.6330 | 2008;<br>0.6071 |

**Supplementary Table S9B.** Kruskal–Wallis and Dunn-adjusted comparisons for LA. Variance across study groups was observed using the Kruskal-Wallis test. For a detailed index of the statistical measures and parameters, refer to Supplementary Table S1B.

| Panel | Endpoint | KW p | 6 Veh vs<br>6 VU01<br>rank<br>diff. | adj. p | 6 Veh vs<br>12 Veh<br>rank<br>diff. | adj. p | 6 Veh vs<br>12 VU01<br>rank<br>diff. | adj. p | 6 VU01<br>vs 12<br>Veh<br>rank<br>diff. | adj. p | 6 VU01<br>vs 12<br>VU01<br>rank<br>diff. | adj. p | 12 Veh<br>vs 12<br>VU01<br>rank<br>diff. | adj. p |
| --- | --- | --- | --- | --- | --- | --- | --- | --- | --- | --- | --- | --- | --- | --- |
| A | Number of<br>all Spines | <0.0001 | -13.49 | >0.9999 | -129.3 | <0.0001 | -124 | <0.0001 | -115.8 | <0.0001 | -110.5 | <0.0001 | 5.26 | >0.9999 |
| B | Stubby<br>Spines | <0.0001 | 21.37 | 0.844 | -114.9 | <0.0001 | -101.1 | <0.0001 | -136.3 | <0.0001 | -122.5 | <0.0001 | 13.83 | >0.9999 |
| C | Filamentous<br>Spines | <0.0001 | -94.76 | 0.0041 | -10.48 | >0.9999 | -18.77 | >0.9999 | 84.28 | <0.0001 | 76 | <0.0001 | -8.287 | >0.9999 |
| D | Mushroom<br>Spines | <0.0001 | 9.774 | >0.9999 | 57.99 | 0.002 | 44.7 | 0.0337 | 48.22 | 0.0011 | 34.92 | 0.0402 | -13.29 | 0.5372 |
| E | Spine<br>Density per<br>10 μm | <0.0001 | -11.06 | >0.9999 | -136.6 | <0.0001 | -124.1 | <0.0001 | -125.5 | <0.0001 | -113.1 | <0.0001 | 12.47 | >0.9999 |
| F | Individual<br>Spine Area | <0.0001 | -9.66 | >0.9999 | 785.4 | <0.0001 | 630.9 | <0.0001 | 795 | <0.0001 | 640.5 | <0.0001 | -154.5 | 0.0001 |
| G | Total<br>Dendritic<br>Area | <0.0001 | -38.58 | 0.0477 | -61.06 | <0.0001 | -84.02 | <0.0001 | -22.47 | 0.48 | -45.43 | 0.0024 | -22.96 | 0.3111 |
| H | Spine<br>Perimeter | <0.0001 | 56.93 | >0.9999 | 612.3 | <0.0001 | 456.6 | <0.0001 | 555.3 | <0.0001 | 399.7 | <0.0001 | -155.6 | <0.0001 |
| I | Dendrite<br>Major Axis | <0.0001 | 80.03 | >0.9999 | 655.2 | <0.0001 | 515.8 | <0.0001 | 575.2 | <0.0001 | 435.8 | <0.0001 | -139.4 | 0.0007 |
| J | Dendrite<br>Minor Axis | <0.0001 | -168.4 | 0.0608 | 461.6 | <0.0001 | 266 | <0.0001 | 629.9 | <0.0001 | 434.4 | <0.0001 | -195.6 | <0.0001 |
| K | Mushroom<br>Spine Head<br>Width | <0.0001 | -16.20 | >0.9999 | 47.71 | 0.0019 | 33.23 | 0.0436 | 63.91 | <0.0001 | 49.43 | 0.0001 | -14.48 | 0.7608 |
| L | Mushroom<br>Spine Neck<br>Length | <0.0001 | -5.479 | >0.9999 | 40.96 | 0.0124 | 36.23 | 0.0242 | 46.44 | 0.0011 | 41.71 | 0.0022 | -4.733 | >0.9999 |

Supplementary note. n denotes animals; number of observations denotes pooled measurement-level observations. Multiple observations were nested within n = 5 animals per group. Dunn adjustment was applied within each endpoint-specific six-comparison family; no global correction was applied across regions and endpoints.

**Supplementary Table S10A.** Descriptive statistics and within-age Mann–Whitney results for BLA. To characterize variations within the BLA region of the amygdala, Mann-Whitney tests were conducted to report U-test statistics, medians and means ± SEM. All statistical metrics and other parameters are detailed further in Supplementary Table S1A.

| Panel | Endpoint | Level | 6 Veh<br>n | 6 Veh<br>mean ±<br>SEM | 6 Veh<br>median | 6 VU01<br>n | 6 VU01<br>mean ±<br>SEM | 6 VU01<br>median | U 6 mo | p 6 mo | 12<br>Veh n | 12 Veh<br>mean ±<br>SEM | 12 Veh<br>median | 12<br>VU01<br>n | 12<br>VU01<br>mean ±<br>SEM | 12<br>VU01<br>median | U; p 12<br>mo |
| --- | --- | --- | --- | --- | --- | --- | --- | --- | --- | --- | --- | --- | --- | --- | --- | --- | --- |
| A | Number of<br>all Spines | Dendritic<br>segment | 45 | 7.711 ±<br>0.4295 | 8 | 50 | 8.74 ±<br>0.4461 | 8 | 937.5 | 0.1612 | 75 | 14.81 ±<br>0.3357 | 15 | 75 | 14.99 ±<br>0.3732 | 15 | 2689;<br>0.6411 |
| B | Stubby<br>Spines | Dendritic<br>segment | 45 | 7.089 ±<br>0.4413 | 7 | 50 | 5.14 ±<br>0.3692 | 5 | 684 | 0.0008 | 75 | 13 ±<br>0.3995 | 13 | 75 | 12.73 ±<br>0.4147 | 13 | 2684;<br>0.6293 |
| C | Filamentous<br>Spines | Dendritic<br>segment | 10 | 0.2 ±<br>0.1333 | 0 | 50 | 5.14 ±<br>0.3692 | 5 | 1 | <0.0001 | 59 | 1.458 ±<br>0.2373 | 1 | 59 | 1.288 ±<br>0.1889 | 1 | 1704;<br>0.8394 |
| D | Mushroom<br>Spines | Mushroom<br>spine | 9 | 2 ±<br>0.4082 | 2 | 14 | 1.714 ±<br>0.2442 | 1 | 55 | 0.602 | 52 | 0.9423 ±<br>0.1611 | 0.5 | 57 | 1.649 ±<br>0.2497 | 1 | 1135;<br>0.0275 |
| E | Spine<br>Density per<br>10 μm | Dendritic<br>segment | 45 | 0.7757 ±<br>0.0435 | 0.783 | 55 | 0.9153 ±<br>0.0455 | 0.8763 | 930 | 0.0328 | 75 | 2.496 ±<br>0.1079 | 2.317 | 75 | 2.43 ±<br>0.0928 | 2.339 | 2771;<br>0.8779 |
| F | Individual<br>Spine Area | Individual<br>spine | 347 | 0.2937 ±<br>0.0232 | 0.194 | 504 | 0.2609 ±<br>0.0107 | 0.188 | 85135 | 0.5123 | 1112 | 0.2126 ±<br>0.0127 | 0.092 | 1125 | 0.2204 ±<br>0.0123 | 0.093 | 603524;<br>0.1502 |
| G | Total<br>Dendritic<br>Area | Dendritic<br>segment | 45 | 7.143 ±<br>0.406 | 6.166 | 55 | 7.477 ±<br>0.2659 | 7.146 | 1009 | 0.1133 | 75 | 9.487 ±<br>0.4101 | 8.586 | 75 | 9.769 ±<br>0.4005 | 9.108 | 2645;<br>0.5307 |
| H | Spine<br>Perimeter | Individual<br>spine | 348 | 2.153 ±<br>0.1037 | 1.723 | 504 | 1.866 ±<br>0.0395 | 1.666 | 79637 | 0.0225 | 1112 | 1.842 ±<br>0.0492 | 1.431 | 1125 | 1.848 ±<br>0.0415 | 1.431 | 614410;<br>0.4678 |
| I | Dendrite<br>Major Axis | Dendritic<br>segment | 350 | 0.9533 ±<br>0.0572 | 0.743 | 506 | 0.7946 ±<br>0.0305 | 0.684 | 77793 | 0.0025 | 1112 | 0.7463 ±<br>0.0192 | 0.6125 | 1125 | 0.7404 ±<br>0.0154 | 0.6 | 622148;<br>0.8263 |
| J | Dendrite<br>Minor Axis | Dendritic<br>segment | 350 | 0.3618 ±<br>0.014 | 0.298 | 506 | 0.3738 ±<br>0.0107 | 0.3195 | 81780 | 0.057 | 1112 | 0.3166 ±<br>0.0084 | 0.212 | 1125 | 0.3305 ±<br>0.0083 | 0.22 | 600306;<br>0.0991 |
| K | Mushroom<br>Spine Head<br>Width | Mushroom<br>spine | 18 | 1.260 ±<br>0.1700 | 1.081 | 24 | 1.023 ±<br>0.06318 | 0.974 | 177.5 | 0.3345 | 50 | 0.8653 ±<br>0.05940 | 0.837 | 94 | 0.7404 ±<br>0.03666 | 0.6445 | 1930;<br>0.0778 |
| L | Mushroom<br>Spine Neck<br>Length | Mushroom<br>spine | 18 | 1.017 ±<br>0.1048 | 0.9135 | 24 | 0.8859 ±<br>0.07127 | 0.884 | 179 | 0.3538 | 49 | 0.7051 ±<br>0.04727 | 0.642 | 97 | 0.6469 ±<br>0.03167 | 0.590 | 2110;<br>0.2709 |

**Supplementary Table S10B.**
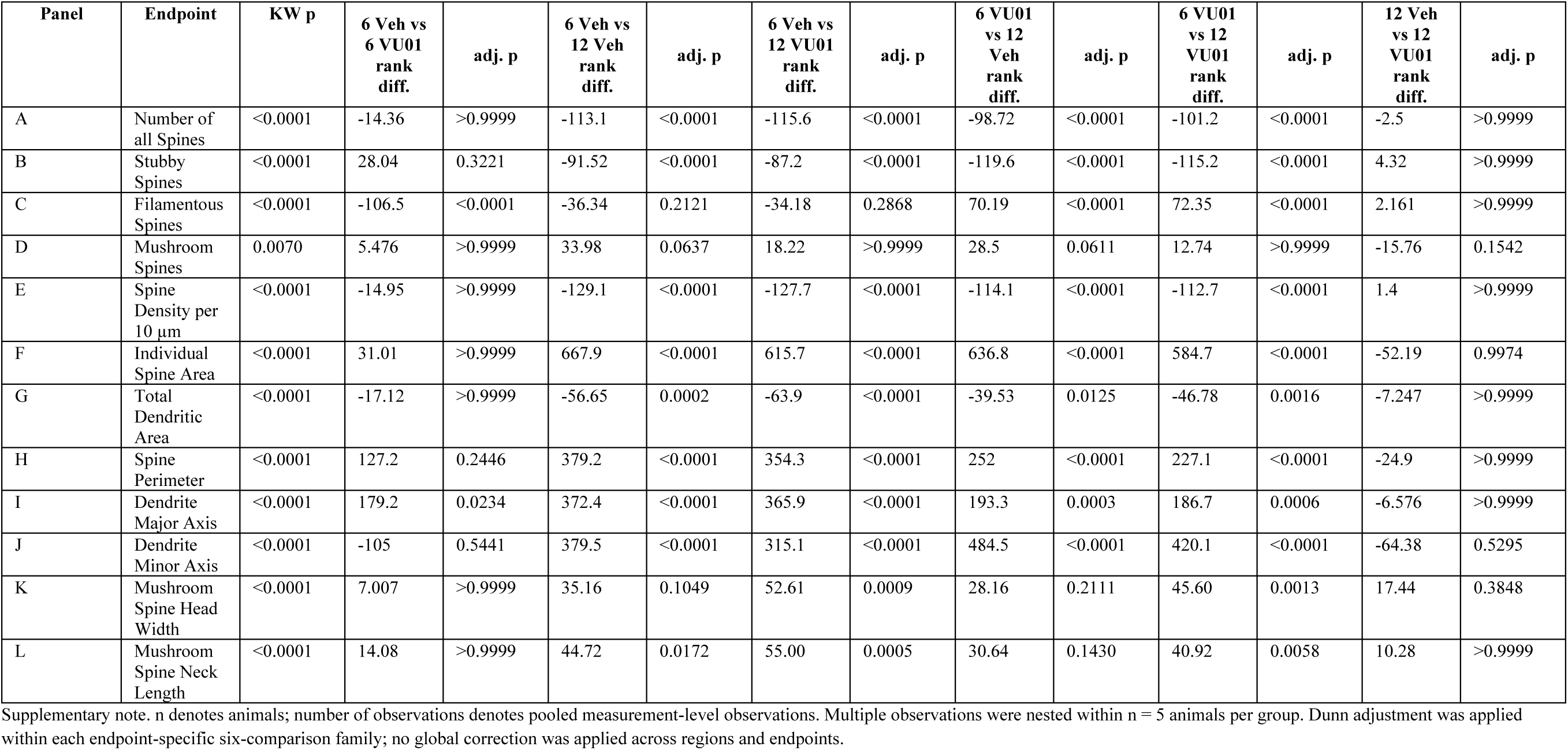
Kruskal–Wallis and Dunn-adjusted comparisons for BLA. For assessment of multi-group variation, Kruskal-Wallis test was utilized. Associated statistical measures and parameters follow the schema of Supplementary Table S1B which can be referenced.

| Panel | Endpoint | KW p | 6 Veh vs<br>6 VU01<br>rank<br>diff. | adj. p | 6 Veh vs<br>12 Veh<br>rank<br>diff. | adj. p | 6 Veh vs<br>12 VU01<br>rank<br>diff. | adj. p | 6 VU01<br>vs 12<br>Veh<br>rank<br>diff. | adj. p | 6 VU01<br>vs 12<br>VU01<br>rank<br>diff. | adj. p | 12 Veh<br>vs 12<br>VU01<br>rank<br>diff. | adj. p |
| --- | --- | --- | --- | --- | --- | --- | --- | --- | --- | --- | --- | --- | --- | --- |
| A | Number of<br>all Spines | <0.0001 | -14.36 | >0.9999 | -113.1 | <0.0001 | -115.6 | <0.0001 | -98.72 | <0.0001 | -101.2 | <0.0001 | -2.5 | >0.9999 |
| B | Stubby<br>Spines | <0.0001 | 28.04 | 0.3221 | -91.52 | <0.0001 | -87.2 | <0.0001 | -119.6 | <0.0001 | -115.2 | <0.0001 | 4.32 | >0.9999 |
| C | Filamentous<br>Spines | <0.0001 | -106.5 | <0.0001 | -36.34 | 0.2121 | -34.18 | 0.2868 | 70.19 | <0.0001 | 72.35 | <0.0001 | 2.161 | >0.9999 |
| D | Mushroom<br>Spines | 0.0070 | 5.476 | >0.9999 | 33.98 | 0.0637 | 18.22 | >0.9999 | 28.5 | 0.0611 | 12.74 | >0.9999 | -15.76 | 0.1542 |
| E | Spine<br>Density per<br>10 μm | <0.0001 | -14.95 | >0.9999 | -129.1 | <0.0001 | -127.7 | <0.0001 | -114.1 | <0.0001 | -112.7 | <0.0001 | 1.4 | >0.9999 |
| F | Individual<br>Spine Area | <0.0001 | 31.01 | >0.9999 | 667.9 | <0.0001 | 615.7 | <0.0001 | 636.8 | <0.0001 | 584.7 | <0.0001 | -52.19 | 0.9974 |
| G | Total<br>Dendritic<br>Area | <0.0001 | -17.12 | >0.9999 | -56.65 | 0.0002 | -63.9 | <0.0001 | -39.53 | 0.0125 | -46.78 | 0.0016 | -7.247 | >0.9999 |
| H | Spine<br>Perimeter | <0.0001 | 127.2 | 0.2446 | 379.2 | <0.0001 | 354.3 | <0.0001 | 252 | <0.0001 | 227.1 | <0.0001 | -24.9 | >0.9999 |
| I | Dendrite<br>Major Axis | <0.0001 | 179.2 | 0.0234 | 372.4 | <0.0001 | 365.9 | <0.0001 | 193.3 | 0.0003 | 186.7 | 0.0006 | -6.576 | >0.9999 |
| J | Dendrite<br>Minor Axis | <0.0001 | -105 | 0.5441 | 379.5 | <0.0001 | 315.1 | <0.0001 | 484.5 | <0.0001 | 420.1 | <0.0001 | -64.38 | 0.5295 |
| K | Mushroom<br>Spine Head<br>Width | <0.0001 | 7.007 | >0.9999 | 35.16 | 0.1049 | 52.61 | 0.0009 | 28.16 | 0.2111 | 45.60 | 0.0013 | 17.44 | 0.3848 |
| L | Mushroom<br>Spine Neck<br>Length | <0.0001 | 14.08 | >0.9999 | 44.72 | 0.0172 | 55.00 | 0.0005 | 30.64 | 0.1430 | 40.92 | 0.0058 | 10.28 | >0.9999 |

Supplementary note. n denotes animals; number of observations denotes pooled measurement-level observations. Multiple observations were nested within n = 5 animals per group. Dunn adjustment was applied within each endpoint-specific six-comparison family; no global correction was applied across regions and endpoints.

## Notes

### Competing Interest Statement

The authors have declared no competing interest.

